# Recognition of specific PIP2-subtype composition triggers the allosteric control mechanism for selective membrane targeting of cargo loading and release functions of the intracellular sterol transporter StarD4

**DOI:** 10.1101/2024.03.26.586881

**Authors:** Hengyi Xie, Harel Weinstein

**Affiliations:** Department of Physiology and Biophysics, Weill Cornell Medicine, New York, NY, 10065, USA; Institute for Computational Biomedicine, Weill Cornell Medicine, New York, NY 10065, USA

**Keywords:** Molecular Dynamics (MD) simulations, cholesterol traffic and distribution in cells, conformational changes coupled to modes of cholesterol (CHL) binding, membrane embedding of StarD4, recognition of membranes containing specific PIP2 subtypes, allosteric network coordination between the CHL binding site, the cargo release gates and the binding of specific PIP2 subtypes, machine learning analysis of MD trajectories, detection of rare events in MD trajectories, non-negative matrix factorization algorithm, N-body Information Theory analysis of MD trajectories, allosteric channels supporting CHL uptake and release on membrane with different composition

## Abstract

StarD4 is an intracellular cholesterol trafficking protein that facilitates the crucial non-vesicular sterol transport between the plasma membrane and the endoplasmic reticulum. It targets both sterol donor and acceptor membranes via interactions with anionic lipids. Experiments have illuminated the kinetics of this sterol transfer and shown it to be modulated by specific phosphatidylinositol phosphates (PIPs) on the target membrane. The distinct subtype distribution of PIPs in the membranes of cellular organelles serves as a guide to direct StarD4 to recognized cell components. However, little is known about the molecular mechanism of the recognition of the PIP2 subtype by StarD4, and how this affects the direction and kinetics of cholesterol transport, as the reaction pathways of the cholesterol uptake and release processes in StarD4 have never been observed. Here, we investigated 1)-how StarD4 transports a cholesterol from/to membranes; 2)-how StarD4 recognizes PIP2-subtypes in membranes; and 3)-how the PIP2-subtype recognition impacts cholesterol transport kinetics, using extensive molecular dynamics (MD) sampling with advanced machine learning and information theory methods for trajectory analysis. The findings revealed function-related allosteric dynamics of StarD4, connecting the identified PIP2-subtype-specific conformational states to the cholesterol binding modes in the pocket, which steers the dynamics of the gates towards conformations that support either cholesterol release or uptake. This reveals the crucial role of PIP2 subtypes in shaping functional StarD4 motifs responsible for organelle selectivity of the cholesterol trafficking, providing fundamental insights into cellular cholesterol regulation.

## Introduction

Interest in the process of dynamic trafficking of cholesterol (CHL) among diverse locations and organelles in the cell has brought to light the role of cellular proteins containing **<u>st</u>**eroidogenic **<u>a</u>**cute **<u>r</u>**egulator-related lipid transfer (START) domains (StarD), such as StarD4 (1–7). The StarD4 protein specializes in binding and transport sterols (8,9), contributing approximately 33% of the non-vesicular sterol transport between the primary cholesterol reservoirs — the plasma membrane (PM) and the endocytic recycling compartment (ERC), and the endoplasmic reticulum (ER) — which governs sterol homeostasis maintenance, cholesterol concentration sensing and regulation (3,10,11).

StarD4 contains a single soluble START domain featured by an α/β helix-grip fold that forms an internal hydrophobic cavity, which is essential for cargo lipid recognition and binding (6,7). Unlike the proteins in the StarD1 subfamily that localize to specific membrane contact sites via dedicated membrane-binding domains, StarD4, with its single soluble START domain, relies on interactions with anionic lipids such as phosphatidylserine (PS) and phosphatidylinositol phosphates (PIPs) to target sterol donor and acceptor membranes (9,12). Recent investigations have revealed the modulation of sterol transfer kinetics by StarD4 via specific subtypes of anionic lipids (12). The sterol transportation by StarD4 on membranes containing phosphatidylinositol bisphosphate (PIP2) is accelerated compared to those containing PS as the only anionic lipid. More interestingly, StarD4 preferentially extracts sterol from donor membranes resembling plasma membranes that contain phosphatidylinositol 4,5-bisphosphate (PI(4,5)P_2_), as opposed to membranes containing phosphatidylinositol 3,5-bisphosphate (PI(3,5)P_2_). Given that distinct PIPs are selectively enriched on cellular organelles (13), these negatively charged lipid molecules can act as organelle-specific fingerprints, guiding soluble StarD4 to specific membrane compartments and modulating its sterol transport kinetics.

Previous studies have attempted to explain StarD4’s anionic lipid specificity using molecular dynamics (MD) simulations (12,14). The difference in sterol transport kinetics between PS and PIP2-containing membranes is attributed to the strength of the membrane-StarD4 interactions. Both studies suggest that StarD4 exhibits more superficial anchoring to membranes containing PS as the sole anionic lipid, leading to a broader range of orientations in different trajectories (12,14). Some of these orientations are less productive for cholesterol transport, as they lack substantial interaction between StarD4’s sterol gate and the membrane. This weak membrane anchoring may be due to the lower charge density and valency of PS compared to PIP2. However, the preference of StarD4 for PI(4,5)P_2_-containing membranes over PI(3,5)P_2_ cannot be explained by electrostatics, as both PIP_2_ subtypes share the same charge valency. It has been proposed that the orientations of StarD4 on PI(3,5)P_2_- and PI(4,5)P_2_-containing membranes exhibit subtle differences (12), but the impact of these differences on cholesterol transport remains unclear. The hypothesis of a “productive orientation” has not been confirmed, as the cholesterol uptake and release pathway from StarD4 has yet to be directly observed. In this study, to investigate the regulatory mechanisms governing sterol transport by StarD4, we conducted extensive millisecond-level molecular dynamics (MD) simulations to characterize the CHL uptake and release pathways of StarD4 on membranes containing PI(4,5)P_2_, PI(3,5)P_2_ and PS.

Although the complete cholesterol transportation pathway between StarD4 and membrane has not been observed, previous studies have revealed the conformation of the sterol-StarD4 complex and the conformational changes required for cholesterol translocation within the hydrophobic pocket of StarD4. As the structure of a START domain in complex with a sterol remains unresolved, these investigations rely primarily on docking and MD simulations (14–22). Initial docking results suggest that cholesterol embeds deeply in the hydrophobic pocket, with its sterol hydroxyl group interacting with S136 and S147 (termed “the Ser-binding mode”) (**Fig. 1A**), but in recent long MD simulations of sterol-StarD4 complex (14,15) show that CHL moves to a new position, where it inserts roughly two-thirds into the pocket near the gate, and its hydroxyl group interacts with W171 in proximity to Y117 and R92 (termed “Trp-binding mode”) (**Fig. 1B**). The dynamic repositioning of CHL in the binding pocket is accompanied by a series of StarD4 conformational changes, including the destabilization of the Ω1-H4 front gate with the H4 move leftwards (**Fig. 1C,D**) and the opening of the β8-Ω4 back corridor (**Fig. 1E,F**), both critical for the membrane interaction of StarD4 (12,14,15). From these sterol-StarD4-membrane complexes, the downstream release/uptake processes and the role of PIP2 subtype remained unobserved, prompting further investigation.

**Figure 1:**
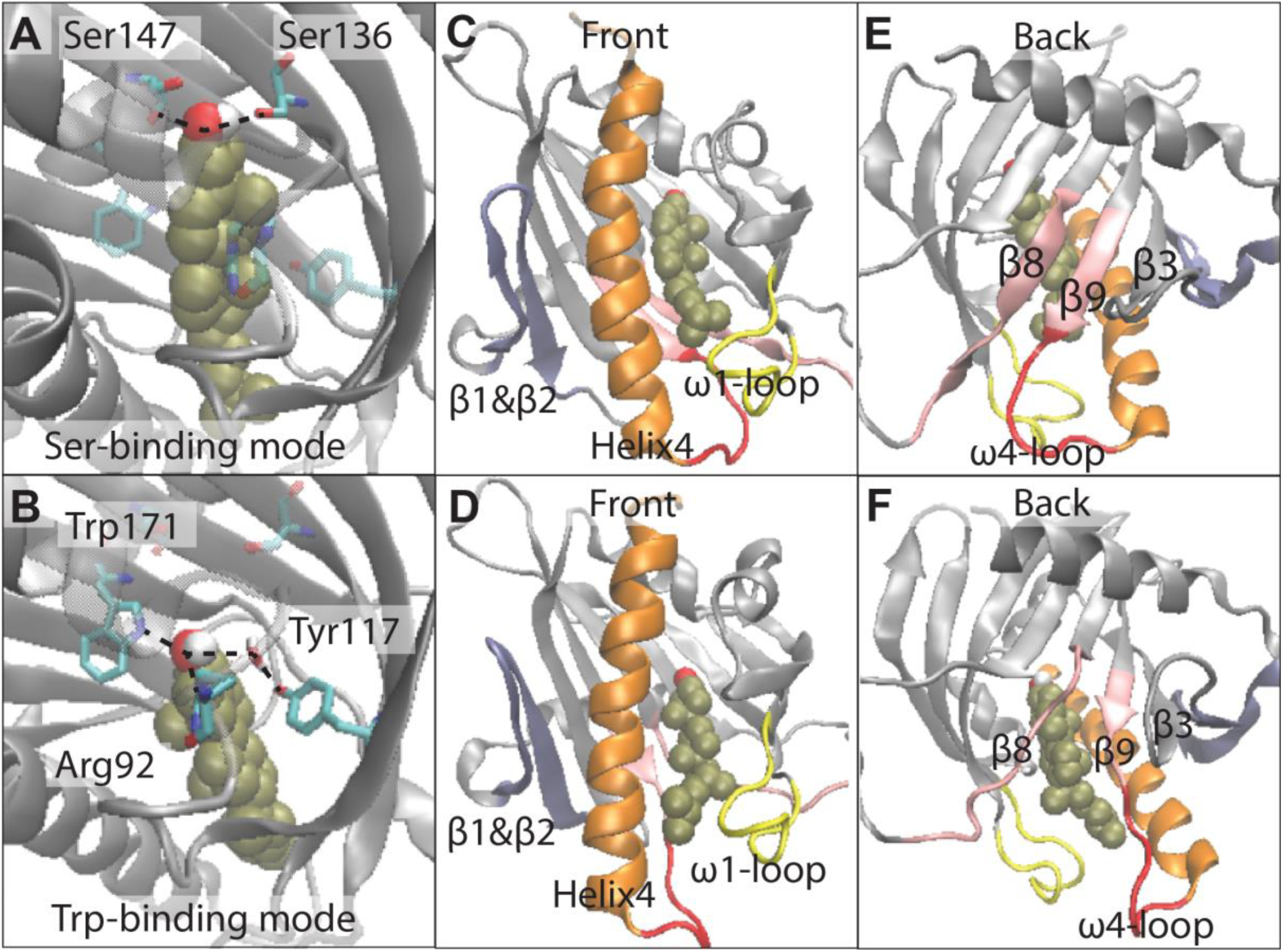
Two modes of cholesterol binding to StarD4 in aqueous solution identified in the long simulation. The two binding modes of CHL in the *holo*-StarD4 complex: **(A)** the “Ser-binding mode”; **(B)** the “Trp-binding mode”. StarD4 is rendered in gray. The residues involved in the binding modes (R92, S136, S147,Y117,W171) are rendered in cylinders, with carbons colored in cyan, nitrogen in blue and oxygen in red (hydrogens are not shown). The CHL (in tan) is shown in VDW rendering, with the hydroxyl group in red and white. **(C-F)** The panels depict conformational changes of StarD4 that occur concurrently (see REF (15)) with the translocation of CHL from one binding mode to the other. The structural segments identified in color are involved in the major changes that include the destabilization of the Ω1-H4 front gate and the opening of the back corridor between β8 and β9 Ω4-loop. Helix4 is colored in orange, the Ω1 in yellow, β1&2 in blue, Ω4 in red, and β8&9 in pink. Note that in **(C),** the front gate between Ω1 and H4 is closed, but as the CHL moves towards the gate and H4 moves towards β1, the gate is destabilized and starts towards opening **(D)**. Similarly, the β8-Ω4 back corridor is closed in **(E)** but opened in **(F)** as β9 and Ω4-loop moved towards β3.

To this end, we employed in the present study a battery of extensive molecular dynamics (MD) simulations to collect the data needed to discover and quantify the key stages of the StarD4 cholesterol trafficking between the membranes it targets. As shown in Figs. 2 and 14, the total simulation time in the progression of sampling stages exceeds 2.5 milliseconds. These include the exploration of molecular mechanisms that determine the selectivity and processes of CHL loading from donor membranes and its release to the acceptor membranes. From the analysis of the MD trajectories, we identified the key conformational changes in StarD4 that are induced by subtype-specific binding of anionic lipids determining the molecular mechanism of cholesterol transport. By employing advanced machine learning (ML) and information theory (IT) analyses, we established allosteric connections underlie the conformational events in this mechanism. This mechanistic understanding reveals how StarD4 recognizes specifically the different PIP2 subtypes and how this recognition modulates the kinetics of sterol transport.

**Figure 2.**
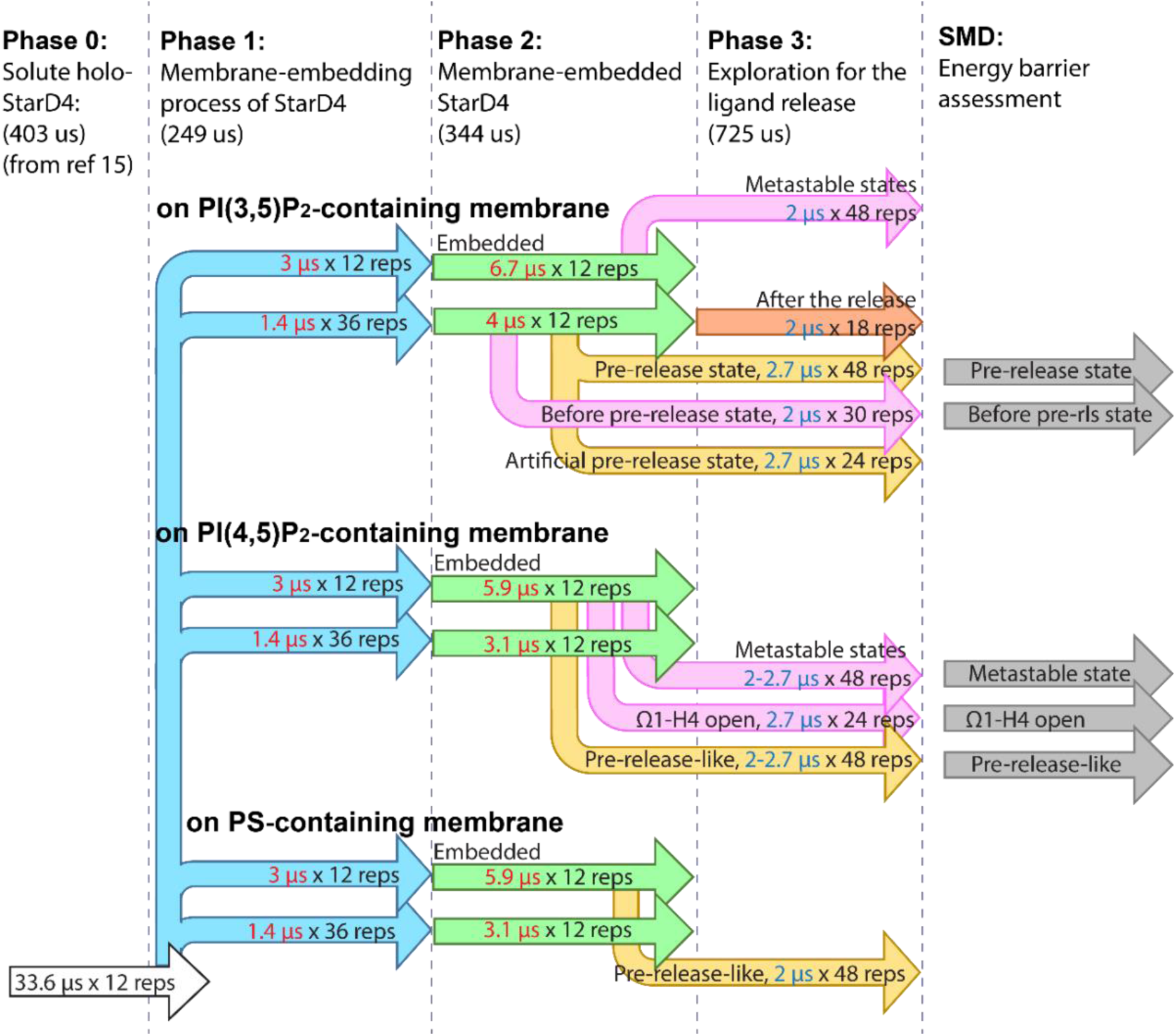
Schematic representation of the atomistic MD simulations of *holo*-StarD4 embedded in the three different membrane models. The labeled components of this schematic plot identify the initial conformations, the simulation time and the number of replicas in each set of simulation contained in Phases 1-3.

Thus, we show that the *<u>release of CHL</u>* occurs in a PIP2-subtype specific manner. The preparatory conformational changes leading to cholesterol release were identified from the application of a recently described machine learning method, Rare Event Detection (RED) (23). These preparatory pre-release conformational changes were found to involve dynamic rearrangements of motifs along an allosteric pathway that channels the information from the PIP2-binding motif to the cholesterol-binding site of StarD4. This allosteric route was quantified with the application of N-body Information Theory (NbIT) analysis (24,25). Employing a Deep Neural Network (DNN) in a separate analysis of the trajectories we found that this allosteric route is a pivotal determinant in distinguishing between StarD4 conformations that interact with PI(3,5)P_2_ vs PI(4,5)P_2_. Remarkably, the PIP2-subtype specific conformational changes of StarD4 are strongly associated with the functional motifs which are responsible for cholesterol release, which reveals the basis for the mechanistic association between the content of the specific PIP2 in the targeted membrane, and the selective release of CHL.

We addressed the *<u>CHL extraction process</u>* from PI(4,5)P_2_-containing membranes into *apo-* StarD4 by exploring with an adaptive sampling protocol of MD simulations and quantified the sequence of events leading from the positioning of membrane cholesterol at the pocket entrance of *apo-*StarD4, to the spontaneous interactions of the hydroxyl group with hydrophilic residues lining the CHL binding pocket. In this process, *apo-*StarD4 visits a conformational space that differs from the one sampled by the *holo*-StarD4 we identified as pre-release and CHL-release states. In contrast, the parallel exploration with extensive simulations of the *apo-*StarD4 interacting with PI(3,5)P_2_-containing membranes (see Fig. 14), does not produce any cholesterol uptake events. We show that when StarD4 interacts with PI(3,5)P_2_-containing membrane, it prefers the conformational states that facilitate cholesterol release, and the only events of cholesterol positioning occur only when StarD4 adopts a pre-release conformational state that precludes the interactions with the residues of the binding pocket seen on the PI(4,5)P_2_-containing membranes.

Collectively, these results provide detailed mechanistic insights regarding the molecular basis for organelle selectivity by StarD4 based on PIP2 subtype recognition. The differences in the modes of interaction with the different subtypes lead to conformational preferences that are propagated through an identified allosteric network from the membrane interaction sites to the sterol ligand binding site, to reach the gate of the hydrophobic pocket and the corridor (see Fig. 1). Consequently, the initial PIP2 subtype recognition regulates the kinetics of the cholesterol trafficking function of StarD4.

## Results

### The process of cholesterol release from *holo*-StarD4 protein

The dynamics of this process were recorded from MD simulations of StarD4 constructs with CHL in the binding site (see Methods and ref (12)) embedded in 3 different membrane models, each containing only one type of negatively charged lipid: PI(3,5)P_2_, or PI(4,5)P_2_, or PS. Notably, the simulation design did not aim to replicate the complexities of different organelles; rather, it aimed to faithfully represent the experimental conditions in the research on membrane vesicles where PIP2-subtype-dependent functional dynamics of were observed (12).

Aiming to understand the sensitivity to PIP2 subtype, the simulation process of each of these *holo*-StarD4-membrane systems consisted of three phases, as depicted in **Figure 2**. After equilibration of membrane lipid rearrangement and StarD4 orientation under Mean-field-model simulations (see Methods), spontaneous StarD4 membrane-embedding processes were sampled by MD simulations in Phase 1 (**Fig 2**). In Phase 2 the membrane-embedded CHL-bound StarD4 system was simulated in 24 independent replica runs of StarD4 embedded in each of the three membrane systems (for a total of 344 microseconds, **Fig 2** Phase 2). During this phase we observed the spontaneous CHL release event illustrated below for Release Trajectory 1 (RT1). In Phase 3, an expanded adaptive sampling protocol (see Methods) was carried out in 312 replicas (for a total of 725 microseconds, **Fig 2** Phase 3) in order to probe whether the CHL release process observed in RT1 is reproduced across different membrane systems, or if alternative release pathways are observed. These simulations yielded four trajectories (RT2-5) that shared the same pre-release characteristics as RT1, and trajectories RT6 and RT7, where CHL release occurred from a distinct conformational state.

The spontaneous releases of the CHL cargo from the StarD4 binding site into the membrane observed in this series of simulations were analyzed to learn the sequence of mechanistic events occurring along the CHL release pathways. This comprehensive analysis, and the mechanistic details it produces, are illustrated below for trajectory RT1 sampled in Phase 2.

### Cholesterol release steps in the RT1 trajectory

The starting structure for the RT1 simulation of the *holo*-StarD4 embedded in PI(3,5)P_2_-containing membranes according to the protocol described in **Fig 2 below,** is the one described in our previous publication (15) as the embedding conformation illustrated in **Figure 1**. The CHL is in its Trp-binding-site, the Ω1-H4 front gate (between the β5-β6-loop and the N-terminal of H4) is slightly open (**Fig 1D**), and the β8-Ω4 back corridor (between the N-terminal of β8 and the β9-H4-loop) is widely open (**Fig 1F**).

In the transition to the CHL release in this trajectory we observed a metastable state that appeared prior to the release of the CHL cargo, with the characteristics quantified in **Figure 3 and Suppl. Fig 1**. In this state, StarD4 adopted a distinct positional orientation relative to the membrane in which the tilting angle between the axis of H4 and the protein axis (defined in **Suppl. Fig 1A)** is increased to an orientation angle >40°, leaning in the direction of the hydrophobic pocket.

**Figure 3.**
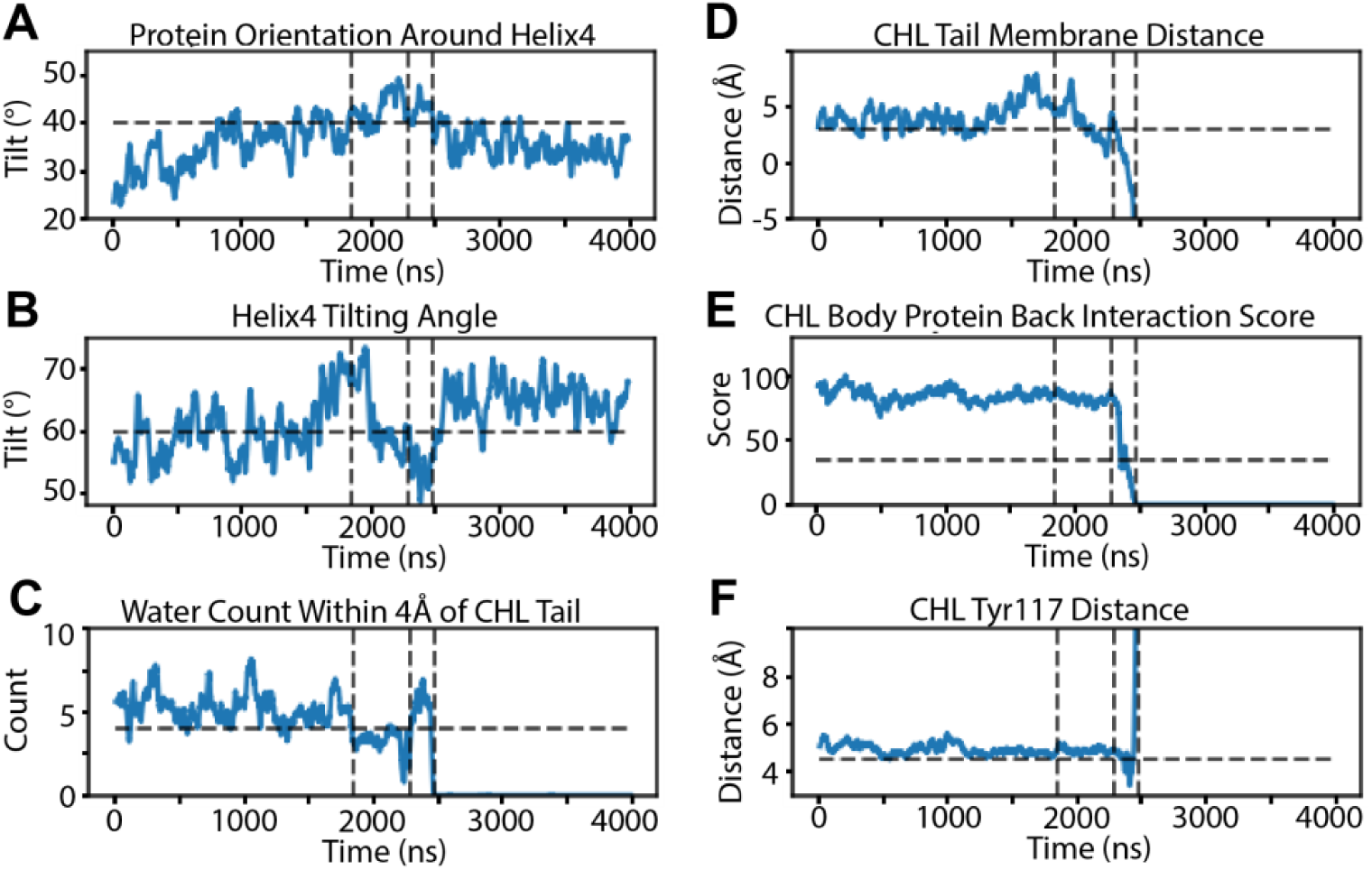
Formation of the pre-release state in the RT1 trajectory of the holo-StarD4 embedded in PI(3,5)P_2_-containing membrane, in which StarD4 adopts a specific orientation and a molecular conformation that promotes the release of its cholesterol cargo. **Panels (A-F)** show the evolution of the structural characteristics of the pre-release state reflected in the measurements of: **(A)** The axial rotation of StarD4 around its H4; **(B)** The tilting angle of H4 in relation to membrane; **(C)** The number of water molecules interacting with the ligand cholesterol; **(D)** The height of cholesterol relative to the membrane; **(E)** The interaction of the CHL hydrocarbon ring with residues of β8&β9; **(F)** The distance between cholesterol and its Y117 binding site. The detailed definitions of the variables are given in the Supplementary Materials, Section 1. The first vertical dashed line marks the entrance into the pre-release; the second vertical dashed line marks the penetration of water; the third vertical dashed line marks the rapid release of cholesterol. The auxiliary horizontal dashed lines indicate the values of the characteristic parameters of the pre-release state to emphasize the transitions.

The evolution of the characteristics is quantified below in **Fig 3A&B** where the occurrence of this “pre-release state” is identified by the vertical dashed lines marking its start and its end when the CHL release phase of the simulation begins. The role of the changes in the facilitation of the subsequent release phase is indicated in **Fig 3C –** which shows that in this time frame the CHL tail is surrounded by a hydrophobic environment (in which the water count is reduced) – and **Fig 3D** showing that this brings the CHL tail closer to the hydrophobic core of the membrane. This allows the cholesterol tail to engage in direct interaction with the membrane through the opened Ω1-H4 gate. Subsequently, the CHL body (hydrocarbon rings) is positioned closer to the Ω1-H4 gate at the front of StarD4, and has weaker hydrophobic contacts with residues N166, C169, I189, T191 on β8 and β9 at the back of StarD4 (**Fig. 3E**), which permits water molecules to enter the created space and disrupt the interaction between cholesterol and StarD4 in this region (**Fig. 3C**). The rapid distancing of the CHL from its initial binding site at the end of this time interval, as seen in **Fig 3F**, indicates that the conditions prevailing in this time interval facilitate the downward repositioning of the CHL to accomplish its release to the membrane.

### After the full release of the CHL

The distinct orientation of StarD4 on the membrane changes rapidly immediately as shown in **Fig 3A&E** where the CHL release time point is marked by the third dashed line marked in all the panels of **Fig. 3**. This indicates that the “maximal leaning towards the membrane” conformation of the StarD4 protein is a “release-enabling” configuration. We note that this particular configuration is rare in the simulations of *holo*-StarD4 on PI(3,5)P_2_-containing membranes, is even less frequently observed when the membranes contain only PI(4,5)P_2_, and never when PS is the anionic lipid as shown in (**Fig 4A-C**). It is also not observed at in our long simulations of *apo*-StarD4, as shown in **Fig 4D-F**.

**Figure 4.**
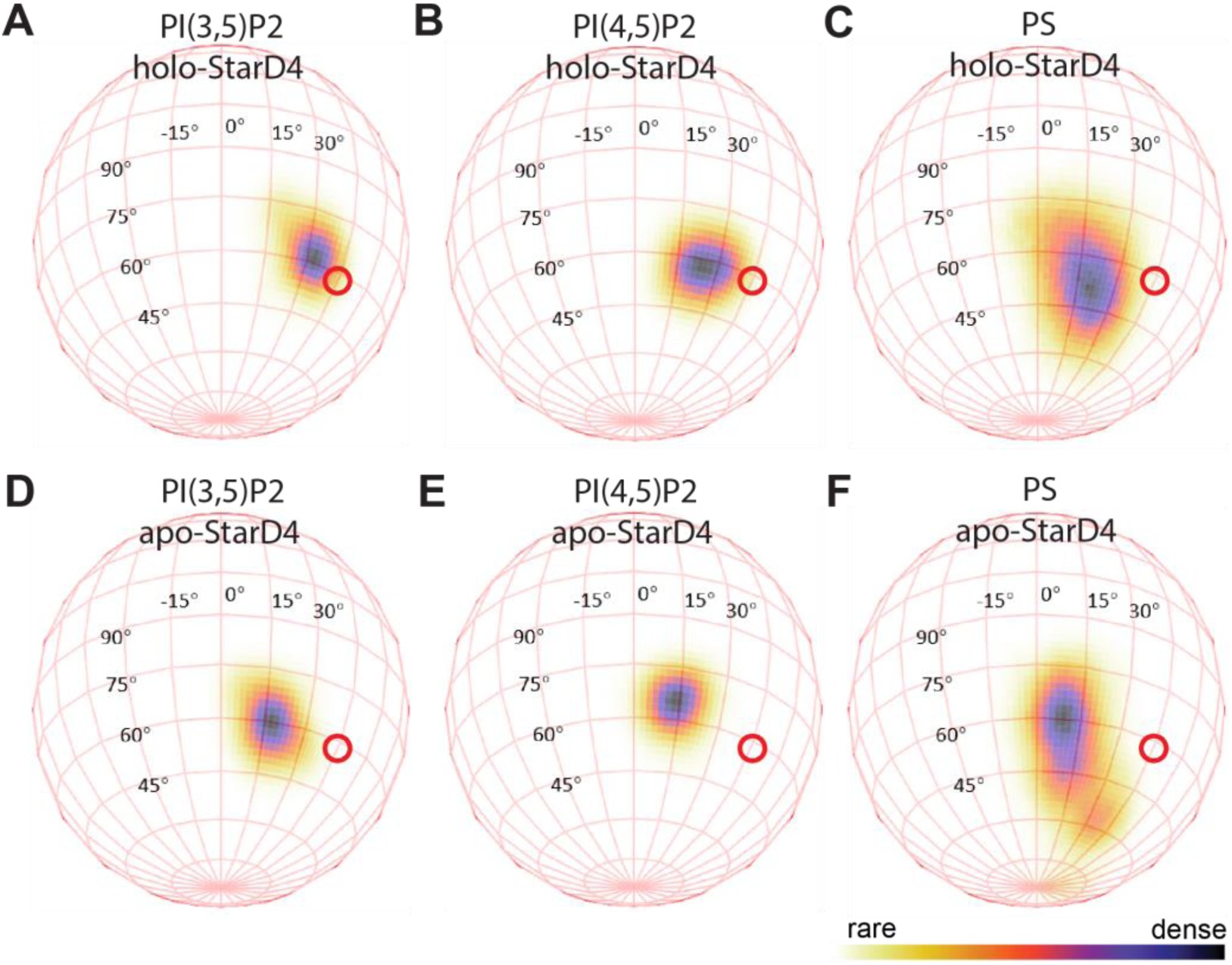
The probability distribution of the of StarD4 orientation in relation to the membrane. In this spherical coordinate system, the vertical axis is the tilting angle of H4, and the horizontal axis measures the axial rotation of protein around H4, as defined in Suppl. Fig. 1. Shown in **A-C** are the orientation distribution of *holo*-StarD4 on PI(3,5)P_2_-, PI(4,5)P_2_- and PS-containing membrane, respectively. Panels **D-F** show the orientation distribution of *apo*-StarD4 on the three types of membranes. The red circle indicates the region that is shared in the spontaneous release events occurring from the pre-release state.

### The temporal sequence of conformational events connecting the initial state of the simulated holo-StarD4 systems to the CHL-release state

To reveal this aspect of the mechanism we applied as described in Methods the Rare Events Detection method (RED) defined in (15,23). Briefly, the RED analysis of the temporal evolution uses the data of residue-residue contacts and of the PIP2-residue-pair interaction in each frame of the trajectory. The machine learning (ML) algorithm then identifies collective motions of the structural motifs, and the PIP2-interacting motifs, in each metastable state encoded in the entire trajectory (15,23). **Fig 5** shows the structural transition components identified by RED in the dynamics of *holo*-StarD4 in the RT1 trajectory (see Methods for the definition of these representations of the structural motifs that dominate the conformation of the membrane embedded StarD4 system during a particular segment of the trajectory). Panel A shows the normalized temporal weight of each component along the RT1 trajectory time. The color code for the components is shown on the right. The weights for all the components are shown at each time point (x-axis) in a representation that stacks their values along the y-axis. For example, at time=2472ns (3^rd^ dashed line), the weight of component 1 (comp 1, in green) is 0.11, comp 3 (in blue) is 0.83, comp 4 (light green) is 0.06, and those of components 0, 2, and 5, are 0.

**Figure 5.**
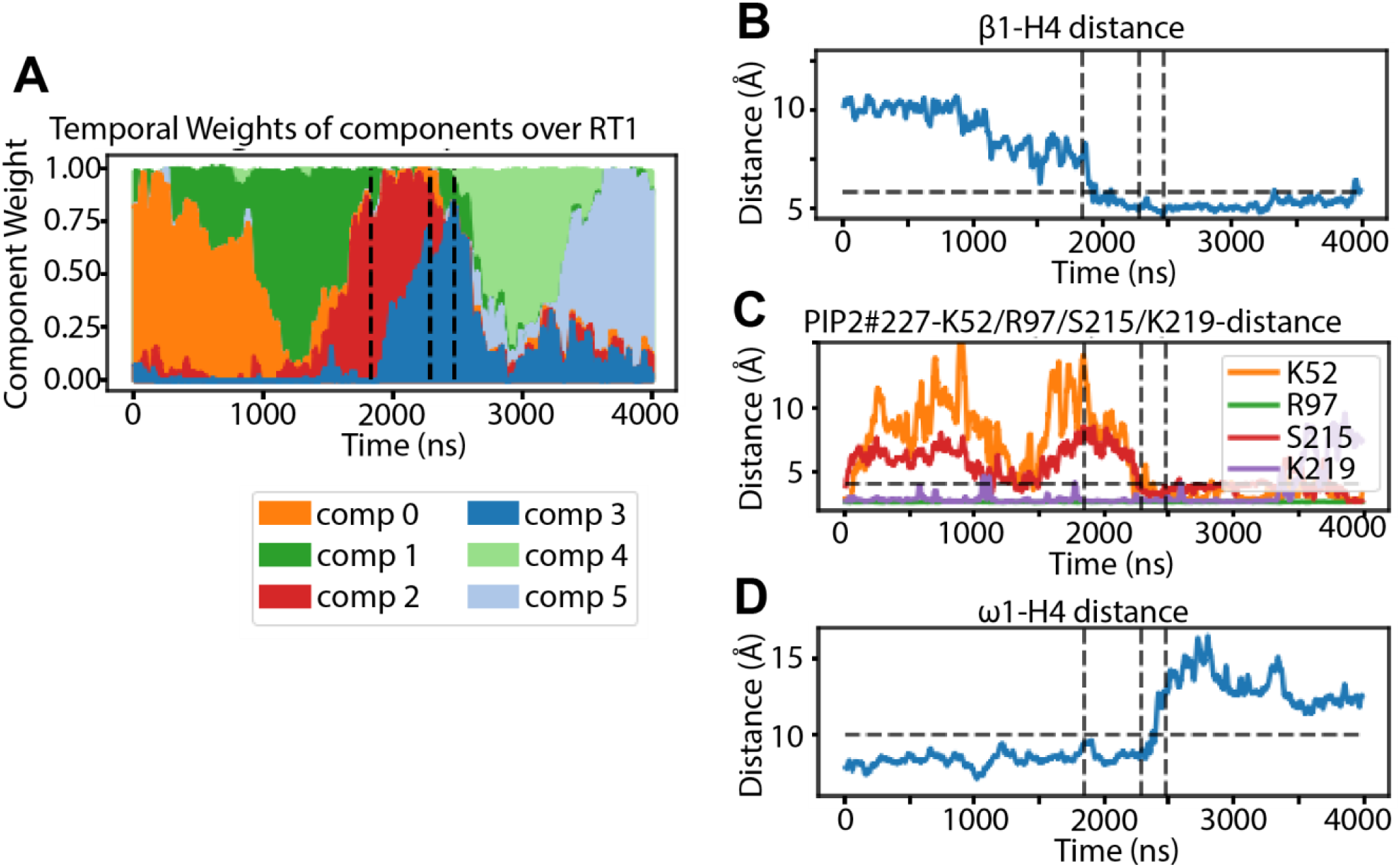
The Rare Event Detection analysis revealed the structural elements that determines the pre-release conformational state. **(A)** The RED-detected normalized temporal weight of the various components along the RT1 trajectory time (see description of the RED protocol in REF (23) and details in Methods). The color code identifies the RED-components as listed on the bottom. **(B-D)** The time-evolution of the structural elements highlighted by the structure-differentiating contact pairs (SDCPs) obtained from the RED analysis, including: **(B)** The interaction of β1-H4; **(C)** the binding of a PIP_2_ simultaneously at K52, R97, S215, K219; **(D)** The destabilization of Ω1-H4 gate. The detailed definitions of the variables are given in the Supplementary Materials, section 1. The vertical dashed lines mark the same events as in Figure 3.

The structural composition of each of the 6 components reveals the conformational change events occurring when each of them becomes dominant (and then wanes) in a specific time segment of trajectory frames. **Fig. 5A** identifies component 3 as the conformational state that is <u>initiated</u> with the establishment of the hydrophobic environment for the CHL tail (described in **Fig. 3C** and discussed above) and <u>ends</u> after the CHL is released. We note that although the information used by the RED algorithm include only data from protein dynamics (i.e., the contact matrices for protein residues and between the residues and PIP2 components of the membrane), and not about the CHL ligand, the analysis identifies the conformation that is dominant when the CHL is released. This suggests an intrinsic allosteric coupling mechanism between the ligand CHL and the membrane-bound protein, as quantified in a subsequent section on the allosteric communication mechanism.

We showed above that the CHL release state identified in the trajectories and corresponding to component 3 in the RED analysis, follows an intermediate pre-release state of the *holo*-StarD4 system. The transition from component 2 to component 3 of the RED analysis (termed event 2→3) is identified to correspond to the transition into the pre-release state. This transition event is indicated in **Fig. 5A** by the reduction in the weight of component 2 as that of component 3 is concurrently becoming dominant. The structural context of this event shows that it is initiated when StarD4 leans towards the membrane to attain the pre-release orientation at the time point indicated by the vertical dashed line at the 1840ns frame. We have shown in the RED analysis of StarD4 (Ref. (15)) that the specific conformational changes associated with such transitions between components are identified by direct comparison of normalized spatial arrays (i.e., the collections of residue pairs in-contact during the conformational states represented by each component) of adjacent components in the RED analysis. As described in the corresponding Method, this comparison identifies **<u>s</u>**tructure-**<u>d</u>**ifferentiating **<u>c</u>**ontact **<u>p</u>**airs (SDCPs) as the contact pairs that contribute significantly to the contact feature in one component, while making minimal contribution in the other component during the transition. These SDCPs are determinant elements for the interpretation of the conformational states represented by the components, as they comprise newly formed contacts in the transition from one component to another, as well as the contacts released during that transition. For transition event 2→3 the structural changes are summarized by the detailed, pairwise distance information given in the in the SDCP maps **Suppl. Fig 2** and in **Suppl. Tables 1-3**. These data report the conformational changes of StarD4 and the concurrent formation of PIP2 binding interactions during the transition into the pre-release state, and are quantified in **Fig. 5** and depicted in **Fig. 6** below.

**Figure 6.**
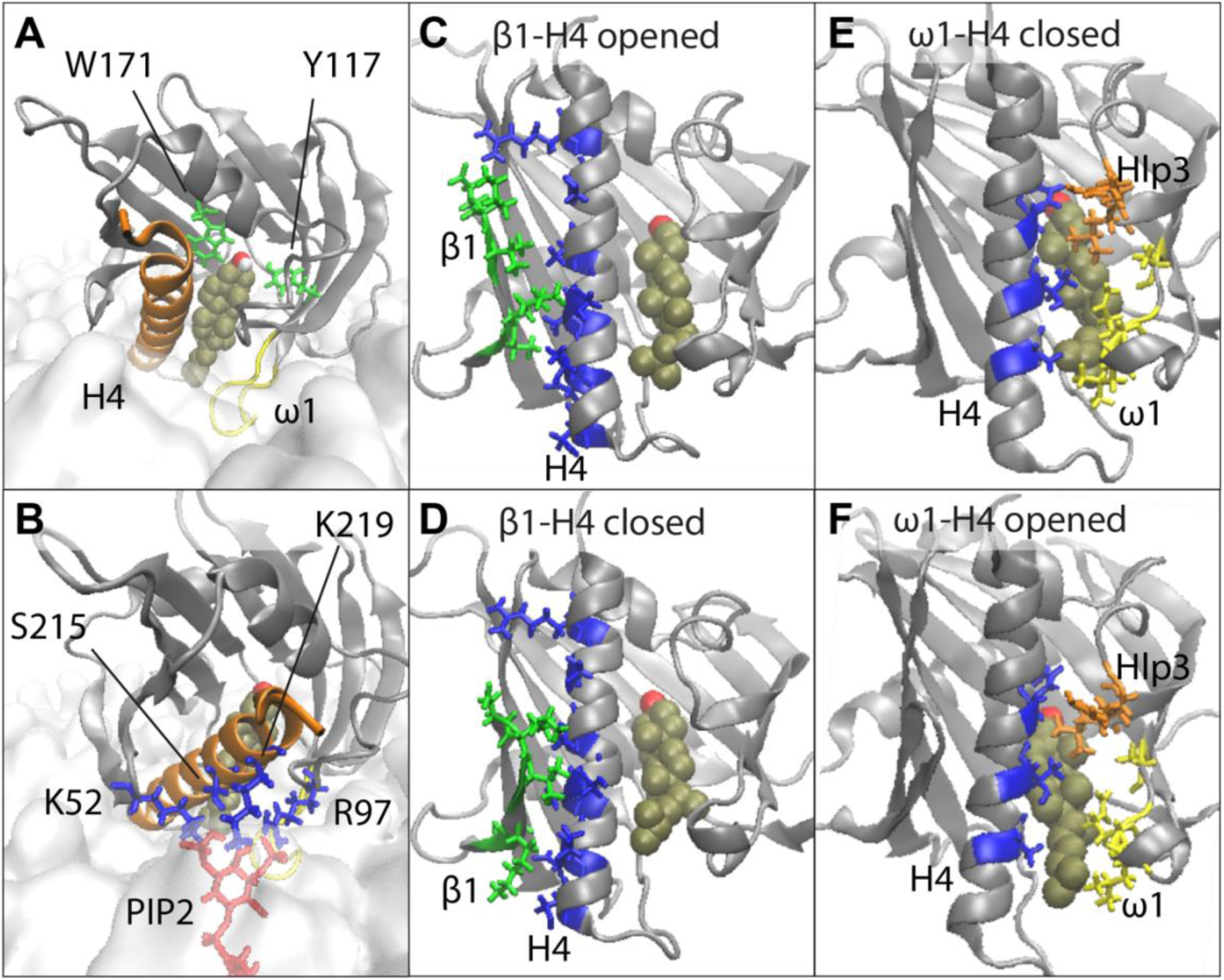
The set of StarD4 conformational changes and PIP_2_-StarD4 interactions shown by the RED analysis to lead to the release of cholesterol. **Panel (A)** shows a representative conformation of the StarD4-CHL-membrane complex in the pre-release state. The StarD4 molecule is rendered in gray cartoon with H4 in orange and Ω1loop in yellow; the CHL is rendered in VDW, and its binding sites W171 and Y117 are in green licorice. Membrane lipids are rendered in lines. **Panels (B,C,E,F)** show conformations of the structural elements identified in the RED analysis: **(B,C)** depict the shift of interactions between β1 and H4; **(E,F)** illustrate the destabilization of the interactions between Ω1 and H4. In **(B,C,E,F)** StarD4 is rendered in gray cartoon and the SDCPs in licorice. The SDCPs residues on β1 are shown in green, on H4 in blue, on Hlp3 in orange, and on the Ω1 loop in yellow. Panel **(D)** shows the PIP_2_-binding mode (across H4) that appeared in the pre-release state. The PIP_2_ is shown in red while the basic residues it interacts with are shown in blue.

Starting with its release from its salt-bridge interaction with the C-terminal of H4, β1 moves to form contacts with the N-terminal half of H4 (**Fig 5B**, **Fig. 6B&C**) and β1 extends downward to interact with the PIP2 whose phosphate bind simultaneously the K52 on the β1, the S215 & K219 on H4, and R97 on Helical loop 3 (H3) (**Fig. 3C**, **Fig. 6D**). As H4 distances itself from the Ω1 loop, the gate at the lower end of the hydrophobic pocket opens further towards the membrane, facilitating the formation of a hydrophobic pathway for CHL release (**Fig. 3D**, **Fig. 6E,F**). While this sequence of events recapitulates the conformational rearrangements we observed as well for the CHL translocation process in the dynamics of *holo*-StarD4 in aqueous solution, here, water penetration into *holo*-StarD4 embedded in the membrane, disrupts the interaction of the cargo CHL interaction with residues of β8 and β9 as the interaction with the bound PIP2 is established (marked by the second dashed lines in the time evolution shown in **Fig. 3C&E, Fig 5C**). The weakening of cargo-protein interactions as they are replaced by interactions with individual water molecules in the hydrophobic environment of the opening gate suggests this process as the preparation for cargo release from StarD4. and they are captured here by the transition event 2→3.

Thus, the structural interpretation of the detailed data from the RED analysis in **Fig. 6** brings to light the correlation between the specific modes of binding of PIP2 and the conformational change of StarD4 involving β1β2, the Ω1-H4 gate, and their interaction with PIP2. This is consistent with prior experimental results (2,9,12) showing that (i)-certain basic residues on the membrane-interacting surface of the protein have a critical role in mediating the cholesterol transport function of StarD4, and (ii)-that different PIP2 subtypes exhibit varying preferences for binding to these basic residues (12). We reasoned, therefore, that the characteristic mode of interaction between PIP2 and the basic residues in the pre-release state defined here is central to the recognition of the specific recognition of PIP2 by StarD4. The subsequent section presents an in-depth analysis of the PIP2-binding modes, and their role in determining whether, and how, StarD4 releases the CHL to membranes with different PIP2 compositions.

### The mechanism of cargo release from the *holo*-StarD4 protein is determined by the PIP2 content of the target membrane

#### The mechanistic role of the pre-release state in the delivery of StarD4 cargo

To reveal the specifics of conformational states visited by the membrane-embedded StarD4 in the CHL release process we employed the dimensionality reduction algorithm tICA (time-structure based independent component analysis) to analyze the trajectory data from Phase 2 of the simulations (see **Fig. 2**) as described in Methods. The projection to the tICA space, of the high dimensional data (i.e., all frames from the simulation trajectory) are projected onto a tICA space spanned by vectors composed of the collective variables (CVs) that describe the CHL binding modes and the function-related changes in structural motifs of StarD4. Six such CVs that determine the functional dynamics of interest were chosen to describe the conformational changes observed in the β1, H4, Ω1, Ω4, and CHL binding site motifs of StarD4 (**Fig. 7C**). These parameters were used to define the tICA space shown in **Fig. 7A-B**. The first two tICA vectors (**Fig. 7C**) captured 61% of the total dynamics of StarD4 in the membrane-embedded phase defined in **Fig. 2**, phase2.

**Figure 7.**
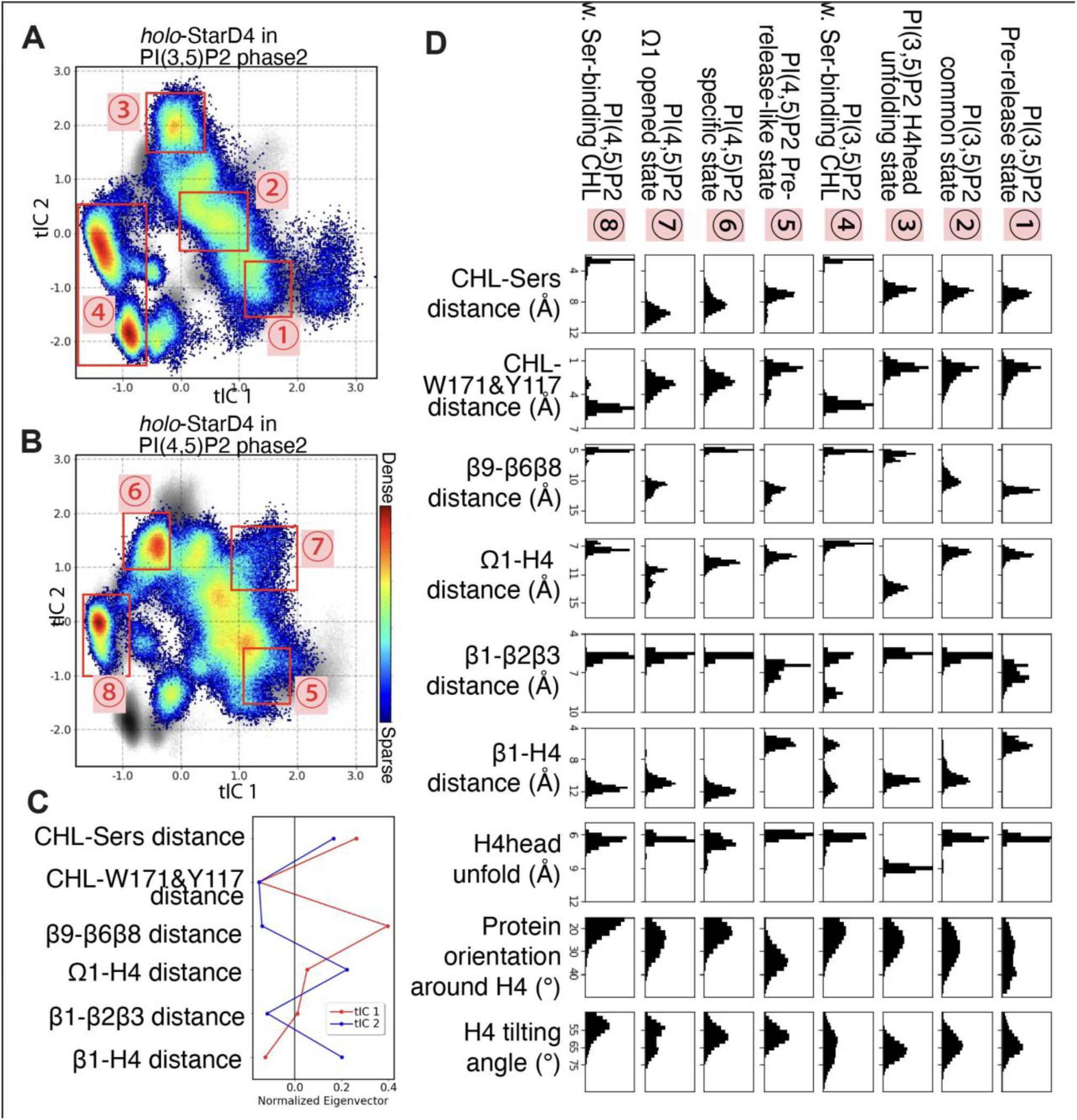
The characteristics of conformational states of membrane-embedded *holo*-StarD4. The high dimensional simulation data are projected onto a 2D tICA space defined on CVs describing the CHL binding modes and the conformational changes of StarD4. **Panels (A,B)** present the population density maps of the conformational space of *holo*-StarD4 embedding on the PI(3,5)P_2_-containing membrane **(in A)** and on the PI(4,5)P_2_-containing membrane **(B).** The tICA space is spanned by the two vectors (tIC 1, tIC 2) defined in **Panel (C)** by the contributions of individual CVs. The definitions of the CVs are listed in the Supplementary Materials, section 1. The color scale for the representation of population densities is shown on the right. The gray scale in the background represents the combined population density of StarD4s on three types of membranes **(Suppl. Fig. 3B)**. **Panel D** presents the structural characteristics of the pre-release state, the PIP_2_-subtype-specific states, and the states common to the protein embedded in membranes regardless of the PIP2 subtype. The states are identified the 8 red boxes in **(A,B)** and their characteristics are represented by probability density histograms of CVs that include the binding of CHL on the Ser-site and on the Trp-site; the interactions of β8-Ω4 corridor, Ω1-H4 gate, H4-β1, β1-β2&3 interactions, H4 partial unraveling, as well as the orientations of StarD4.

The tICA representations in **Fig. 7A-B** illustrate the specific ensemble of conformational states adopted by the membrane-embedded *holo*-StarD4 on membranes containing different subtypes of PIP2s. The PS-specific conformational ensemble and the combined ensemble are depicted in **Suppl. Fig. 3A,B**. Superimposing the time-evolution of RT1 onto the tICA space (**Suppl. Fig. 3C**) reveals that RT1 originated from state **2** in **Fig. 7A** and transited to state **1** where the spontaneous cholesterol release event occurs. The structural characteristics of these PIP2-specific states were discerned from the distribution of collective variables (CVs) representing the motif dynamics as detailed in **Fig. 7D**. Notably, it confirms that the pre-release state of StarD4 observed in RT1 as described above, is characterized by the interactions between β1-H4 (6^th^ column), the change in protein orientation (8^th^ column), and the wide opening of the β8-Ω4 corridor (3^rd^ column).

In addition to the pre-release state (**Fig. 7A&D**, state **1**), the other identified states (states **2-8)** include those in which CHL is at the Ser-binding site (**Fig. 7**, states **4**&**8**), and conformational states with CHL at the Trp-binding site. The conformational states with CHL at Trp-binding site include some that are found in all membrane-embedded *holo*-StarD4 trajectories (**Fig. 7**, states **2&5**) and others that are specific to one or the other kind of PIP2 subtype composition (**Fig. 7**, states **3,6,7**).

#### The characteristic features of the CHL-release observed in RT1 are reproduced as well in trajectories from Phase 3 simulations

To follow the CHL-release process under various conditions we conducted adaptive samplings simulations from initial seeds taken along the series of conformational changes that lead to the pre-release states of *holo*-StarD4. These pre-release states share similar characteristic features in each of the membrane systems studied – the PI(3,5)P_2_-containing membranes (**Fig. 7A**, state 1), the PI(4,5)P_2_-containing membranes (**Fig. 7B**, state 5) and PS-containing membranes. As shown in (**Fig. 2**, Phase3, orange set) the cumulative simulation time was 355 microseconds. In this set of simulations, the CHL release process described for **RT1** was successfully reproduced in two additional simulation on PI(3,5)P_2_-containing membrane (termed **RT2** and **RT3**), and in one trajectory from the StarD4 embedded in PS-containing membranes (**RT4**) which also shared features of the pre-release state. Additionally, we started a set of 24 trajectories from artificial pre-release states in which the β1-H4 interaction was maintained and the StarD4 was manually positioned in the strongly tilted orientation of the pre-release state (see **Fig 4**) on the PI(3,5)P_2_-containing membrane containing a PIP2 bound in the characteristic cross-H4 mode identified for RT1. One of these trajectories (**RT5**) also reproduced the CHL release process, but with cholesterol moving only halfway out of the pocket (**Suppl. Fig. 4A**).

The characteristic features observed in RT1 that re-appeared during the cholesterol release process in all these trajectories included the binding of an anionic lipid at K52 and R97 (**Suppl. Fig. 4B**), the β1-H4 interaction and destabilization of the Ω1-H4 gate (**Suppl. Fig. 4C**), the wide opening of the β8-Ω4 corridor (**Suppl. Fig. 4C**), and the orientation of StarD4 relative to the membrane surface (**Suppl. Fig. 4D**). While StarD4 may deviate from this pre-release orientation just before the release process in some of these trajectories, when cholesterol is destabilized from its binding sites in the pocket it consistently rearranges back into the rarely visited orientation on the membrane, leaning towards the hydrophobic pocket. This underscores the pivotal role that the StarD4 orientation relative to the membrane (**Fig. 4)** plays in accomplishing cholesterol transportation.

#### A low probability release mechanism from a high energy initial state

By initiating additional simulations in Phase 3 (**Fig. 2**, phase3, pink set, 334 μs) we probed the importance of the pre-release state formation on progressing to cargo delivery by explored possible alternatives for the starting points for CHL release pathways. Such simulations were initiated from different conformational states of StarD4 with the CHL bound at the Trp-binding site (**Fig. 7A,B**, states 2,3,6,7), as indicated by the scatter points on the tICA maps in **Suppl. Fig. 5**. Only two CHL release trajectories were observed (**RT6**, **RT7**), and remarkably, both of them started from a high-energy state (see initial frames situated at the upper right corner of state 7 in **Fig. 7B**) where the population density is very low. The dynamics of the transition to CHL release in the trajectories (**RT6**, **RT7**) are detailed in the Supplementary Information, **Suppl. Figs. 6**.

Intrigued by the possibility of cargo release from the alternative starting point, we assessed the likelihood of CHL release from the pre-release state shared by RT1-RT5, compared to a release from RT6&7 by comparing the free energy barriers of CHL release in these trajectories. To this end, we explored first the release paths using Steered Molecular Dynamics (SMD) simulations (26,27) as described in Methods. The initial frames were selected from the states of the pre-release conformations on PI(3,5)P_2_-containing membrane (**Fig. 7A**, state **1**) and on PI(4,5)P_2_-containing membrane (**Fig. 7A**, state **5**), as well as from the state containing the conformations with opened Ω1 on PI(4,5)P_2_-containing membrane (**Fig. 7B**, state **7**). The ligand CHL was steered along the z-axis towards to membrane at an average speed of 0.05 Å/ns for 400 ns in each run, with 24 replica runs per group. The free energy profiles were then evaluated with umbrella sampling protocols (see (28,29) and Methods) along the pathway resulting from the SMD.

The free energy profiles calculated for membrane systems containing either PI(3,5)P_2_ or PI(4,5)P_2_ are shown in **(Fig. 8**). The calculated total energy barrier for CHL release from the identified pre-release state was 2.6 kcal/mol (**Fig. 8A**), but it was 4.0 kcal/mol for the release pathway from the Ω1-opened state (**Fig. 8B**). This observation suggests that the kinetics of CHL release from the pre-release state would be approximately 10-fold faster than from the Ω1 opened state. The same free energy profile evaluations were applied to other highly populated StarD4 conformational states with CHL at the Trp-binding site (**Fig. 7**, states **2,5,6**), which showed higher free energy barriers compared to those starting from the pre-release structures on PI(3,5)P_2_-containing membrane (**Suppl. Fig. 7**).

**Figure 8.**
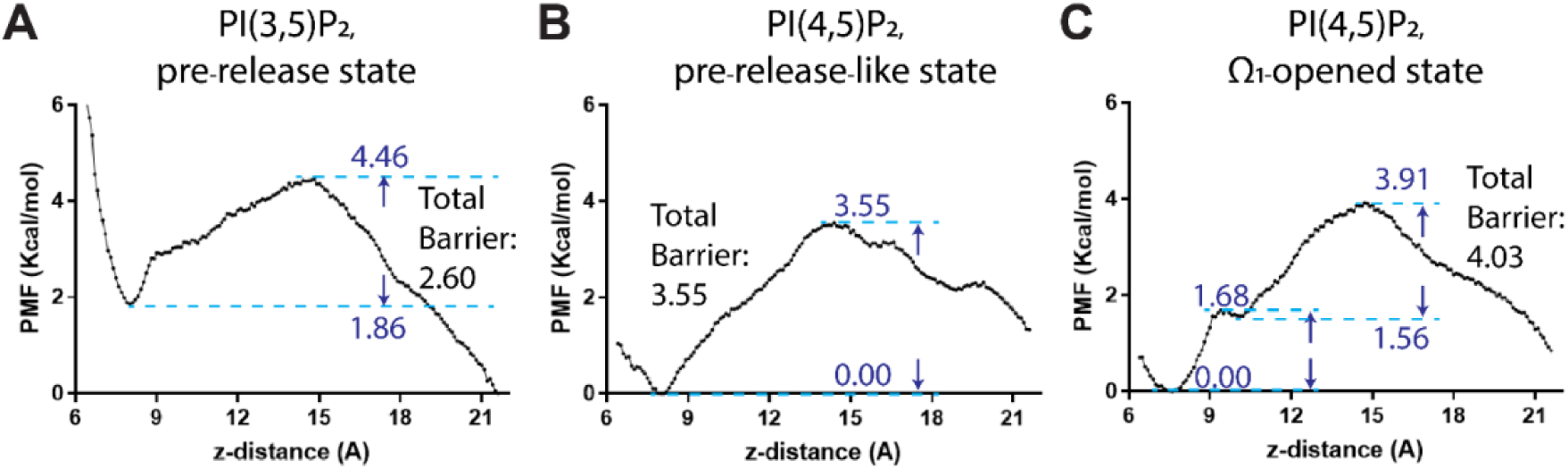
The energy barrier along the CHL release path sampled with Steered MD simulation and evaluated with umbrella sampling. The barrier is represented in terms of the calculated potential of mean force (PMF). The reaction coordination is the z-distance between the CHL and StarD4. Panels **(A-C)** show the PMF values for CHL release initiated from: **(A)** the pre-release state of StarD4 on PI(3,5)P_2_-containing membrane; **(B)** the pre-release-like state on PI(4,5)P_2_-containing membrane; and **(C)** the low probability Ω1-opened state on PI(4,5)P_2_-containing membrane. Local maxima and minima are labeled along the reaction pathways, and the calculated Total Barrier is indicated.

Thus, we find that CHL release from *holo*-StarD4 is energetically preferred from a state exhibiting the features we identified for the pre-release state. The results show that among the conformational states explored in the various long trajectories of the extensive simulations, the productive pre-release state of *holo*-StarD4 shares the following features: (1) A position of StarD4 on the membrane maximally tilted (leaning) in one particular direction (**Fig. 4**); (2) A conformation including an interaction between β1 and the H4 N-terminal region; (3) An open β8-Ω4 corridor; (4) A PIP2 lipid bound at K52 on β1 and R97 on H3. These features result from specific modes of interaction with the membrane PIP2 content and are communicated within the membrane embedded *holo-*StarD4 through the allosteric mechanisms we described. They determine the dynamics of CHL release as described below.

### The effect of PIP2-subtype binding modes to *holo-*StarD4 on the dynamics and repositioning of the β1 motif

A key feature of the pre-release state is the PIP2 binding at the K52 of β1 and R97 of H3 which positions it across H4. To learn whether the PIP2-StarD4 interaction mode correlates with specific conformational changes of StarD4, we investigated all the binding modes of the PIP2 membrane components to StarD4. We found three distinct groups of basic residues of StarD4 that interact with membrane anionic lipids, as illustrated in **Fig 9A**.

**Figure 9.**
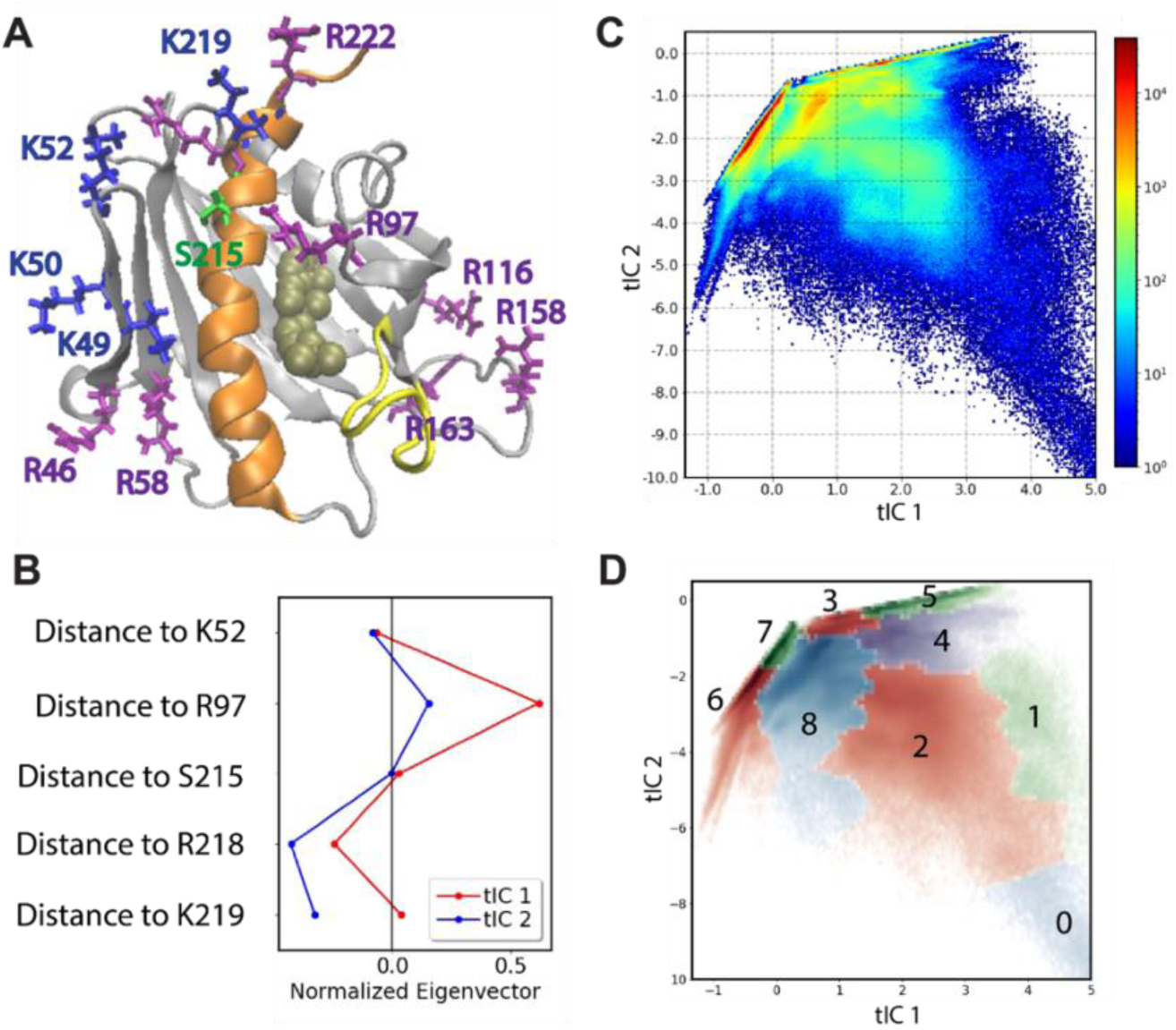
The probability map of PIP2 on putative interaction sites. **(A)** Basic residues on the StarD4-membrane interface. The StarD4-CHL complex is rendered the same way as figure 1, with the lysine’s shown in blue licorice, Arginine’s in Purple, and Ser215 in green. **(C)** Contributions of individual CVs to the first 2 dimension of the tICA space. The definitions of the CVs shown in this panel are detailed in the supplementary materials section 2. **(C)** The population density map of the 2D tICA space representing the StarD4-interaction modes of PIP2s. The color scale represents the density and is shown on the right. **(D)** The 2D tICA map is color-coded by the lipid binding modes 0-8. Based on their specific locations they are termed the *upper-group* (K52, R97, S215, R218, K219, R222), the *left-group* (R46, K49, K50, K52, R58) and the *right-group* (R116, R158, R163, R130, R194). Residues within a particular group are located adjacently, leading to a tendency for sharing of PIP2 interactions among them. The average number of anionic lipids that bind simultaneously on residues in each of the groups are summarized in Suppl. Table 4.

The specific mode of PIP2 binding in the pre-release state involves a PIP2 traversing the K52, S215, R218 and R97, which is an example of the PIP2-interaction at the upper-group. To facilitate the analysis of the PIP2 subtype interaction mode preferences in this *upper-group*, we constructed an Interaction Mode Landscape (IML) shown in (**Fig 9B-C)** as described in Methods. This binding mode map is further discretized into 9 macrostates (termed 0-8) based on their kinetic similarities (**Fig 9D**) as detailed in Methods. Each binding mode is characterized by the basic residues with which it interacts, as depicted in the histograms of PIP2-residue distances (**Suppl. Fig 8**) and are summarized in **Table 1** where binding mode 7 corresponds to that observed in the pre-release state.

**Table 1:**
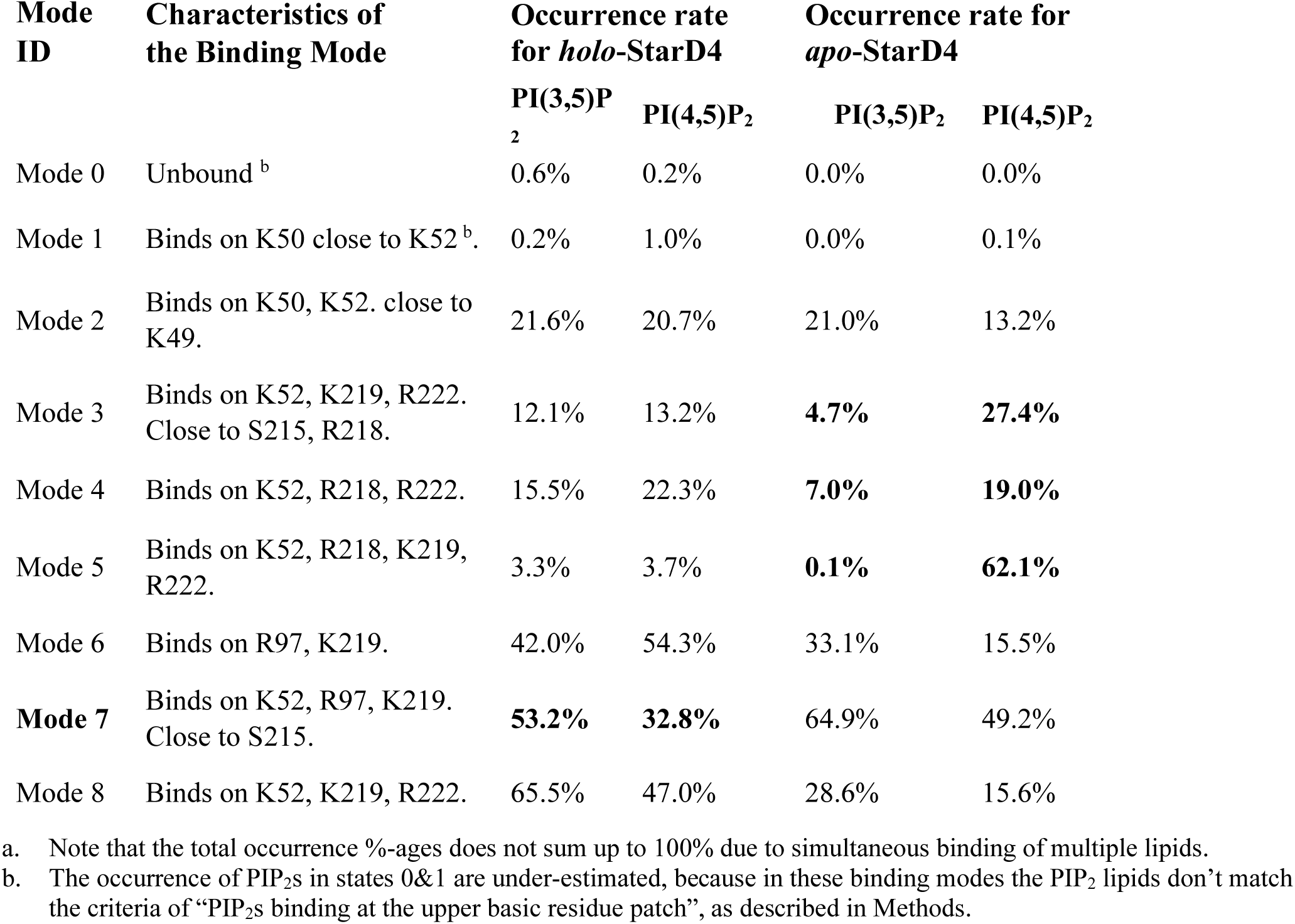
The occurrence rates^a^ of the PIP_2_ binding modes in the *holo* and *apo* StarD4 systems on membranes with different PIP2 compositions.

| Mode ID | Characteristics of the Binding Mode | Occurrence rate for *holo*-StarD4 |  | Occurrence rate for *apo*-StarD4 |  |
| --- | --- | --- | --- | --- | --- |
|  |  | PI(3,5)P<sub>2</sub> | PI(4,5)P<sub>2</sub> | PI(3,5)P<sub>2</sub> | PI(4,5)P<sub>2</sub> |
| Mode 0 | Unbound <sup>b</sup> | 0.6% | 0.2% | 0.0% | 0.0% |
| Mode 1 | Binds on K50 close to K52 <sup>b</sup> | 0.2% | 1.0% | 0.0% | 0.1% |
| Mode 2 | Binds on K50, K52. close to K49. | 21.6% | 20.7% | 21.0% | 13.2% |
| Mode 3 | Binds on K52, K219, R222. Close to S215, R218. | 12.1% | 13.2% | **4.7%** | **27.4%** |
| Mode 4 | Binds on K52, R218, R222. | 15.5% | 22.3% | **7.0%** | **19.0%** |
| Mode 5 | Binds on K52, R218, K219, R222. | 3.3% | 3.7% | **0.1%** | **62.1%** |
| Mode 6 | Binds on R97, K219. | 42.0% | 54.3% | 33.1% | 15.5% |
| **Mode 7** | Binds on K52, R97, K219. Close to S215. | **53.2%** | **32.8%** | 64.9% | 49.2% |
| Mode 8 | Binds on K52, K219, R222. | 65.5% | 47.0% | 28.6% | 15.6% |

a. Note that the total occurrence %-ages does not sum up to 100% due to simultaneous binding of multiple lipids.
b. The occurrence of PIP<sub>2</sub>s in states 0&1 are under-estimated, because in these binding modes the PIP<sub>2</sub> lipids don't match the criteria of "PIP<sub>2</sub>s binding at the upper basic residue patch", as described in Methods.

To study the subtype-based interaction mode preference, we separately projected the position of PI(3,5)P_2_ and of PI(4,5)P_2_ on the 2D IML map. As shown in **Fig. 10A,B** and **Table 1**, PI(4,5)P_2_ exhibits a preference for states 3,4 and 5 where its phosphate groups gather in a narrow region. In contrast, PI(3,5)P_2_ shows a preference for binding mode 7, where its phosphate groups bind across H4 from β1 to Hlp3 (**Fig. 10C,D**).

**Figure 10.**
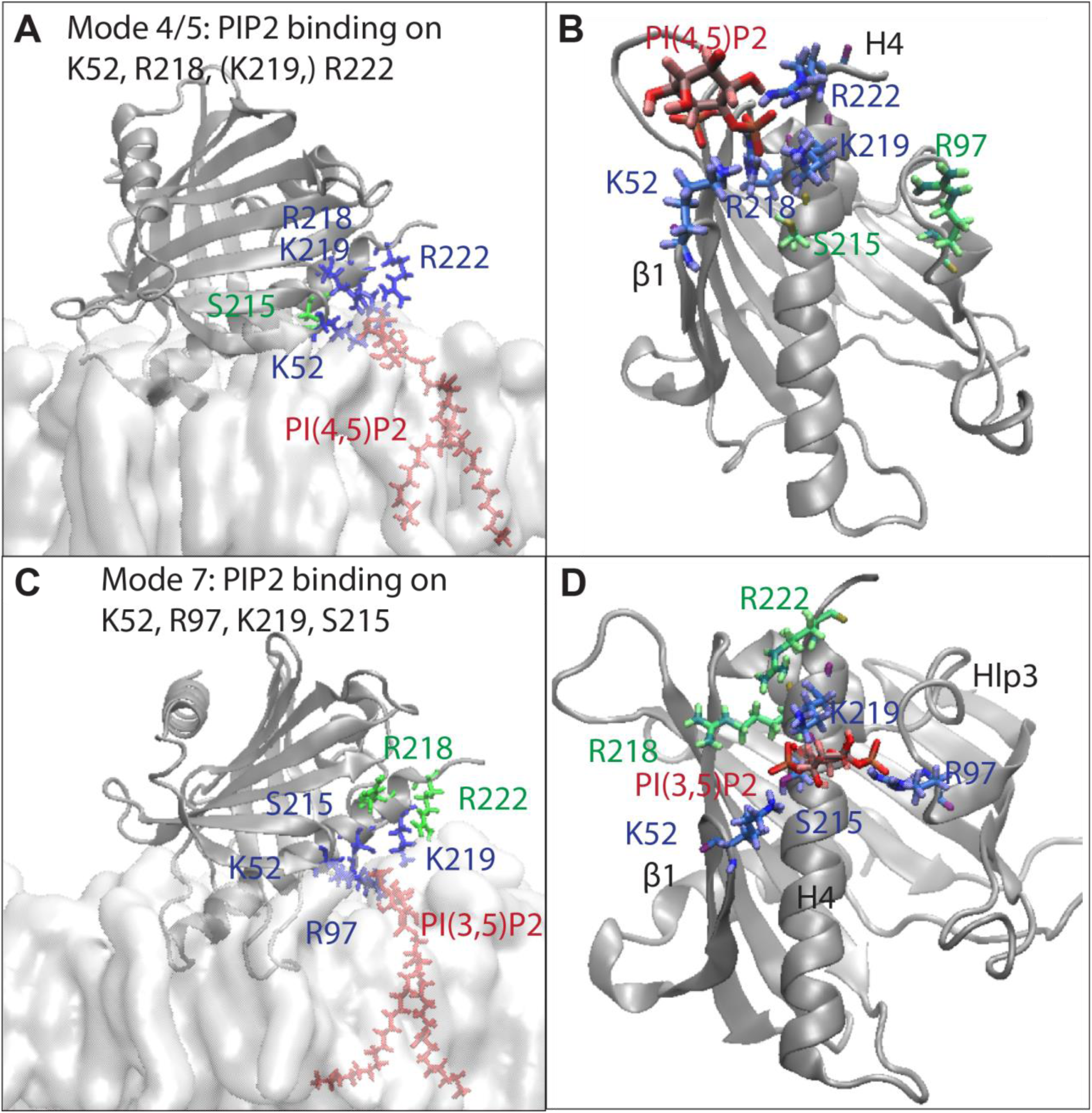
Representative conformations of PIP_2_-subtype-specific binding modes. **(A,B)** The modes of PIP_2_ binding to the *upper-group* residues K52 on β1, and R218 K219 R222 on H4, in lateral view **(A)** and upward view **(B)**. **(C,D)** The “cross-H4”-binding mode to the *upper-group* residues in which the PIP_2_ is binding with K52 on β1, S215 K219 on H4, and R97 on Hlp3, in lateral view **(C)** and upward view **(D)**. The StarD4 is rendered in gray cartoon. The *upper-group* basic residues K52 R97 S215 R218 K219 R222 are shown in licorice, with the PIP_2_-binding residues in blue and the non-interacting residues in green. The PIP_2_ is shown in red licorice. The membrane is shown only in the lateral views.

As detailed in a section above, the rare events detection algorithm (RED) showed that the conformational changes of StarD4 in the transition event 2→3 into the pre-release state with PIP2 in binding in mode 7 occur when the β1 establishes interactions with the N-terminal half of H4 (**Fig. 5A,C**). We therefore investigated the role of this binding mode by comparing the β1 to H4 distance in the presence and absence of a PIP2 binding at mode 7. The result (**Fig. 11A**) reveals the function-enabling correlation between PIP2 binding and the conformational changes in StarD4. Specifically, thecorrelation between the PIP2 binding mode and the conformational change at the β1-H4 is present throughout the entire simulation of membrane-embedded StarD4, not only in the pre-release state. This is important mechanistically because we had found this β1-H4 interaction to be associated with the opening of the β8-Ω4 back corridor and the destabilization of the Ω1-H4 gates on the pathway of cholesterol release (15). In the immediately following section the correlation between these rearrangements is quantified and the allosteric path is outlined in a structural context.

**Figure 11.**
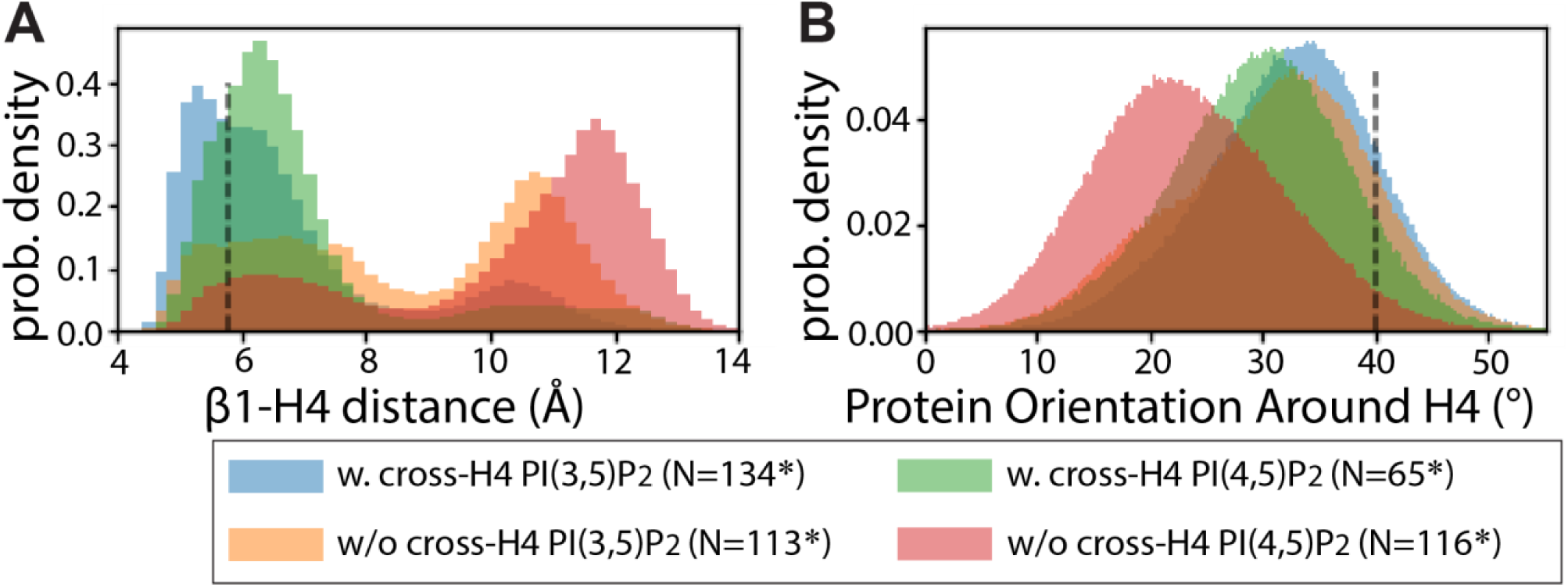
Comparison of the *holo*-StarD4 dynamics with and without PIP_2_ binding in the cross-H4 binding mode. **(A)** The probability density histogram of the β1-H4 distance. **(B)** The probability density histogram of the protein orientation. The vertical dashed lines indicate the values of these characteristic parameters in the pre-release state. Color code: <u>In blue:</u> the dynamics of StarD4 with a PI(3,5)P_2_ binding at the cross-H4 binding mode. <u>In orange:</u> the dynamics of StarD4 on PI(3,5)P_2_-containing membrane without a cross-H4-binding lipid. <u>In green:</u> the dynamics of StarD4 with a PI(4,5)P_2_ binding at the cross-H4 binding mode. <u>In red:</u> the dynamics of StarD4 on PI(4,5)P_2_-containing membrane without a cross-H4-binding lipid. <u>Statistical test:</u> with a time-interval of 2μs per snapshot to eliminate the autocorrelation, the simulations are equivalent to 134, 113, 65, 116 independent samples in the four categories separately. For the comparison of β1-H4 distance, the statistical significance are tested using Mann-Whitney U Test: with & without cross-H4-binding-PI(3,5)P_2_, p=3.4e^-15^, with & without cross-H4-binding-PI(4,5)P_2_, p=1.1e^-16^. For the comparison of protein orientation around H4, the statistical significance are tested using T Test: with & without cross-H4-binding-PI(3,5)P_2_, p=0.073, with & without cross-H4-binding-PI(4,5)P_2_, p=4.8e^-11^.

The other feature of the release state promoted by the binding of PI(4,5)P_2_ in mode 7 is the reorientation of the StarD4 relative to the membrane. In this special orientation the protein tilts towards the side of hydrophobic pocket (**Fig. 11B**). Collectively, these findings suggest that the occurrence of the cross-H4-binding of PIP2 is preferred in the pre-release conformational changes in StarD4, underscoring the mechanism of PIP2-subtype recognition by StarD4 and its role in the regulation of the cholesterol trafficking function.

#### The allosteric communication between the PIP2-binding modes and the dynamics of the CHL binding pocket

Having identified the relation between structural motifs of StarD4 involved in the conformational changes to the pre-release state of *holo-*StarD4 on the PI(3,5)P_2_-containing membrane, and the motifs responsible for the differential response to interactions with different PIP2 subtypes, we proceeded to quantify their allosteric communication with the cholesterol binding site with the N-body Information Theory (NbIT) analysis (24,25). The relevant component of the NbIT analysis, which can reveal the structural composition of the allosteric network connecting these sites and quantify its strength, is the normalized coordination information (NCI) between motifs. It is quantified from the information shared between a specified *receiver* motif and a *transmitter* motif, as described in Methods. The calculated NCI values between motifs of mechanistic interest are presented in **Table 2** using the residue indexes listed in Methods.

**Table 2.**
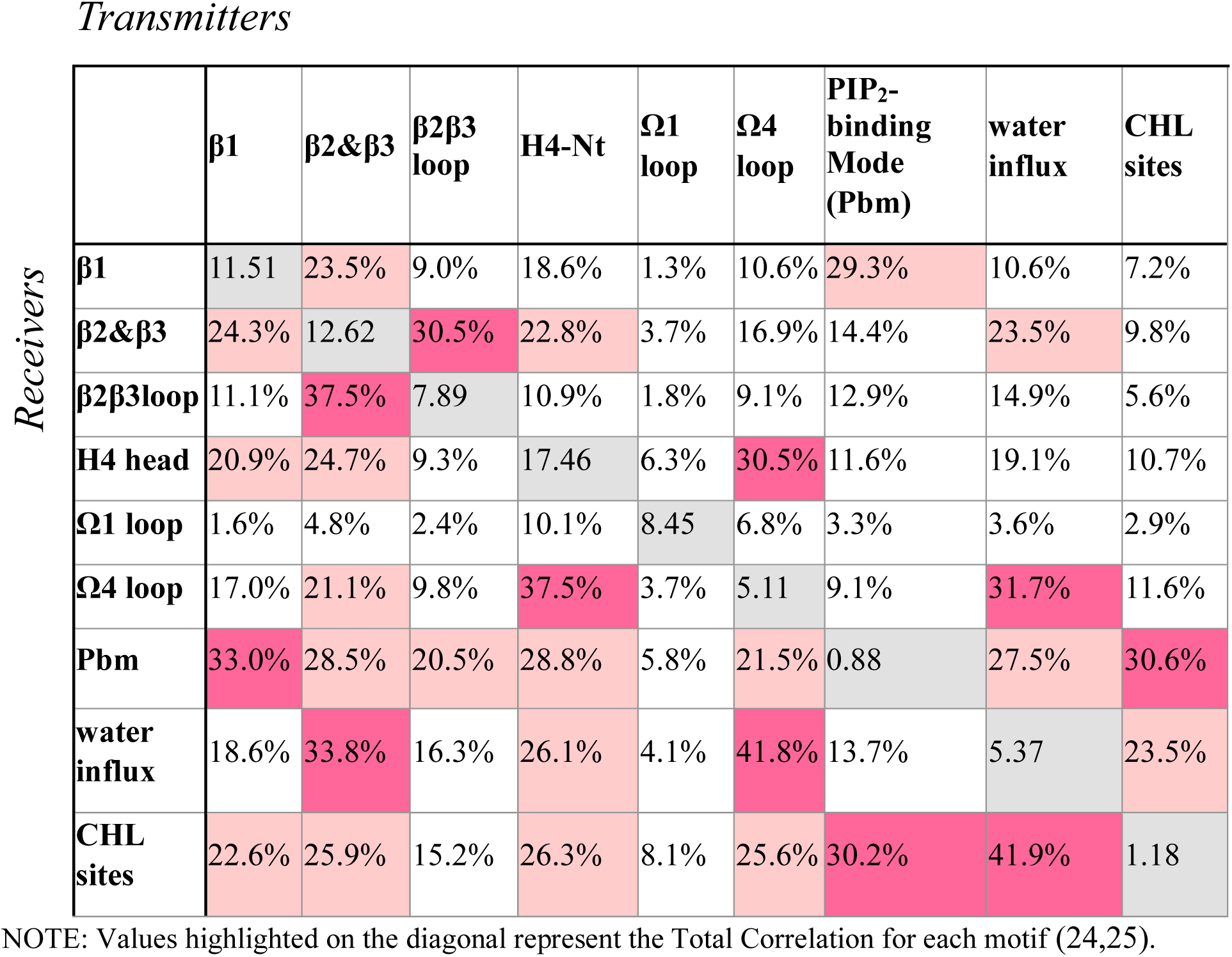
Normalized Coordination Information values calculated between various structural motifs in the *holo*-StarD4-membrane complex acting as *transmitters* (columns) and *receivers* (rows).

*Transmitters*
| | $\beta 1$ | $\beta 2 \& \beta 3$ | $\beta 2 \beta 3$ loop | H4-Nt | $\Omega 1$ loop | $\Omega 4$ loop | PIP<sub>2</sub>-binding Mode (Pbm) | water influx | CHL sites |
| --- | --- | --- | --- | --- | --- | --- | --- | --- | --- |
| *Receivers* |  |  |  |  |  |  |  |  |  |
| $\beta 1$ | 11.51 | 23.5% | 9.0% | 18.6% | 1.3% | 10.6% | 29.3% | 10.6% | 7.2% |
| $\beta 2 \& \beta 3$ | 24.3% | 12.62 | 30.5% | 22.8% | 3.7% | 16.9% | 14.4% | 23.5% | 9.8% |
| $\beta 2 \beta 3$ loop | 11.1% | 37.5% | 7.89 | 10.9% | 1.8% | 9.1% | 12.9% | 14.9% | 5.6% |
| H4 head | 20.9% | 24.7% | 9.3% | 17.46 | 6.3% | 30.5% | 11.6% | 19.1% | 10.7% |
| $\Omega 1$ loop | 1.6% | 4.8% | 2.4% | 10.1% | 8.45 | 6.8% | 3.3% | 3.6% | 2.9% |
| $\Omega 4$ loop | 17.0% | 21.1% | 9.8% | 37.5% | 3.7% | 5.11 | 9.1% | 31.7% | 11.6% |
| Pbm | 33.0% | 28.5% | 20.5% | 28.8% | 5.8% | 21.5% | 0.88 | 27.5% | 30.6% |
| water influx | 18.6% | 33.8% | 16.3% | 26.1% | 4.1% | 41.8% | 13.7% | 5.37 | 23.5% |
| CHL sites | 22.6% | 25.9% | 15.2% | 26.3% | 8.1% | 25.6% | 30.2% | 41.9% | 1.18 |

NOTE: Values highlighted on the diagonal represent the Total Correlation for each motif (24,25).

The pattern of communication revealed by the NCI values in Table 2 recapitulates the allosteric network reported recently (15) from simulations of the same system in aqueous solution, indicating their source in the structure of the protein. For example, β1 remains a strong coordinator to the cholesterol binding site in the membrane-embedded StarD4 although no direct interaction is formed between these motifs. The importance the of this allosteric connection between β1 and the CHL binding site is underscored by the finding that the in the membrane-embedded system it is the basic residues of β1 that respond to the PIP2-subtype and is further coordinated with H4 dynamics. Thus, the allosteric network that channels the information from β1 to the CHL binding site in StarD4 involves the information sharing between β1 and the β2, β3, the H4 N-terminal (H4-Nt) regions. In turn, β2, β3, H4 share dynamic information with β9 & Ω4-loop, and H4 β9 & Ω4 then directly gating the release of CHL (**Fig. 12A**).

**Figure 12.**
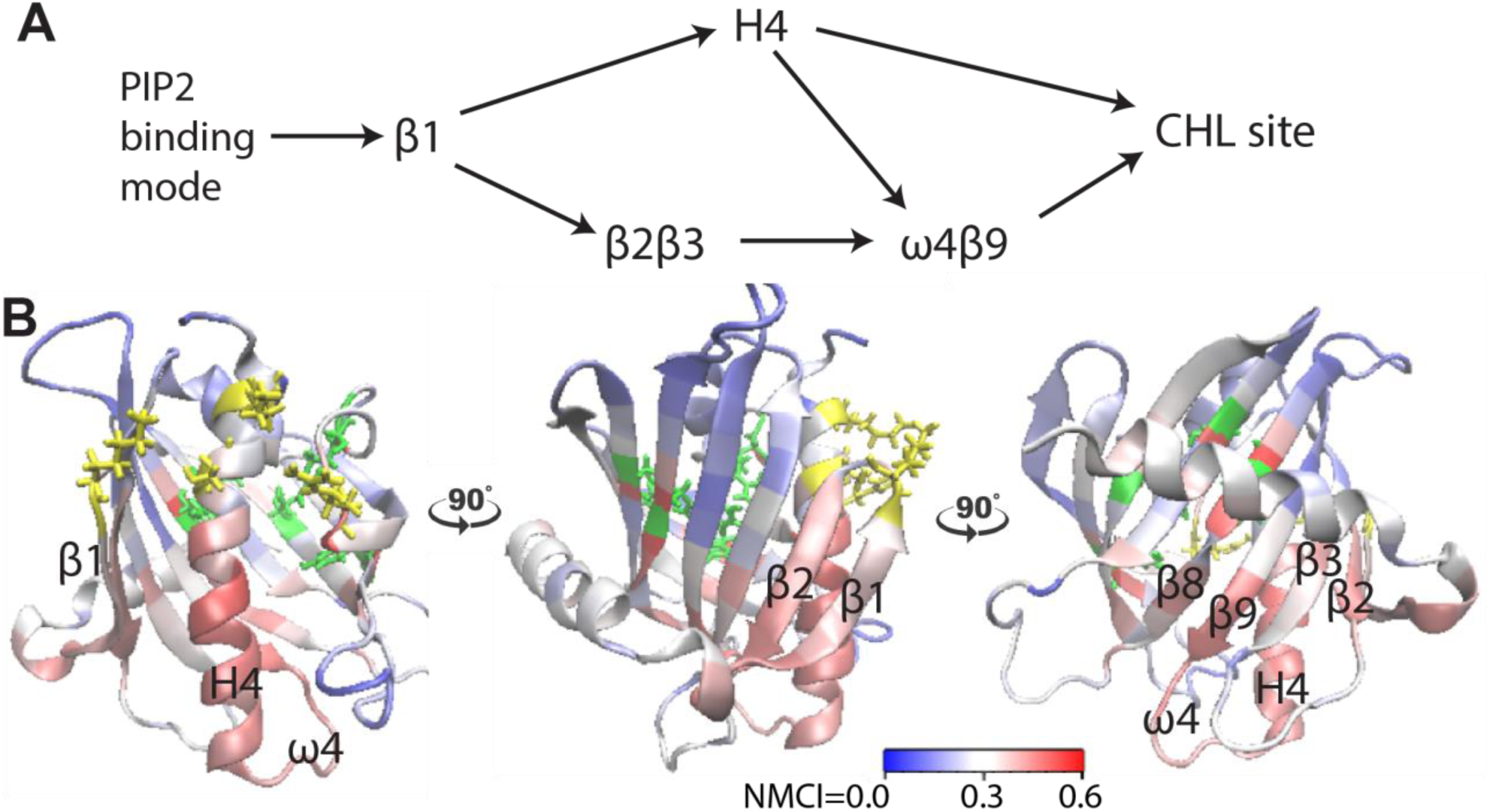
The allosteric coordination channel between the PIP_2_ binding site and the CHL binding site, which involves Helix4, β1, β2, β3 β9 sheets and Ω4 loop. **(A)** An allosteric model of information transmission from the PIP_2_ at cross-H4-binding mode to the cholesterol binding site. The arrows denote the direct information flow identified by the NbIT analysis between interacting motifs. **(B)** The cartoon model of StarD4 is colored according to the value of the normalized mutual coordination information (NCMI) calculated with the NbIT analysis. The color scale that quantifies the Normalized Mutual Coordination Information is at the lower right.

Remarkably, all the motifs in the function-related allosteric pathway are identified as strong coordinators to the cholesterol binding site (see last row of Table 2). The strong information sharing between the PIP2-subtype-specific binding sites on StarD4 (PIP2-binding mode, Pbm), and the β1 (Pbm/ β1=33%) confirms and interprets the central roles of β1 in differentiating the PIP2-subtype, and of the Pbm as a strong coordinator of the cholesterol binding site.

Thus, the values in Table 2 quantify the extent of information sharing between the motifs, illuminating their functional connectivity. As a necessary control, we calculated values of NCI for motifs that are not expected to be involved in allosteric pathways and found that these are much smaller than for the identified function-related motifs (see **Suppl. Table 5**). These small control values represent the background level of coordination information between unrelated motifs in the massive dynamics data contained in the MD trajectories.

The information sharing pattern identified by the NCI forms a coordination channel for information transmission from the PIP2 recognition site to the cholesterol binding site (**Fig. 12A**). This schematic channel is quantified in the NbIT formalism by the Normalized Mutual Coordination Information (NMCI) (24,25), calculated as described in Methods. The panels in **Fig. 12B** show this coordination channel in the structural context of StarD4. Together, the three panels in **Fig. 12B** depict the pattern of NMCI values of the transmission of information from the PIP2 binding site to the cholesterol binding site. The path of the PIP2 subtype-specific information transmitted through the strong information sharing among β1, β2, Ω4, H4-Nt to the CHL site determines the preference for pre-release conformational changes that govern CHL release in StarD4.

Notably, the pre-release state is achieved, and the CHL release stage is triggered, the allosteric communication pattern changes. This is revealed by the application of NbIT analysis to trajectories starting from the already formed pre-release state of StarD4 on PI(3,5)P_2_-containning membrane (see **Table 3**). The lower values indicate that as the necessary conformational changes in StarD4 have already occurred in the pre-release state, the dynamics of the CHL binding site are coupled primarily to residues N166 C169 on β8, and I189 T191 on β9, which are referred to as the “water influx” site in both **Table 2&3**. Indeed, we had observed that the last event before the release of CHL from the pre-release state of the StarD4 is the destabilization of the interactions of these motifs and the penetration of water to the site.

**Table 3.** Normalized coordination information between allosteric sites in *holo*-StarD4-membrane complex, analysis in the pre-release state only.

| Transmitters |  |  |  |  |  |  |  |  |  |  |
| --- | --- | --- | --- | --- | --- | --- | --- | --- | --- | --- |
| Receivers | | β1 | β2&β3 | β2β3 loop | H4-Nt | Ω1 loop | Ω4 loop | PIP<sub>2</sub>-binding Mode (Pbm) | water influx | CHL sites |
| | β1 | 11.93 | 18.8% | 5.6% | 14.0% | 1.3% | 6.3% | 24.6% | 5.5% | 3.9% |
| | β2&β3 | 20.1% | 11.25 | 27.6% | 20.5% | 2.7% | 16.9% | 9.8% | 24.6% | 7.5% |
| | β2β3 loop | 6.3% | 35.0% | 7.00 | 12.1% | 1.7% | 12.2% | 10.2% | 18.0% | 4.9% |
|  | H4-Nt | 16.5% | 22.6% | 9.3% | 16.70 | 3.4% | 24.2% | 8.7% | 12.7% | 6.3% |
| | Ω1 loop | 1.6% | 2.2% | 1.2% | 2.7% | 6.91 | 2.3% | 2.8% | 2.3% | 2.2% |
| | Ω4 loop | 7.0% | 12.3% | 7.0% | 21.2% | 2.1% | 5.54 | 4.0% | 17.8% | 3.5% |
|  | Pbm | 23.0% | 19.0% | 14.3% | 20.1% | 6.7% | 13.9% | 0.83 | 17.8% | 25.4% |
|  | water influx | 8.0% | 31.9% | 18.5% | 13.3% | 2.8% | 31.5% | 7.5% | 4.45 | 13.3% |
|  | CHL sites | 11.1% | 13.8% | 9.3% | 13.1% | 5.4% | 9.5% | 24.9% | 30.8% | 1.00 |

NOTE: Values on highlighted on the diagonal represent the Total Correlation for each motif (24,25).

### The structural motifs that respond dynamically to the recognition of specific PIP2-subtypes are identified with Deep Neural Networks (DNN) analysis

As the subtype-dependent PIP2 binding mode is found to be allosterically coupled with the cholesterol binding site through a series of conformational changes along an allosteric network in StarD4, we sought to identify the structural motifs that respond to the specific subtype of PIP2 in the membrane composition. To this end we used DNN analysis to differentiate first between the PIP2-subtype dependent StarD4 conformations, and then point out the molecular determinants responsible for the PIP2-subtype dependent StarD4 conformational changes.

To prepare the DNN to distinguish between StarD4 modes of interaction with PI(3,5)P_2_-containing membranes and with PI(4,5)P_2_-containing membranes in the *holo*-StarD4 trajectories we employed a supervised DNN training pipeline developed in our lab (30). The input datasets consisted of ∼46,000 structures defined by the coordinates of protein atoms only, and the DNN was designed to return the PIP2-subtype in the membrane with which the protein structure was interacting. These included data from both the membrane-embedding phase 1 and the membrane-embedded phase 2, sampled at 8ns time-intervals in >180 μs trajectories for each of the systems (**Suppl. Table 6**). The complete dataset was split into sets for training, validation, and test in the ratio of 56:24:20, respectively.

Test set results reached an accuracy of >85% in predicting the subtype of PIP2 (**Table 4**, left). As the “full test set” includes the trajectories of the membrane embedding phase of StarD4, in which it had not yet undergone any PIP2-induced conformational changes, it is difficult to improve further this accuracy. However, for the test set based on the membrane-embedded trajectory data, the DNNs achieved significantly higher accuracy, with 95.0% accuracy in predicting and differentiating the interacting PIP2 subtypes based solely on the conformational features of *holo*-StarD4 (**Table 4**, right).

**Table 4.**
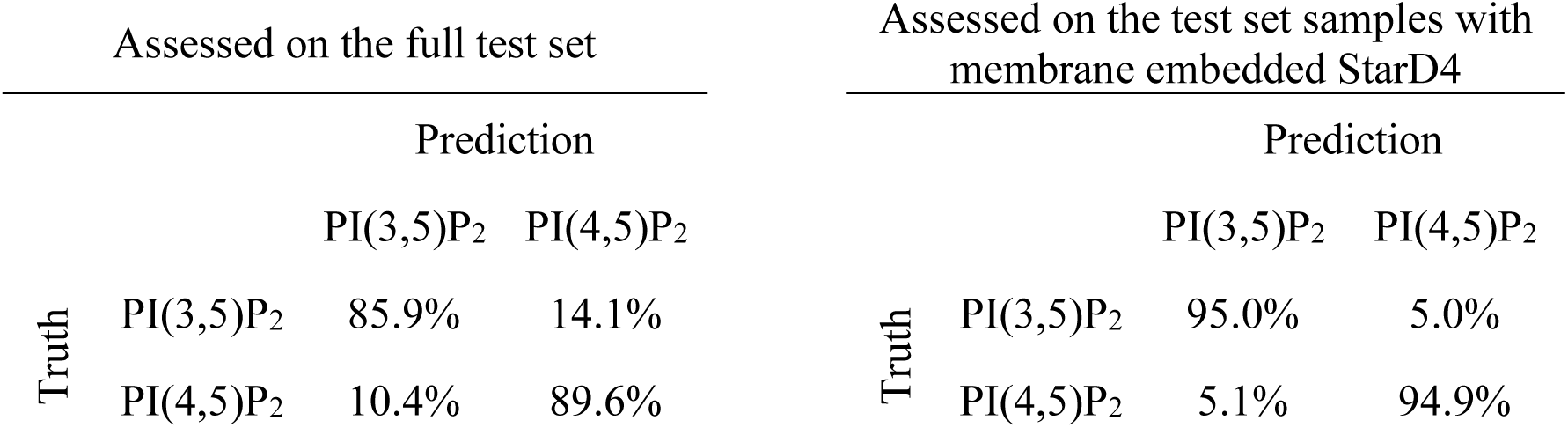
Results of the classification by DNNs.

| Assessed on the full test set |  |  |  | Assessed on the test set samples with<br>membrane embedded StarD4 |  |  |  |
| --- | --- | --- | --- | --- | --- | --- | --- |
|  |  | Prediction |  |  |  | Prediction |  |
|  |  | PI(3,5)P<sub>2</sub> | PI(4,5)P<sub>2</sub> |  |  | PI(3,5)P<sub>2</sub> | PI(4,5)P<sub>2</sub> |
| Truth | PI(3,5)P<sub>2</sub> | 85.9% | 14.1% | Truth | PI(3,5)P<sub>2</sub> | 95.0% | 5.0% |
|  | PI(4,5)P<sub>2</sub> | 10.4% | 89.6% |  | PI(4,5)P<sub>2</sub> | 5.1% | 94.9% |

As the high-accuracy results suggest that the neural networks were successful in recognizing the subtype-specific PIP2-induced conformational changes in the *holo*-StarD4, we proceeded to extract the salient structural features in the PIP2-subtype prediction by the DNNs. To this end we computed *attention heatmaps* featuring the atoms whose conformational changes exerted the greatest influence on the decision of the DNNs (**Suppl. Fig. 9**). **Fig 13** shows the motifs containing the high attention values: β1, β2, β3, Ω1-loop, β8, β9, and Ω4-loop. The salient residues on β1 are those interacting with anionic lipids as shown in (12), and critical components of the allosteric network described above (the β8-Ω4 corridor; the Ω1-H4 gate). Notably, the NbIT analysis had shown that these same structural motifs play a crucial role in coupling the CHL binding site with the anionic lipid binding site on β1.

**Figure 13.**
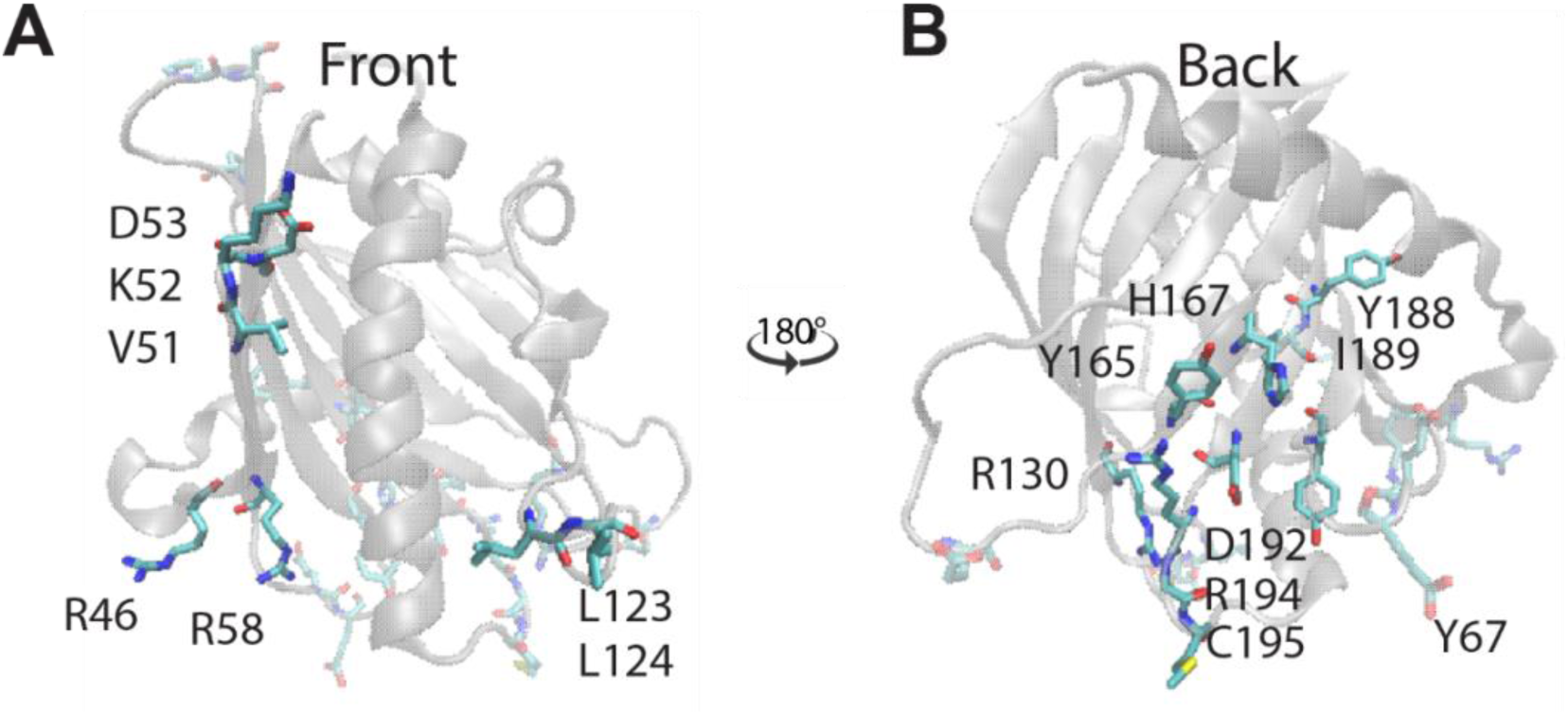
The molecular determinants identified by the DNN to be salient in the differentiation the interaction of StarD4 with PI(3,5)P_2_-containing membranes vs. PI(4,5)P_2_-containing membranes. **In (A,B)** the most important residues for the differentiation are labeled in licorice on the transparent cartoon model of StarD4 viewed from the front **(A),** and from the back **(B)**. The important residues on β1&2 and Ω1 are highlighted non-transparently when viewed from the front, while the important residues on β3 and the β8-Ω4 corridor are highlighted non-transparently when viewed from the back of the protein.

**Figure 14.**
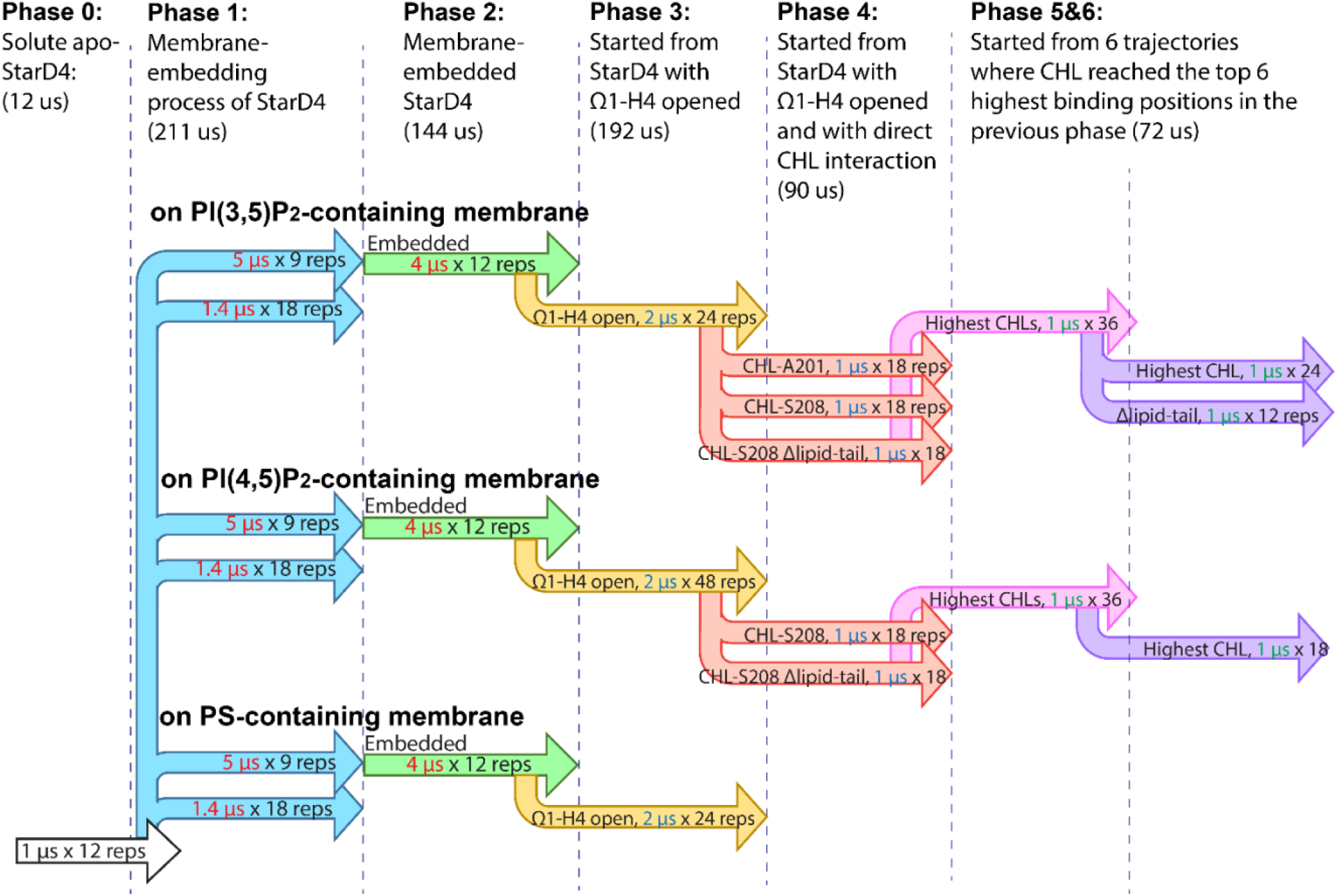
Schematic representation of the atomistic MD simulations of the *apo*-StarD4 membrane complex. The criteria for the choice of initial conformations, the simulation times, and the number of replicas for each set of simulation shown in the labels of the schematic plot.

#### Verification of the salient residues

We constructed a refined DNN based <u>only</u> on the coordinates of the salient residues in the motifs along the allosteric pathway described above, and trained it for the same recognition. Remarkably, this <u>specialized</u> DNN achieved an accuracy exceeding 96% (**Table 5**). While the information provided to the DNN by the data composed of only the salient residues is a proper subset of the original input, the results in Table 5 showed that the corresponding DNN managed to outperform the one based on the full protein. This suggests that the information derived from the salient residues is sufficient for accurate predictions and is also less affected by noise.

**Table 5.**
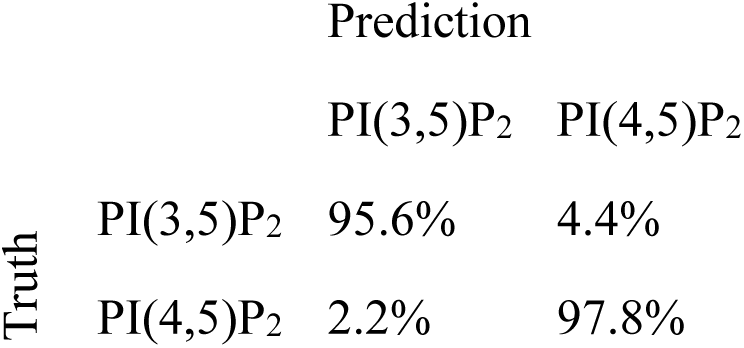
Results of the classification by DNNs trained using only the coordinates of the salient residues, assessed on the test set samples from membrane embedded StarD4.

|  |  | Prediction |  |
| --- | --- | --- | --- |
|  |  | PI(3,5)P<sub>2</sub> | PI(4,5)P<sub>2</sub> |
| Truth | PI(3,5)P<sub>2</sub> | 95.6% | 4.4% |
|  | PI(4,5)P<sub>2</sub> | 2.2% | 97.8% |

The recognition by the DNNs of these allosterically coupled motifs as pivotal discriminators, substantiates the distinctive dynamics of StarD4 responding to the difference in PIP2 composition of the membranes. This observation supports our mechanistic conclusion that information about PIP2 subtype is effectively transmitted through the identified allosteric network that regulates the dynamics of the CHL ligand.

The quantitative mechanistic inferences from the *holo-*StarD4 conformation analysis, the definition of PIP2 lipid binding modes, and the quantitative allosteric pathway definition come together in a mutually supportive manner. Together, they interpret in full structural detail the mode in which the dynamic response of membrane embedded *holo-*StarD4 recognition of different PIP2 subtypes regulates its preference for release of CHL according to the PIP2-subtype prevalent at the targeted membrane.

### The path of Cholesterol uptake by the membrane-embedded *apo-*StarD4

#### Repositioning of CHL by embedded apo-StarD4

Experimental evidence demonstrated that the PI(4,5)P_2_-containing membrane is a more efficient cholesterol donor to StarD4 compared to membranes containing only PI(3,5)P_2_, or PS (12). Employing the same investigated protocol that produced the mechanistic insights for *holo-StarD4,* we investigated the process of cholesterol extraction by StarD4 from extensive MD simulations of *apo*-StarD4. For membrane systems containing CHL and one of the negatively charged lipids (PI(4,5)P_2_, PI(3,5)P_2_ or PS) the simulations were carried out for 547 microseconds (>166 microseconds per membrane composition) using multi-phase simulations: Phase 1 (70μs) to observe the membrane embedding of StarD4; Phase 2 (48μs) to reveal the conformational changes of membrane-embedded StarD4; and Phase 3 for >48μs to explore the cholesterol extraction process by StarD4 with an open Ω1-H4 gate (see **Fig. 14**, Phases 1-3). The StarD4-membrane interactions examined in phases 2&3 revealed a strong response of the membrane cholesterol to StarD4 that was especially pronounced in PI(4,5)P_2_-containing membrane.

#### Membrane Cholesterol repositions in the presence of the membrane embedded apo-StarD4

At the concentration used, membrane-residing cholesterols are located predominantly beneath the membrane surface, with their oxygen closest to the lipid phosphate groups, and tend to have an inclination of ∼20° from the upright position (**Fig. 15G-I**). We found that with the StarD4 embedded in the membrane, a subset of membrane cholesterols neighboring either the Ω1 or H4 motifs is repositioned to reach above the membrane or tilt beyond 40° (**Fig. 15D-F**). This repositioning is more pronounced for cholesterols located right at the Ω1-H4 gate with interactions involving both motifs (**Fig. 15A-C**). Moreover, this repositioning, and especially the rising CHL, is more prevalent within PI(4,5)P_2_-containing membrane. It is less prominent in PI(3,5)P_2_-containing membrane, and weakest in PS-containing membrane.

**Figure 15.**
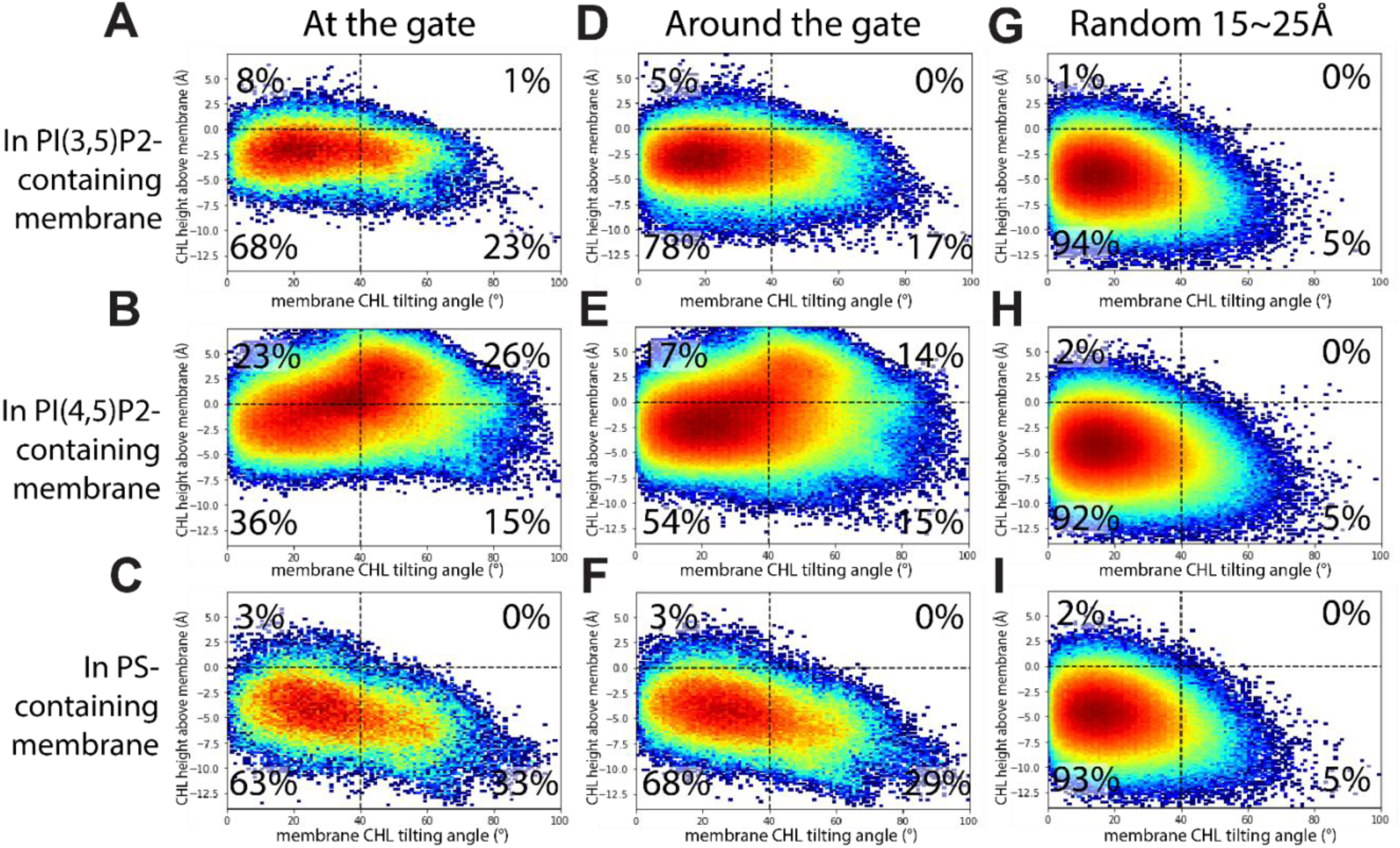
The probability distribution of the tilt and height positions of membrane-CHLs in relation to the membrane. **X-axis:** the tilting angle of the membrane-cholesterol; **Y-axis:** the height above the membrane surface. The positions are compared at three categories of positions relative to the embedded StarD4: when the membrane-CHL is at the H4-Ω1 gate **(A-C)**; when it is around the H4-Ω1 gate **(D-F)**; when it is far from the H4-Ω1 gate **(G-I)**. The negative lipid compositions of the membranes are shown in rows: **(A,D,G)** PI(3,5)P_2_-containing membranes; **(B,E,H)** PI(4,5)P_2_-containing membranes; **(C,F,I)** PS-containing membranes. The horizontal and vertical dashed lines are auxiliary lines that facilitate the comparison of the probability distributions by splitting the map to four regions, and the populations in each quadrant are indicated at the four corners of each figure. The figures are color-coded with densest regions in red and sparsest regions in blue. The detailed definitions of the variables and the criteria of the three categories are listed in the supplementary materials section 3.

Consistently, the dwell time of cholesterol at the Ω1-H4 gate follows the same trend (**Suppl. Fig. 10**). These observations align with experimental findings that the kinetics of cholesterol extraction by StarD4 is faster in PI(4,5)P_2_-containing membrane compared to those containing PI(3,5)P_2_ and PS (12), which suggests that this repositioning of CHL is a preparatory step in its process of extraction by StarD4.

#### The modes of interaction of the membrane-residing CHL with the embedded StarD4

Analysis of the simulation trajectories of *apo*-StarD4 in the three membrane systems showed three different sites of major interaction. These are indicated in **Fig. 16A-C** by the high-density regions in the 2D-space constructed for each membrane composition. The 2D-space is spanned by the distance of the CHL oxygen to StarD4 residues A201 (on the Y-axis), and to S208 (on the X-axis). The sites are indicated in the **Fig. 16A-C** panels, and the structural definition of the sites is illustrated in **Fig. 16D-F**. Site **1** represents A201-binding, Site **2** represents S208-binding, while Site **0** represents membrane-CHL molecules that are distant from both A201 and S208 and have no direct interaction with StarD4.

**Figure 16.**
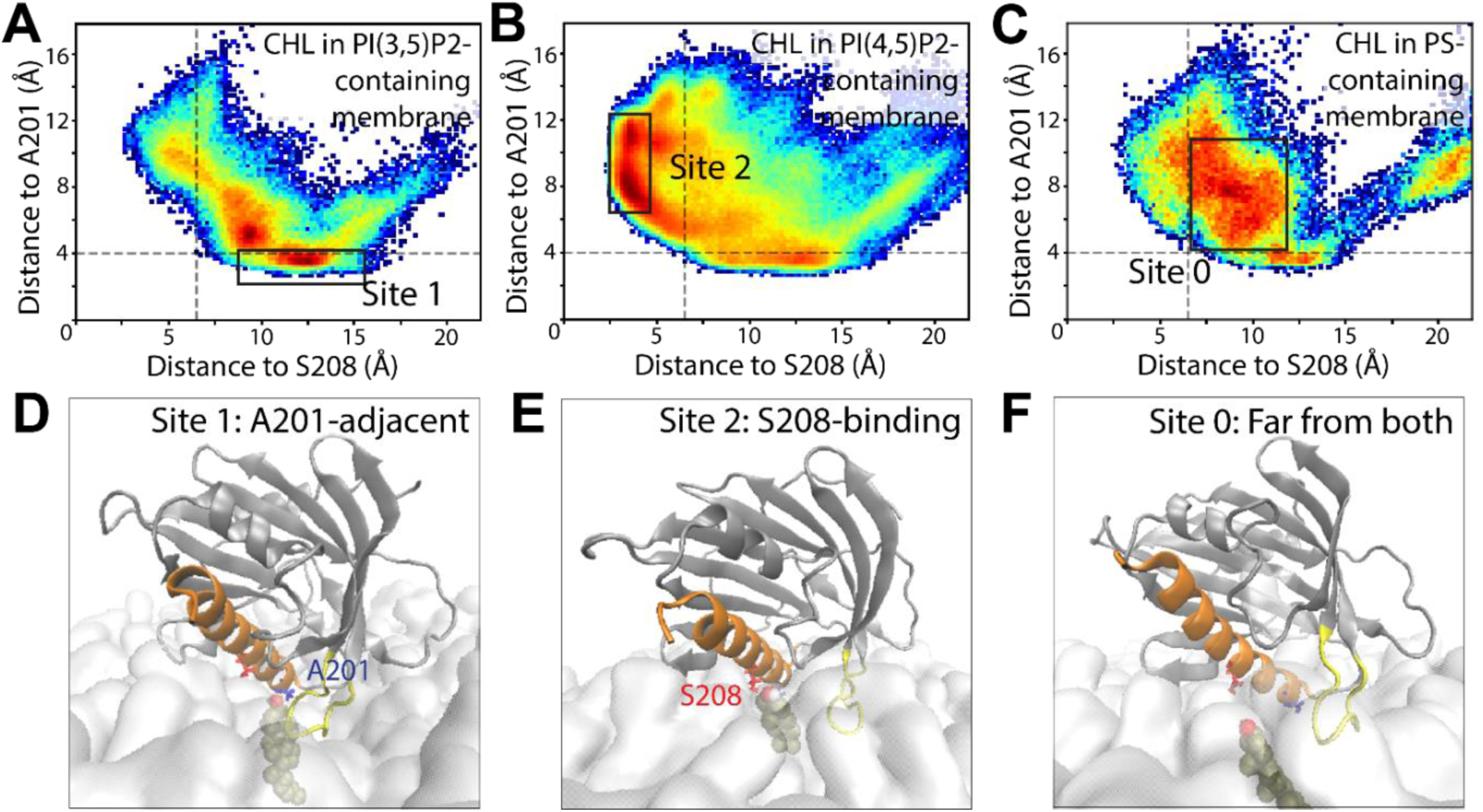
The interactions of membrane-residing cholesterols with membrane-embedded StarD4. **(A-C)** The probability distribution of the position of membrane-CHLs in relation to the StarD4 (X-axis: the distance to S208; Y-axis: the distance to A201), compared between the three types of membranes: **(A)** in PI(3,5)P_2_-containing membrane; **(B)** in PI(4,5)P_2_-containing membrane; **(C)** in PS-containing membrane. The figures are color-coded with dense regions in red and sparse regions in blue. The rectangles mark the metastable sites of CHL binding. **(D-F)** The representative conformations of *apo*-StarD4 complexes with membrane-CHL: Site 1, adjacent to A201 **(D)**; Site 2, binding to S208 **(E)**; Site 0, far from both A201 and S208 **(F)**. The protein is rendered in transparent cartoon in gray, with H4 in orange and Ω1 loop in yellow. The membrane cholesterol adjacent to StarD4 is shown in VDW. The cholesterol binding site residues are shown in licorice: S208 in red, A201 in blue. The membrane surface is denoted by the lipid phosphate headgroups shown in lines rendering.

Comparison of the distributions in **Figs. 16A, 16B, and 16C** shows that in membranes containing PI(3,5)P_2_ and PS, CHL favors Sites 0&1 (**Fig. 16A,C**), while Site 2 is prominent in the PI(4,5)P_2_-containing membrane but not in the PI(3,5)P_2_-containing membrane (**Fig. 16A,B**). Notably, the CHL molecule is reaching above the membrane surface <u>only at Site 2</u> (S208-binding), suggesting that the S208-binding site likely serves as a crucial intermediate state in preparing for the subsequent uptake of the CHL into StarD4. *In this scenario, the low occurrence of the S208-binding mode of cholesterol in the PI*(*3,5*)*P_2_-containing membrane, and even more so in the PS-containing membrane, relates directly to the observed slower kinetics of CHL uptake in these membranes*.

#### The pathway of Cholesterol uptake

The uptake of CHL by *apo*-StarD4 from PI(4,5)P_2_-containing and PI(3,5)P_2_-containing membranes was sampled in Phase 4 MD simulations (**Fig. 14)** with an adaptive sampling protocol starting from the Site 2 binding configuration described above (CHL at S208, **Fig. 16B&E**). With an open Ω1-H4 gate, adaptive sampling simulations were initiated from both the unaltered adaptive sampling seeds and the modified ones with “pocket-penetrating” lipid tails removed (**Suppl. Fig. 12**). Additionally, the *apo-*StarD4 with cholesterol at the A201-binding preferred on PI(3,5)P_2_-containing membrane (Site 1 above, CHL at A201, **Fig. 16A&D)** was also included as a set of adaptive sampling seeds. As shown in (**Fig. 14**, phase 4), this simulation setup results in a total of 90 replica runs, each of 1 microsecond in length. In the next adaptive step, 6 trajectories that had reached the highest position within the Ω1-H4 gate during the previous phase were selected from each membrane composition and used as initial points for new, 72 replica runs of 1 microsecond each. This iterative process was repeated across two phases to expand the sampling of CHL along its extraction pathway (**Fig. 14**, phases 5,6). As the cholesterol molecule moved up progressively from the gate to the W171, Y117 binding site, we represent these dynamics by mapping the interaction mode of CHL along its uptake pathway with its location described by the CVs listed in **Fig. 17A** (see further details in Supplementary Methods, Section 3).

**Figure 17.**
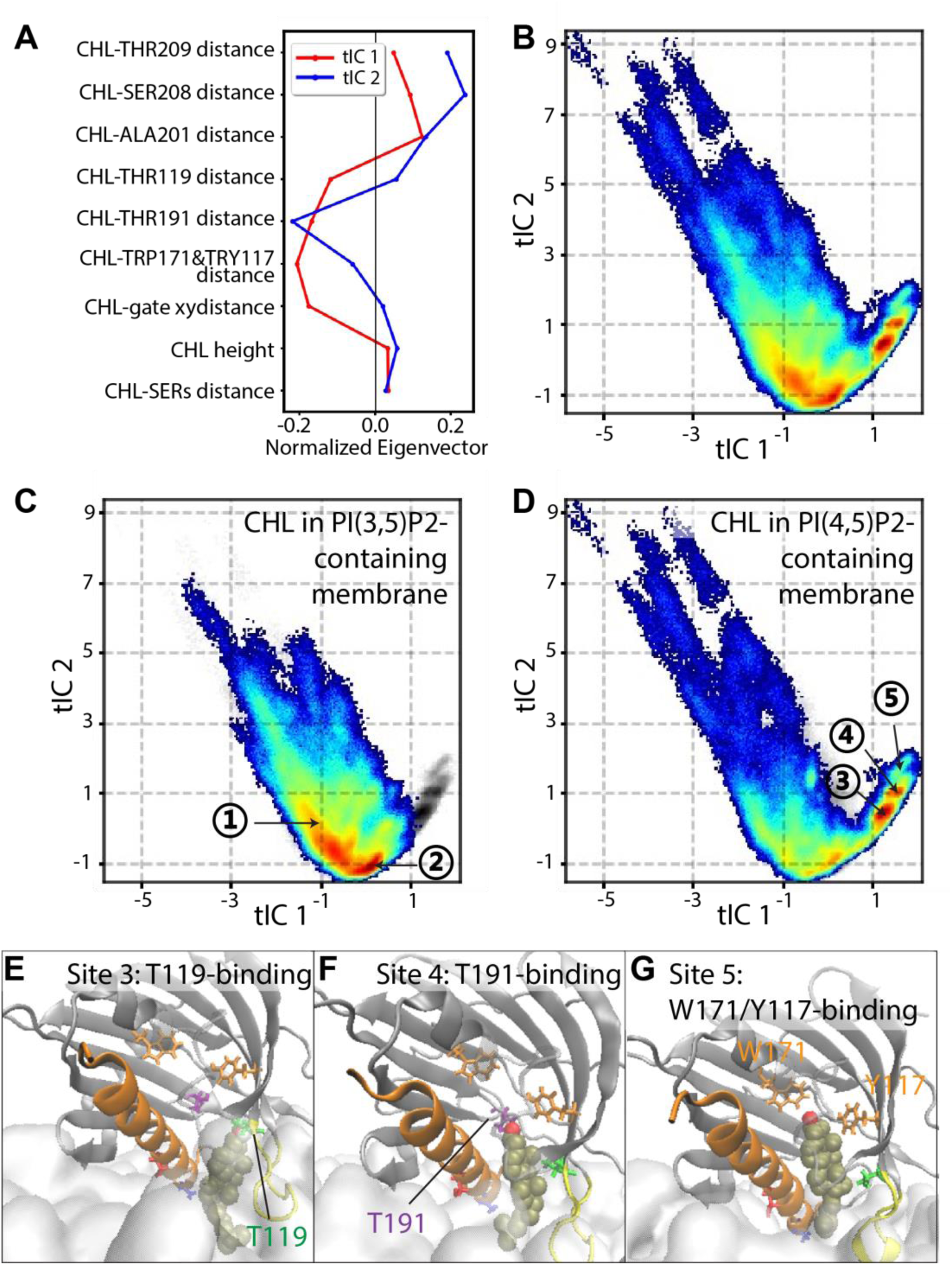
The cholesterol extraction pathway. (A-D) Extraction dynamics mapped on a 2D tICA space. **(A)** Contributions of individual CVs to the first 2 dimension of the tICA space they span. **(B)** The population density map of the position of the extracted cholesterol in relationship to StarD4. **(C)** On the PI(3,5)P_2_-containing membrane; **(D)** On the PI(4,5)P_2_-containing membrane. The figures are color-coded with dense regions in red and sparse regions in blue. Metastable CHL binding sites are marked in **(C)** and **(D)**. **(E-G)** The representative conformations of metastable CHL binding sites along the StarD4 hydrophobic pocket. The protein-cholesterol-membrane complexes are rendered as in Fig. 16. **(E)** Site 3, T119-binding; **(F)** Site 4, T191-binding; **(G)** Site 5, W171/Y117-binding. The cholesterol binding site residues are shown in licorice: S208 T209 in red, A201 in blue, T119 in green, T191 in purple, W171 Y117 in orange.

The CVs describing the upward movement of the CHL were used as parameters for in the tICA dimensionality reduction transformation of the CHL dynamics data as described above (e.g., see **Fig.7**) and detailed in Methods. The 2D map of the tICA space spanned by the first 2 tICA vectors which describe 92% of the total dynamics of the cholesterol along the uptake pathway (**Fig. 17B**) displays multiple metastable states. These include, in addition to Site **1** (A201-binding) and Site **2** (S208-binding) at the protein-membrane interface (**Fig. 17C)**, three stepwise binding states when the CHL is inside the hydrophobic pocket of StarD4: Site **3**, which represents T119-binding (**Fig. 17D,E**); Site **4** representing T191-binding (**Fig. 17D,F**); and Site 5 representing W171,Y117-binding (**Fig. 17D,G**). Site **5** marks the end of the uptake pathway as the W171,Y117-binding mode is commonly found in *holo*-StarD4. The identification of these sites as metastable states of the CHL dynamics is confirmed by their kinetic similarity, and is characterized by their CV distributions detailed in **Suppl. Fig. 13**.

The key comparative finding from this analysis of *apo-*StarD4 on membranes containing different PIP2 subtypes, is that the intermediate binding sites inside the hydrophobic pocket were observed exclusively for the PI(4,5)P_2_-containing membrane system (**Fig. 17D**), while the *apo-*StarD4 embedded on the PI(3,5)P_2_-containing membrane managed only rarely to extract a membrane CHL even up to the lowest binding site (Site 3 or 4) inside the gate (**Fig. 17C**). This difference agrees well with the experimental findings (12).

### Comparing the membrane interactions and conformations of the *apo-* and *holo*-StarD4 in their respective functional mechanisms

In the comparison of the *apo*-StarD4 dynamics on membranes with different PIP_2_ subtype compositions shown in **Fig. 18**, the data in shades of red in panel **A** show that the interaction with the N-terminal of H4 (H4-Nt) is absent for *apo-*StarD4 on PI(4,5)P_2_-containing membrane throughout Phases 4-6 (CV value for “β1-H4 interaction” is >10Å). From the allosteric pathway data for the recognition of the PI(4,5)P_2_ phospholipid binding mode to *apo-*StarD4, this conformational preference is due to the strong preference for binding mode 5 in PI(4,5)P_2_-*apo-*StarD4 interactions (cf. occurrence rates of binding mode probabilities of PIP2 phospholipids to StarD4 in **Table 1)**. In mode 5, the phosphate groups of the lipid are inserted into the upper-left corner on the basic residue patch (**Fig. 10A,B**), and this arrangement is likely to hinder β1 from shifting downwards and forming the interaction with the N-terminal of H4 (H4-Nt).

**Figure 18.**
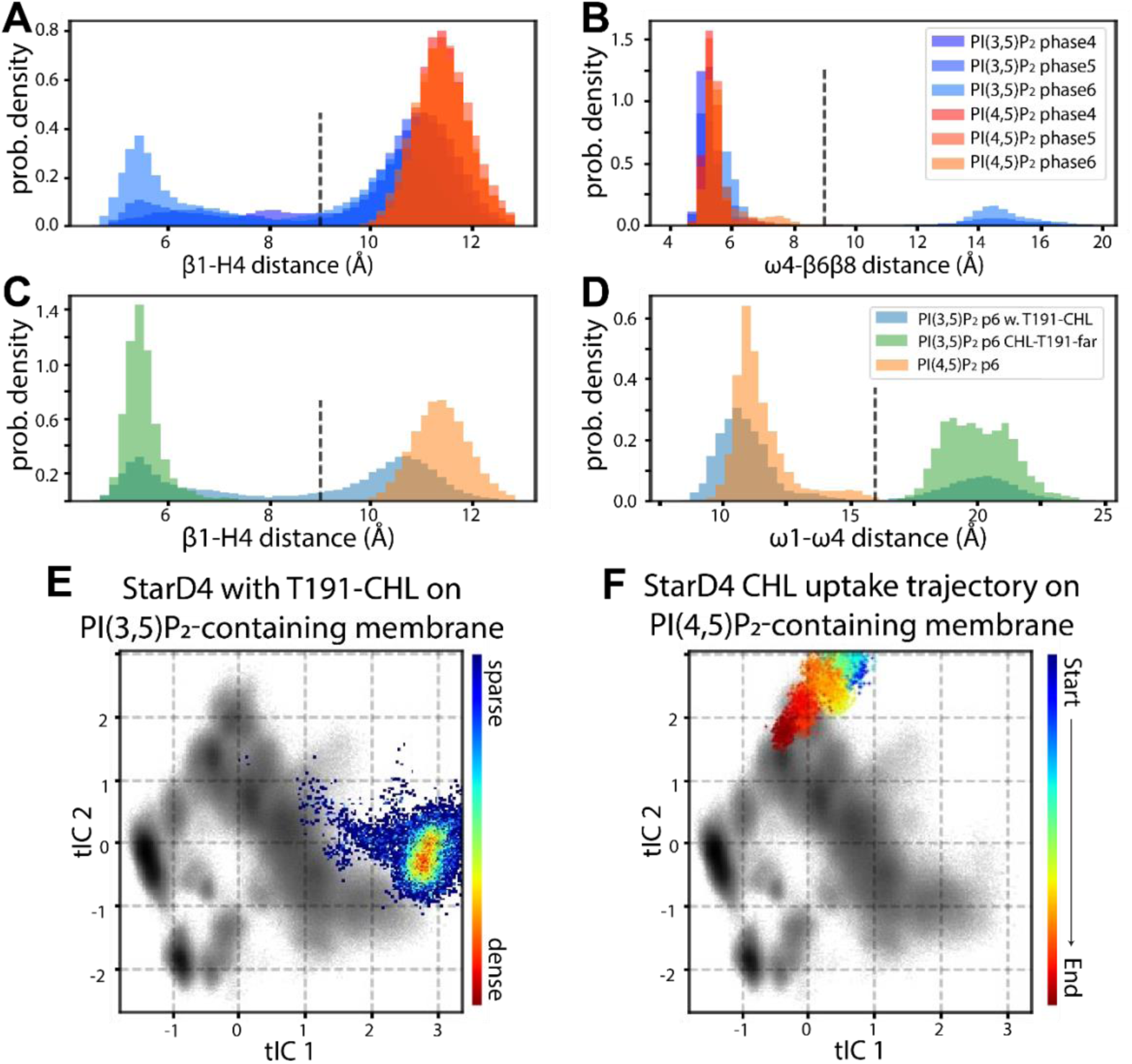
Comparison of the *apo*-StarD4 dynamics on membranes with different PIP_2_ subtype compositions and different modes of membrane-CHL binding. (**A,B**) The probability density histogram compared between the *apo*-StarD4 on PI(3,5)P_2_-containing membrane (red-orange scheme) and on the PI(4,5)P_2_-containing membrane (blue-purple scheme) during different phases of the adaptive sampling: **(A)** The interaction between β1 & H4; **(B)** The opening of the β8-Ω4 back corridor. The dashed line in Panels (A-D) indicate the state transition. **(C,D)** Color coded probability density histograms comparing StarD4 on PI(3,5)P_2_-containing membrane with a CHL close to the T191 binding site (green) and without any CHL close to the T191 binding site (blue). StarD4 on PI(4,5)P_2_-containing membranes is in orange. **(C)** The β1-H4 distance; **(D)** The Ω1-Ω4 distance compared. **(E)** The probability distribution of the conformation of the CHL-T191-adjacent *apo*-StarD4 obtained from the simulations on PI(3,5)P_2_-containing membranes, projected on the tICA space defined for the *holo*-StarD4, with the background represents the combined population density of *holo*-StarD4 in gray scale, as in Fig 7. The population density of *apo*-StarD4 is color-coded with dense regions in red and sparse regions in blue. **(F)** The time-evolution of the trajectory that successfully extracted a CHL from PI(4,5)P_2_-containing membrane, projected on the tICA space defined for the *holo*-StarD4. The evolution of the conformation encoded in the trajectory is color-coded from the starting point (in blue) to the end point (in red).

In contrast, **Fig. 18A** shows that for the *apo-*StarD4 on PI(3,5)P_2_-containing membrane, which has no PIP2 binding in mode 5) the β1-H4 distance diminishes throughout the same Phases (data in shades of blue), associating binding mode 7 of PIP2 which is associated with the increasing β1-H4 interaction. We had shown previously (15) that the movement of H4 towards β1 is correlated with the movement of Ω4-loop towards β2&3 and away from β8 & Ω1-loop, which leads to the opening of the β8-Ω4 back corridor. Indeed, the diminished β1-H4 interaction due to the large distance observed for PI(4,5)P_2_-membrane-embedded *apo*-StarD4, eliminates the opening of β8-Ω4 back corridor (**Fig. 18B**). The conformation with the back corridor β8-Ω4 closed on PI(4,5)P_2_-containing membrane generates a hydrophobic environment that stabilizes the incoming CHL that is being extracted from the PI(4,5)P_2_-containing membrane. This is supported by the observation that as the back corridor of the apo-StarD4 on PI(3,5)P_2_-containing membranes opens, it hinders the uptake. Indeed, in the few occurrences (4% of cases) when a CHL appears near THR191 in a position poised for incipient uptake, the protein features a temporarily strong β1-H4 interaction (**Fig. 18C**) and a widened back corridor between Ω1 and Ω4 (**Fig. 18D**). But this conformational state of the *apo*-StarD4 on PI(3,5)P_2_-containing membranes is akin to the pre-release state of *holo*-StarD4 which supports cargo movement in the opposite direction. **Fig. 18E** confirms this similarity of *apo*-StarD4 on PI(3,5)P_2_-containing membranes to the defined pre-release state by showing that the projection of the distribution of this conformation on the tICA space for the *holo*-StarD4 (see **Fig. 7** for details of the tICA map), lands close to the pre-release state. In contrast, **Fig. 18F** shows that the trajectory of the CHL uptake process from PI(4,5)P_2_-containing membrane projects to a canonical, highly populated PI(4,5)P_2_-specific state of *holo*-StarD4. This comparison of the dynamics shows that the *apo*-StarD4 on PI(3,5)P_2_ adopts a conformation favoring CHL release, not uptake, whereas the *apo*-StarD4 on PI(4,5)P_2_ adopts a conformation favoring the entrance and binding of CHL in the canonical site. *These findings are in agreement with experimentally observed preference ranking of CHL uptake from PIP2-containing membranes, and provide a detailed mechanistic explanation for these observations*.

Key findings from our probing approach, which is accessible at this point only computationally, predict that

***1.*** StarD4 interacts differently with membranes targeted by their enrichment in different negatively charged lipids it targets – PI(3,5)P_2_; PI(4,5)P_2_; and the PS – as a result of their distinct structural features;
***2.*** These targeted recognition lipids prefer different binding modes to the StarD4 protein, which allows it to recognize the membrane as a target for release of cargo uptake;
***3.*** A defined allosteric network involving specific combinations of structural motifs – β1, β2, β9, Ω4 and H4 – is involved in the dynamic response of StarD4 to the specific binding mode recognition through structural rearrangement events;
***4.*** The conformational changes produced by allosteric communication pattern triggered by the PIP2-subtype recognition dynamics, produce the specific, and different, conformations of the protein involved in either the uptake or release of sterol;

## Discussion

Experimental evidence has demonstrated that sterol traffic by StarD4 – a cytosolic protein in the family of STARD proteins involved in non-vesicular sterol transport between organelles – is modulated by its interaction with anionic lipids in the target membrane, particularly PI(4,5)P_2_ (9,12,31). As such traffic is essential for subcellular cholesterol homeostasis (1–7), we considered it essential to discover the nature of the molecular the mechanisms of enabling recognition of such anionic phospholipids by StarD4, and how it guides the functional mechanism. We therefore carried out extensive MD simulations mirroring experimental conditions to explore the processes of CHL uptake and release by the StarD4 protein interacting with membranes containing each of the anionic lipids examined in these experiments.

As described above, our quantitative analyses of the massive data from the trajectories of these extensive MD simulations have brought to light a detailed mechanistic description of the CHL uptake and release by StarD4 interacting with membranes that contain different PIP2 subtypes. The components of this quantitative mechanistic model emerging from the computational investigation described here, agree qualitatively and quantitatively with results and inferences from cognate experimental studies. By providing an otherwise not attainable interpretation of the experimental finding that we formulated in quantitative models of specific molecular events, they enrich the understanding of this form of non-vesicular cholesterol trafficking by StarD4.

Prior studies of such non-vesicular sterol transport, and sterol transfer proteins (19,20,22), have identified the START domains discuss in the Introduction, and a sterol binding pocket they contain. From these studies emerged the consideration of an orientation of the ligand in which it places the hydroxyl group at the base of the hydrophobic pocket (in StarD4, this means interactions with S136 and S147, **Fig. 1A**), but thus far no directly determined structures of any sterol-bound START domain have been reported. Some recent structural analyses (21) suggested that the size of the hydrophobic pocket of StarD4 is marginally insufficient to accommodate a cholesterol molecule, underscoring the necessity for a conformational change of the protein frame. Indeed, our simulation found a series of conformational changes of StarD4 in its complex with cholesterol. They showed (**Fig. 7D)** that the opening of Ω1-H4 front gate does enlarge the hydrophobic pocket and facilitates the translocation of CHL from the Ser-binding site to the Trp-binding site we defined. It is noteworthy that this Trp-binding-cholesterol-StarD4 complex we described recently (12,14,15) closely resembles the reported structures of cholesterol-binding StARkin domains of LAM proteins (**Suppl. Fig. 14**) (16–18). This similarity is especially relevant because previous *in silico* studies of the Lam4S2 protein have demonstrated the release of a sterol ligand from a similar complex (32). Similarly, we found from our MD simulations of the *holo-*StarD4 in an aqueous environment – before it embeds in the membrane (15) – that CHL can continue to move further down from the Trp-binding site to the farthest end of the hydrophobic cavity to position itself close to the entrance of the pocket, and we suggested that this represents a preparatory step before its release (15). The mechanism of CHL release from the *holo-* StarD4 embedded in the membrane that emerged from the results in the present work, supports this prediction of a preparatory state to which we refer as the pre-release state.

Thus, our findings from the analysis of the MD simulations mimicking membrane systems investigated experimentally (9,12) lead to a comprehensive molecular model of the dynamic mechanisms of CHL release from membrane-embedded *holo*-StarD4. We show that this mechanism is driven by a specific allosteric communication network along a path that depends on the nature of the anionic lipids enriched in the membrane. The functional outcome agrees with and interprets in a structural context the cognate experimental results that show clear differences between the functional behaviors of StarD4 when embedded in membranes enriched in either PI(3,5)P_2_, PI(4,5)P_2_, or PS phospholipids. In particular, we show how the modes of binding of different anionic lipids to StarD4 are different so that the activation of the allosteric channels (**Fig. 12A**) leads to different conformational changes, as described in **Fig. 11**.

The key findings underpinning the mechanistic model resulted from quantitative analyses of the time evolution of structure and properties in the MD trajectories of the various constructs with quantitative methods we have developed. These included (a)-the RED machine learning algorithm (15,23), (b)-the NbIT method (24,25,33) for quantification of the allosteric communication network among structural motifs in various StarD4 constructs, and (c)-the construction and implementation of a special machine learning DNN algorithm (30). The resulting mechanistic models and inferences, expressed in a discrete atomistic context, are poised for specific experimental probing as illustrated in our cognate publications (12,15,23,30,33–36). Indeed, the mechanistic model that emerged from our parallel studies of *holo-* and *apo-*StarD4 described here agrees with the experimentally observed CHL traffic differences in the assayed systems (12) and <u>shows how</u> the preferred functional activity of StarD4 (CHL uptake vs release) relates to the different PIP2-subtypes in which the tested membranes were enriched.

Thus, we uncovered here the discrete atomistic reasons for the kinetically preferred CHL release into membranes that contain PI(3,5)P_2_ lipids, and those favoring CHL uptake from membranes containing PI(4,5)P_2._ We showed that recognition of the target for one, or the other functional activity (uptake vs release of CHL), is determined by the differences in the preferred modes of binding of these anionic lipids to StarD4. The main structural insight is that the consequence of this selective recognition is the emergence of a defined state we termed “pre-release state” of StarD4 in which the Ω1-H4 gate and the β8-Ω4 corridor are open so that the ligand in the hydrophobic pocket is exposed to the membrane surface and destabilized by water molecules (**Fig. 6**). In tandem, the entire *holo-*StarD4 complex tilts strongly towards the membrane as defined by the spherical coordinates in **Fig. 4**. Our mechanistic outline shows that the CHL release process on the PI(3,5)P_2_-containing membranes starts from this pre-release state, and that this state is not visited in the uptake process that is preferred on PI(4,5)P_2_ containing membranes.

The detailed atomistic representations of the mechanism comprising the entire preparation and release process emerged from the analysis of the uncommonly rich data set from the ∼1.7 millisecond simulations trajectory we collected for *holo-*StarD4. **Figures 7, 11, 17, 18** illuminate the biophysical determinants for PIP2-subtype recognition by StarD4, and for the rearrangements of the β1, Ω4 and H4 structural motifs we found to be related to the interaction with anionic lipids (**Figs. 11&18)**. The ensuing water penetration and decrease of the interaction of CHL in the pocket induce the conformational changes conducive to cargo release.

We note here that we observed CHL release from an additional state (**Fig. 7D** state 7) in which both the β8-Ω4 back corridor and the Ω1-H4 gate are open, but the gate is held open by a penetrating lipid tail, not by the interaction between β1 and H4. While the essential structural determinants for CHL release are met by this alternative Ω1-H4 gate opening mode, the energy barrier of release from state 7 is much higher (**Fig. 8**). The energy profile reaches its peak at z-dist=15Å where the bulky hydrocarbon rings are passing through the gate in the absence of water penetration, and the barrier height suggests a 10x slower kinetics compared to the release from state 1. This underscores the importance of water penetration through the β8-Ω4 back corridor for CHL release, together with the facilitating role of the β1-H4 interaction in this process.

In stark contrast, the process of CHL uptake is observed exclusively for a StarD4 conformation lacking the shifted β1-H4 interaction and the opening of the β8-Ω4 back corridor (**Fig. 18**). Our mechanistic model of selectivity between uptake and release in the function of StarD4, which is dependent on different conformational changes of the membrane binding motifs, explains how these changes are driven by the allosteric network we defined in quantitative detail. Thus, the analysis of the extensive MD trajectory data revealed the allosteric network connecting the membrane-interacting region to the different regions of the StarD4 structure involved in the diverse modes of interaction with the specific PIP2 subtypes enriched in the membrane.

We found that the difference in the modes of interaction of StarD4 with different types of anionic lipids relates to the geometry of the anionic headgroups, from the unique polar site in PS, to the larger distance between the phosphate groups in PI(3,5)P_2_ than in PI(4,5)P_2_. Specifically, we found the modes of interaction by which the larger distance in PI(3,5)P_2_ favors the “cross-H4” binding mode that spans β1, H4 and Hlp3. In contrast, when the membrane contains the PI(4,5)P_2_ whose phosphates groups are adjacent, the “corner” binding mode (with K52, R218, K219, R222) is preferred. (**Fig. 10**, **Table 1**). Previous research has highlighted subtype preferences in the binding of PIP2 lipids to R46 and R58 at the N-terminal of β1 and the C-terminal of β2 (12). The different modes of β1 recognition by the PIP2 subtypes affect the dynamics in the allosteric network that regulates the actions of StarD4 in CHL traffic.

Regarding the findings from experiments with membrane in which the anionic lipid is phosphatidylserine (PS) (12,14), we also found that its smaller negative charge leads to a weaker adsorption energy of StarD4. Using the Mean-Field Model we calculated the adsorption energy when StarD4 approached a 10% PS-containing membrane the adsorption energy, and found that it is only −1.1 kBT compared to the −3.8 kBT energy observed when approaching a 10% PIP2-containing membrane. Correspondingly, the average number of PS lipids binding simultaneously at each of the three basic residue groups (**Suppl. Table 4**) is significantly lower than for the PIP2 lipids, the calculated anionic lipid dwell time is shorter for PS than for the PIP2 lipids (**Suppl. Fig. 11**), and the orientation of StarD4 on PS-containing membranes is less stable than on the PIP2-containing membranes (**Fig. 4**). The broader range of orientations of StarD4 on PS-containing membranes relates to a weaker repositioning effect of the membrane cholesterol (**Fig. 15,16**).

Our mechanistic analysis has brought to light the central role of the β1 strand in sensing the type of anionic lipid enriched in the membrane, and in transmitting the information to the CHL binding site through its dynamic interactions with H4, Ω4, and β9. Notably, both the β1 and β2 motifs are absent in the StARkin domain of LAM proteins despite the general conservation of various structural motifs among START and StARkin domains (19,20,22). This difference is, however, consistent with the difference in the functional mechanisms of the protein between StarD4 and the LAM StARkins which are constrained to specific membrane sites via their membrane anchors. The structural difference represents a functional adaptation that provides the soluble StarD4 family proteins with a membrane-target sensing mechanism that regulates their sterol trafficking directions.

We note that the comparative experiments investigating the sterol trafficking properties of StarD4 utilized the mouse StarD4 (mStarD4) protein (9,12,37) and thus our simulations were done with this same system. The membrane-interaction interface is highly conserved between mStarD4 and the human hStarD4, including the basic residues involved in the interactions with anionic phospholipids. Thus, most residues are identical, except for R97 and S215 in mStarD4 which correspond to S82 and G199 in the human, and R222 in mStarD4 which extends beyond the human StarD4 sequence (**Fig. 9A**). The S215A mutation has been reported to lower the kinetics of mStarD4 on PI(3,5)P_2_-containing membrane to a level that is significantly below than on PS, while the acceleration of the kinetics by the PI(4,5)P_2_-containing membrane relative to PS persists in the S215A mutant (12). Our findings are aligned with the experimental results, as we found the S215A mutation to block the favorite binding mode of PI(3,5)P_2_, which underscores the functional importance of the “cross-H4” binding mode we described and its subtype preference. On the other hand, the “corner” binding mode we found to be favored by PI(4,5)P_2_ is centered on the interaction at R218. In agreement with this finding, the R218A mutation has been reported (12) to slow the kinetics of StarD4 on PI(4,5)P_2_-containing membranes to the level measured on PS-containing membrane. Similarly congruent are the findings that mutations of K52 and K219 – which we showed to have essential roles in both the PI(3,5)P_2_ “cross-H4” binding mode and the “corner” binding mode favored by PI(4,5)P_2_ – impede StarD4 cholesterol transportation on anionic membranes (9). Together, these findings substantiate the insights we obtained about the mechanistic role of the specific binding motifs for each PIP2 subtype and their components in modulating the cholesterol transport dynamics of StarD4.

Naturally occurring mutations of StarD4 have been documented in the context of dysfunctions associated with various diseases. While expression rates of StarD4 have been related to cancer (38,39), it remained unclear how mutations of the basic residues may contribute. For example, the naturally occurring R202Q mutation in human StarD4 (corresponding to a mutation of R218 in the mouse StarD4 discussed here (**Fig. 10A,B**)) has been associated with 13 malignant melanoma and breast carcinoma samples in five independent studies (40–45). Similarly, the G199C mutation (corresponding to S215 in mouse StarD4 (**Fig. 10C,D**)) has also been reported in a case of skin melanoma (46). As our results underscore the mechanistic role of subtype specific PIP2 binding modes in the modulation of cholesterol transport function of StarD4, the dysregulation caused by mutations in the preferred PIP2 binding sites can explain a role in disease.

In conclusion, the PIP2-subtype-dependent regulatory mechanism of the StarD4 CHL uptake and delivery functions that emerged in this study, leverages differences in the dynamic response of the protein (both the *apo* and *holo* forms) to interactions with different anionic lipids. As described, these differences are propagated through the allosteric channels to the CHL binding site to modulate the stability of the cholesterol cargo, thereby regulating the conformations and the orientations of the protein on the target membranes that determine the steps and directions of the cholesterol traffic in cellular processes.

## MATERIALS AND METHOD

### Modeling the initial structures

#### The cholesterol-StarD4 complex

The structure of *apo*-StarD4 was derived from the X-ray structure (PDB ID: 1jss) where residues 24 to 222 are resolved (8). To complete the structure, the missing residues K223 and A224 are added using Modeler 9.23 software, resulting in the conformation of StarD4 segment spanning residues 22-224, with acetylation on the N-terminus and carboxylation on the C-terminus (47). The initial structure of cholesterol bound (*holo*) StarD4 was obtained by docking the cholesterol into the hydrophobic pocket of *apo*-StarD4 using the Schrodinger Induced Fit Docking protocol (48–50). Both the *apo-* and *holo-*StarD4 constructs were then solvated in 0.15M K^+^Cl^-^ ionic aqueous solution and equilibrated using the NAMD simulation platform version 2.12 (51).

#### Assembly of the initial lipid membrane

Explicit all-atom lipid membrane models were constructed with 400 lipids in symmetric bilayers composed of a 44:23:23:10 mixture of POPC, POPE, cholesterol, and anionic lipids, (either POPS, POPI(4,5)P_2_, or POPI(3,5)P_2_), which is consistent with the experimental conditions that revealed the PIP2-subtype-dependent StarD4 kinetics modulations (15). The initial structure of membrane bilayers was assembled using the CHARMM-GUI webserver (52). The full equilibration of the membrane patch was done with StarD4 restrained at a 2Å distance adjacent to the membrane as described in the section below.

#### Construction of StarD4-membrane complexes

A three-stage simulation protocol was employed to obtain the StarD4-membrane complexes:

**Stage 1.** *<u>Equilibration of StarD4 in aqueous solution.</u>* The conformational space of the StarD4 constructs prior to its interaction with membrane was explored for *apo-* and *holo-*StarD4 in aqueous environments in all atom MD simulations with OpenMM software (53). The protein was solvated in a 70×70×70 Å^3^ water box ionized with 0.15M K^+^Cl^-^, resulting in a system containing approximately 32,500 atoms. The OpenMM simulations were conducted in NPT ensemble (T=310K, P=1 atm) using a 4fs integration time-step and a Monte Carlo Barostat. As described recently (15), 12 statistically independent replicas were generated after equilibration for each system and simulated for a duration of 1μs/each for the *apo-*StarD4 system, and for 33.6 μs/each for the *holo-*StarD4 system. The analyses and results were previously reported (15). Following the simulations, clustering analysis was performed to identify distinct conformations, resulting in 3 representative conformations for *apo-* StarD4, and 9 snapshots in 3 representative conformation states for *holo-*StarD4. These were used to build the atomistic models for the simulations of StarD4-membrane complexes.

**Stage 2.** *<u>Distribution of the membrane anionic lipids in the electrostatic field of approaching StarD4</u> <u>molecules.</u>* To speed up the convergence of the membrane reorganization in the electrostatic field of the approaching StarD4, we used the Mean-field model (MFM) approach developed in the lab to assess the impact of long-range electrostatic interactions between StarD4 and membrane (12,54–57). MFM sampling accelerates the exploration under important degrees of freedom, particularly electrostatics and lipid mixing, which play crucial roles in the long-range interaction to establish an optimized state for the StarD4-membrane combination. This allowed us to investigate the effect of the mutual interaction on the orientation of StarD4 as it approaches the negatively charged membranes, as well as the corresponding rearrangement of anionic lipids in the membrane in response to the presence of StarD4 before the initiation of the atomistic MD simulations of StarD4 embedding in the membrane. The detailed definition and equations used in the MFM are presented in the Supplementary Method section 4.

**<u>Stage 3.</u>** *<u>Equilibration of the mutual StarD4/membrane interaction at 2Å above the membrane</u>*. The mutually adapted system from Stage 2 was solvated in 0.15M K^+^Cl^-^ ionic solution. The equilibration of the system consisting of ∼138,000 atoms was carried out in 3 steps with NAMD version 2.12, at a temperature of 310K in NPT ensemble using Monte Carlo Membrane Barostat, with the isotropic XY ratio as established previously (12). The 3 consecutive steps start with restraining forces of 1 kcal/mol·Å^2^ applied to the protein backbone atoms, the heavy atoms of the docked cholesterol, and the heavy atoms of the membrane lipids heads. After the first 0.5 ns simulations the restraints are decreased gradually, first to 0.5 kcal/mol·Å^2^, and then to 0.1 kcal/mol·Å^2^ in the subsequent runs of 0.5ns/each.

Following these 3 equilibration steps, the subsequent MD simulations were carried out with the OpenMM software (53), using 4 fs integration time-step, in NPT ensemble (T=310K, P=1 atm) under Monte Carlo Membrane Barostat. These started with a 500 ns equilibration phase in which the relative height of StarD4 (z-distance to the membrane surface) and the orientation of StarD4 (the z-distance differences between the left-side-half and right-side-half of StarD4, and between the top-side-half and the bottom-side-half the StarD4) are strongly restrained with a force constant of 20 kcal/mol·Å^2^ This constraint ensured proximity and consistent orientation of StarD4 with the membrane, facilitating the equilibration of membrane lipid rearrangement without direct contact. After the initial 500 nanoseconds, all these constrains were released, allowing StarD4 to establish direct contact with the membrane, and initiating the production run of the StarD4-membrane systems.

### Production run simulations of the membrane embedded *holo*-StarD4

Production runs of the membrane embedded *holo*-StarD4 were carried out with the OpenMM software with the same parameter and conditions as above (4 fs integration time-step, NPT ensemble (T=310K, P=1 atm), Monte Carlo Membrane Barostat). The 3-phase design of the entire unbiased simulation protocol of the membrane embedded *holo*-StarD4 is shown in **Fig. 2**, with the following aims:

**Phase 1 of 0.25 milliseconds:** To establish an equilibrated system of the StarD4 embedded in the membrane. Nine conformations from three representative states in phase 0 (*holo*-StarD4 in aqueous solution) were brought to the vicinity of the membranes using the *preparation protocol* described above. Parallel simulations were conducted on the three different types of membranes studied, utilizing 141 statistically independent replicas, amounting to a total simulation time of 249 microseconds.

StarD4 was considered embedded in membrane when the Ω1-H4 gate established substantial contact with the membrane: during the phase 1 simulation, the relative position of the Ω1-loop to the membrane surface converged to 2 stable states that contain >95% of the samples, either with no substantial contact with membrane (contact area <70Å^2^), or with multiple hydrophobic residues inserted into the membrane and the contact surface area >300Å^2^. Thus, we considered the Ω1-loop embedded in membrane when its membrane contact surface area is >200 Å^2^, and similarly, the β9-H4 motif is considered embedded when its membrane contact surface area is >150 Å^2^. StarD4 was considered embedded in membrane when both the Ω1 loop and the β9-H4 motif are membrane embedded simultaneously.

**Phase 2 of 0.34 milliseconds:** To explore StarD4 conformational changes and CHL translocation after membrane embedding. For each membrane type, 4 trajectories of embedded StarD4 from Phase 1 were replicated ×6 and continued in this Phase to expedite exploration of the cholesterol release pathway. These involved 72 independent runs (24/each for each membrane type), totaling 344 microseconds of cumulative simulation time.

**Phase 3 of 0.73 milliseconds:** To explore the cholesterol release pathways. A total of 48 conformations were selected from different states described in the text, which included the pre-release state, highly populated states, and states of PIP2-subtype-specfic binding (as illustrated in **Fig. 2** and **Fig. 7A,B**), as listed below:

- With bound PI(3,5)P_2_: 8 from the pre-release state; 5 from highly populated states; 11 from PIP2-subtype-specific states
- With bound PI(4,5)P_2_:8 from pre-release-like states; 8 from highly populated states; 4 from PIP2-subtype-specific states
- With bound PS: 4 from pre-release-like states.
- The conformations referred to in the text as “artificial pre-release state”, were manually constructed to mimic the pre-release state on the PI(3,5)P_2_-containing membrane, and repositioned on the membrane to adopt the pre-release orientation.

After an equilibration as described in the preparatory steps, adaptive sampling runs were carried out for a total of 312 replicas (6 replicas from each initial point), resulting in a total simulation time of 725 microseconds.

### Steered MD simulation and the calculation of potential mean force using umbrella sampling

***<u>The Steered MD simulation</u>*** was undertaken for accelerated sampling of the release paths of CHL from StarD4 in different conformational states. The collective variable guiding this process was the z-distance between the center of mass of cholesterol and the StarD4 α/β helix-grip structure (residues 70-76, 79-86, 103-108, 113-118, 131-140, 146-150, 170-175, 184-189, 211-220). Using a spring constant of 5 kcal/mol·Å^2^, the pulling phase of target was carried out with a very low pulling velocity of 0.0002Å/ps for 2ns, followed by an equilibration phase without target movement for 8ns. This 10ns cycle was continued within the first 200 ns. Subsequently, the velocity was increased to 0.0003Å/ps in the pulling phase, and the cycle is continued for another 200 ns. The total pulling lengths of 20Å was covered in 400ns, which moved the CHL from the binding site to the membrane. The 24 replicas of SMD generated from each state of interest (states 1,2,5,6,7 in **Fig. 7A,B**), were run according to this protocol, amounting to a total of 48 microseconds of SMD exploration.

***<u>The potential of mean force calculations</u>*** used the same collective variable – i.e. the z-distance between the cholesterol and the α/β helix-grip structure – as the reaction coordinate. Fifteen umbrella sampling windows were defined along the path, covering z=7 to 21Å with a spacing of 1Å. Each window utilized a harmonic potential function with a force constant of 1 kcal/mol·Å^2^. For each window, 15 independent runs of 40 ns were performed. The initial 10 ns of each run were considered equilibration and were excluded from subsequent analysis. The potential of mean force energy profiles were constructed using the Weighted Histogram Analysis Method (WHAM) algorithms (58).

### Production run simulations of the membrane embedded *apo*-StarD4

The simulations of the membrane embedded *apo*-StarD4 were carried out on the same computational platform and protocol type as the simulations of *holo*-StarD4. The simulation design is outlined in **Fig. 14**.

**Phases 1&2 of 0.21, 0.14 milliseconds:** The initial two phases mirror the design employed for *holo*-StarD4 simulations. In Phase 1 we establish an equilibrated membrane embedded StarD4-membrane, utilizing 81 statistically independent replicas amounting to 211 microseconds. This system is explored in Phase 2 to reveal the conformational changes after membrane embedding of StarD4 in the presence of membrane cholesterol, utilizing 36 statistically independent replicas amounting to 144 microseconds.

**Phase 3 of 0.19 milliseconds:** Probes the movement of membrane cholesterols near StarD4 with its Ω1-H4 gate open. The starting points were conformations resulting from the simulation of membrane embedded apo-StarD4 in phase 2 with a Ω1-H4-distance (defined as described in Supplementary Methods section 1) of >9Å. These conformations were replicated and continued. The protocol of Phase 3 includes 24 independent runs for the PI(3,5)P_2_-or PS-containing membrane, and 48 independent runs for the PI(4,5)P_2_-containing membrane, totaling 192 microseconds of cumulative simulation time.

**Phases 4-6 of 90, 72, 72 microseconds:** The protocols of the Phase 4-6 simulations explore the CHL uptake pathways from the membrane to the hydrophobic pocket. As shown in **Fig. 14**, the initial configurations for Phase 4 originated in Phase 3 and were selected if their CHL molecule had raised above the membrane surface, interacting with StarD4 at S208 or A201. Subsequently, the 6 trajectories in which cholesterol had reached highest above the membrane surface in Phase 4 simulations, were replicated as shown in **Fig. 14**, and continued in Phase 5. This process was repeated in going from Phase 5 to Phase 6. In the groups termed “ Δlipid-tail”, a specific alteration was made: the membrane lipid tails that occasionally also extended above the membrane surface were manually removed. This adjustment aimed to speed uptake convergence by preventing the cycles of competition for the hydrophobic pocket with the extracted CHL. The modification involved substituting the lipid with a conformation where both lipid tails resided within the membrane, while maintaining the interaction of the protein with the lipid head (**Suppl. Fig 12**). The initial seeds that underwent this manual modification were re-equilibrated before the production run, with constrains applied on the protein backbone and the target cholesterol (following the preparation protocol described for the of StarD4-membrane complex).

### Rare Event Detection analysis

As described in the main text, we used the Rare Event Detection (RED) protocol detailed in our publications (15,23) to find the structural motifs that move together to establish new conformations and PIP2-protein interaction modes that dominate during specific time segments of the trajectory. Briefly, the protocol implements the Non-negative Matrix Factorization (NMF) module in the Scikit-Learn Python package (59) to identify rare structural rearrangement events in the molecular dynamics (MD) trajectory data by decomposing a trajectory into structural motifs and detect those undergoing simultaneous changes that dominate the conformation of the protein for specific periods along the trajectory. The input to the NMF algorithm is a contact matrix constructed from the entire trajectory, which therefore contains the information about the evolving protein structure. In the NMF algorithm this matrix is further decomposed into a spatial array (W) and a temporal array (H) as described in (23), with the former defining “components” (which are sets of conformational features that evolve concurrently in the trajectory), and the latter depicting the time-ordered contribution of each component along the trajectory (15,23).

Here, the contact matrix contains residue-residue interactions and residue-pair/PIP2 interactions calculated in each of the 10,000 frames sampled along the 4-microseconds simulation of RT1. For residue-residue contacts matrix, a contact is marked as 1 if any atom from one residue is within 3.5Å of that of another residue, resulting in a 201^2^×10000 matrix. In the residue-pair-PIP2 contact matrix, a diagonal entity is marked as 1 if any atom from one residue is within 3.5Å of a PIP2, while an off-diagonal entity is marked as 1 if both residues are within 3.5Å of a shared PIP2, resulting in another 201^2^×10000 matrix for 10000 frames. These two matrices of size 40401×10000 are concatenated and result in an 80802×10000 matrix representing the residue-pair/PIP2 interactions for every frame. To capture dynamic information, contacts that remain unchanged by these criteria throughout the simulation are excluded. The resulting contact map contains 2171 columns of pairs of residues that alter their contact states during the simulation, and has 10,000 rows, recorded at intervals of 0.4ns. Smoothing (see (15,23)) is applied across the time dimension using a sliding window of 20 nanoseconds to refine the array.

The NMF algorithm then decomposes this contact matrix into the W and H components that place structural states in spatial and temporal matrices. In this study, the choice of six components (see (15,23)) reveals the transition of conformational states in RT1. The spatial arrays are normalized, allowing direct comparison between components, and the differences between components help identify structure-differentiating contact pairs (SDCPs). These SDCPs, highlighted in the transition from one component to another as described in (15), denote pairs that contribute significantly to structural variations, which yields a comprehensive identification of rare events within the trajectory data.

### tICA - the time-structure based independent component analysis used for dimensionality reduction and identification of metastable states

The dimensionality reduction approach we used for the analysis of the conformational space as described in the main text was carried out with the well-known tICA method (60–64). The tICA is constructed by projecting the simulation data onto low dimensionality (usually 2D) spaces (see **Figs. 7C, 9B, 17A**) defined by several collective variables (CVs) that quantify the dynamics of interest. The CVs we used are described in detail in Supplementary Methods. The tICA transformation of the simulation data identifies the slowest reaction coordinates in the conformational space by solving for the generalized eigenvectors and eigenvalues of the time-lagged covariance matrix:

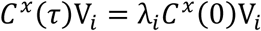

where *C*^*x*^(τ) is the time-lagged covariance matrix (*C* ^*x*^(τ) = 〈*X*(*t*)*X*^*T*^(*t* + τ)〉_*t*_) between collective variables, *C* ^*x*^(τ) is the regular covariance matrix (*C* ^*x*^(0) = 〈*X*(*t*)*X*^*T*^(*t*)〉_*t*_), *X*(*t*) is the data vector at time *t*, τ is the lag-time of the covariance matrix, V_*i*_ is the i^th^ tICA eigenvector, and λ_*i*_ is the i^th^ tICA eigenvalue of the covariance matrix. In this study, the trajectory was sampled at a stride of 0.4 ns/frame, and the lag-time τ for constructing the covariance matrix was set to 20 ns (50 frames). This choice allowed the exclusion of fluctuations faster than 20 ns, which were considered not relevant to the slow processes in the system. The first 200 ns in every independent run were discarded to reduce equilibration artifacts before analysis. This dimensionality reduction technique facilitated a focused exploration of the slow processes and relevant reaction coordinates in the simulated dynamics as described below.

#### Mapping the cholesterol uptake pathway and assignment of macrostates in the tICA space

This analysis focused on the Phases 4-6 of the simulation of *apo*-StarD4 membrane complex (**Fig. 14**) in which adaptive sampling was used to simulate the progress of membrane cholesterol into the hydrophobic pocket of StarD4. Key collective variables were recorded to assess the position of the cholesterol (as defined supplementary methods) in the 234 simulation trajectories conducted in these three phases. The CHL position data included the height reached above the membrane surface, the distance to the Ω1-H4 gate, and its distance to the hydrophilic residues that served as a ladder into the hydrophobic pocket of StarD4. These data were used as CVs to define the tICA transformation. To identify metastable states in the cholesterol uptake process, the 2D tICA space was discretized into 100 microstates using K-means clustering. Subsequently, these microstates were further clustered into 10 macrostates based on their kinetic similarities, determined from the transitional rate between microstates. This clustering process was achieved using the Robust Perron Cluster Analysis (PCCA+) algorithm (65).

### Construction of the PIP2 Interaction Mode Landscape (IML) and assignment of PIP2 binding modes

The IML for PIP2 binding modes to the StarD4 (**Fig. 9C,D**) represents the positions the PIP2 molecules that had established stable interactions with the basic residues. The criterion for considering a PIP2 molecule to be binding to the upper basic residue patch is that it remains for at least 10 ns within a distance of <4.5Å from at least two of residues K49, K50, K52, R97, S215, R218, K219, R222 (“upper basic group”). The position of such a PIP2 molecule was recorded from the first frame where it formed an interaction with upper-group residue until the last frame of its binding in the same trajectory. If unbinding and later rebinding occur in the same trajectory for the same PIP2, its position is continuously recorded.

This analysis was performed on the trajectories of Phases 2&3 (**Figs. 2&14)** of both *apo*- and *holo*-StarD4 embedded in either PI(3,5)P_2_-or PI(4,5)P_2_-containing membranes. A total of 288 trajectories of StarD4-membrane complexes were included in this analysis. From this set of simulations, a total of 872 PIP2s that bind to upper group residues were recorded. (Note that multiple PIP2s can bind simultaneously to each StarD4). The positions of bound PIP2 were defined using 5 collective variables (CVs), each measuring the distance between the PIP2 anionic atoms and the closest basic atoms in the K52/R97/S215/R218/K219 patch, resulting in 872 position-time matrices representing the evolution of the PIP2-StarD4 interactions over time. These CVs served in the construction of the tICA dimensionality reduction analysis as described above.

The first two tICA vectors composed of the 5 CVs were found to describe 85% of the total dynamics of the PIP2 interacting with the upper basic patch. To facilitate the comparison of population densities and preferences between PI(3,5)P_2_ and PI(4,5)P_2_ binding modes, the 2D tICA space was discretized into 100 microstates using K-means clustering, and segregated into 9 macrostates based on their kinetics similarities (see above). The assignment of macrostates provides a valuable mapping tool that facilitates direct comparison between PI(3,5)P_2_ and PI(4,5)P_2_ population densities in the states they represent. These macrostates are herein designated as the definition of the binding modes 0 to 8 of PIP2. Every timestep in the 872 position-time matrices of PIP2 is assigned to its macrostate, representing the binding mode of a certain lipid at the given time. Then, these 872 trajectories of PIP2 are mapped back to the original 288 trajectories of StarD4-membrane system where multiple PIP2 may binding simultaneously on one StarD4, resulting a “matrix of binding mode over time” which quantifies the number of lipid(s) in each lipid binding mode on a StarD4 at any given time. This matrix provides a summary of the occurrence and distribution of different binding modes on StarD4 throughout the simulation. We note, however, that because the kinetics similarities of the microstates that served in the definition of the 9 macrostates were derived from a mixture of PI(3,5)P_2_ and PI(4,5)P_2_ trajectories, the macrostate assignment from this map may contain microstates from both, and cannot be used to study the dynamics of the protein in the individual environments.

### The Deep Neural Network

#### Input data structure

The DNNs described in the main text were trained to study the difference between the StarD4 interacting with membranes containing PI(3,5)P_2_ vs membranes containing PI(4,5)P_2_. The input dataset for the DNNs consisted of the ensemble of trajectories containing ∼46,000 StarD4 protein structures, without the information regarding the ligand CHL or the membrane lipids. The protein structures were sampled from the simulations of StarD4-membrane systems including both the membrane-embedding phase and the membrane-embedded phase, at 8ns time-intervals. The first 160ns of every trajectory were considered equilibration time and excluded from the data set. The simulations used for the DNN dataset sampling are listed in **Suppl. Table 6**.

The trajectory data were subjected first to a *debiasing protocol* that consists of random scrambling of the position and orientation of the protein to eliminate their influence on the decision-making process of the DNNs. With the debiased data, the DNNs rely solely on intramolecular protein conformational changes to predict the labeled lipid type. To further augment the data, each protein structure is scrambled into three different random orientations, resulting in approximately 14,000 samples. This dataset was divided into training, validation, and test sets in a ratio of 56:24:20, respectively, keeping track of the three samples derived from the orientation scrambling of the same structure.

The analysis of the molecular dynamics trajectories employed image-recognition convolutional neural networks in a pipeline developed in our lab (30) in which the coordinates of the 3D protein structures are transformed into 2D pixel representations that retain their 3D information. Briefly, to convert a structure into a picture, the (X,Y,Z) coordinates of each atom are normalized and converted into a set of (R,G,B) values of a color point. These color points are then arranged in sequential order according to the protein sequence from top left to bottom right in a 2D input picture readable by a convolutional neural network. In the case of DNN based on the full StarD4 protein, the first 3192 atoms of the residues in the 24 to 223 range are converted into a picture representation with a size of (56,57,3), where the 56 rows and 57 columns collectively contain 3192 pixels representing each atom, and the third dimension contains 3 (R,G,B) colors representing the X,Y,Z of the atom. In the case of DNN based on the salient residues, the first 400 atoms in the residues 46 51 52 53 58 62 63 67 123 124 130 160 165 167 175 180 181 188 189 192 194 195 196 are converted into a picture representation with a size of (20,20,3). This method allows for the efficient application of image-recognition convolutional neural networks to the analysis of molecular dynamics trajectories.

#### Building the DNN

The recognition of the protein structures in their image representation is carried out with custom Dense Convolutional Networks in Keras with a Tensorflow based on a pre-existing implementation (66). The DNNs utilize Dense Blocks, where feature-maps from all the preceding layers are used as inputs for the subsequent layer. In the case of the DNN based on full protein atoms, the DNNs for StarD4 take images of size (56,57,3) as inputs, starting with a convolutional layer with a stride of 2, followed by 3 dense blocks containing 6, 12, 24 convolutional layers respectively. When only the residues identified as the salient contributing features by the DNN (see above) were used as input to another DNN analysis run as described in the main text, the input was constructed as images of size (20,20,3), starting with a convolutional layer with a stride of 1, followed by 3 dense blocks containing 6, 12, 24 convolutional layers. The DNNs utilize a growth rate of 48 filters per layer, 96 initial filters, a reduction ratio of 0.5 in the transition between Dense Blocks. A dropout rate of 0.2 is applied in every convolutional layer during training.

#### Visualization of the salient contributing features in DNN

The saliency analyses are conducted among the top 1000 samples, where the DNN reported the highest confidences in correctly identifying the PIP2 subtype. The contribution of each atom to the decision-making by DNN was evaluated using the visual saliency package provided by keras-vis (67) to calculate the gradient of the output label over the input features with guided back-propagation. Averaging the contributions of each atom over the 1000 samples results in a saliency map of the same shape as the input sample size.

### Quantifying the allosteric mechanism using N-body information theory (NbIT)

We used the information theory-based NbIT analysis methodology (24,25) to calculate the terms that quantify the information sharing between structural motifs involved in the allosteric communication and pathways. The structural motifs involved in the analysis are defined below, at the end of this section.

A key term defined in detail (24,25) and used as described in recent publications (33–36) is the *Coordination Information* (CI) between motifs that is calculated from the entropy in each motif *H*(*X*) as described in detail in (24) where

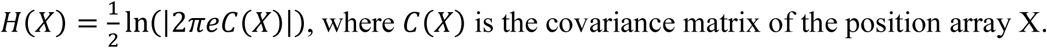

Briefly summarized, the *Coordination Information calculation* starts from the total information shared by structural motifs within a set, termed *Total Correlation* (TC), that is calculated from differential entropy terms as:

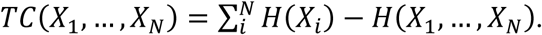

A *Conditional Total Correlation*, indicating the remaining information within a set of motifs {*X*_1_,…, *X*_*N*_} given the dynamics of another motif *X*_*m*_, is then obtained as:

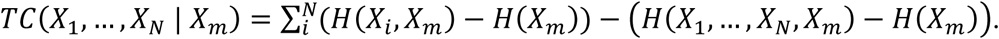

By relating the *Total Correlation* and the *Conditional Total Correlation*, we obtain the *Coordination Information* (CI) that quantifies the portion of information shared in the structural motifs described in the main text as *receptors* {*X*_1_,…, *X*_*N*_}, that is also shared with *X*_*m*_ motif that functions as the *transmitter*:

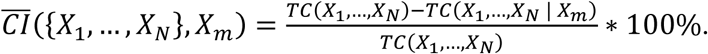

In **Tables 2&3** and the **Suppl. Table 5**, the *Total Correlation* within each motif is displayed along the diagonal. The off-diagonal elements quantify the *Normalized Coordination Information*. In these Tables, the structural motifs shown at the top of each column are acting as the *Transmitter* and those on the left side of each row indicate the coordinated *Receiver*.

The same information is used to identify the allosteric channels that connect distant motifs, identified by calculating a term named *Normalized Mutual Coordination Information* (*NMCI*) (see (24)), which measures the portion of coordination information shared between the *receiver* {*X*_1_,…, *X*_*N*_} and the *transmitter* motifs *X*_*m*_ that is also shared with another structural element *X*_*n*_:

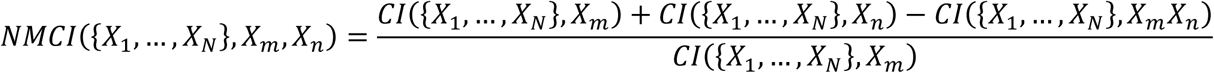

#### Definition of the structural motifs included in the NbIT analysis

**CHL sites**: S136 S147 W171 R92 Y117; **β1**: R46 V47 A48 K49 K50 V51; **β2&β3**: R58 K59 P60 Y67 L68 Y69; **β2β3 loop**: E62 E63 F64 N65; **H4-N-terminal (H4-Nt)**: Q199 S200 A201 D203 T204 A207; **Ω1 loop**: L124 N125 I126; **Ω4 loop**: D192 R194 G195; **PIP2 binding mode (Pbm)**: K52, R97, S215, K219; **Water influx**: N166, C169, I189, T191; **β8β9 Loop**: K177 D178 S179 P180; **β4-Nt**: H107 F108 E109; **H4-Ct**: G220 L221 K223 A224.

## Supporting information

Supplementary Information- Complete

## ACKNOWLEDGEMENTS

We gratefully acknowledge helpful discussions with Drs. Ambrose Plante and Derek M. Shore, and with Prof. George Khelashvili. HX gratefully acknowledges their helpful guidance regarding the principles, applicability, implementation, and interpretation of the RED, NbIT and TPT analysis. We are grateful for very helpful discussions with Prof. Frederick R. Maxfield and experimental data from his lab members. Support from the 1923 Foundation for the project “*How Needed Molecular Precision is Achieved for Addressing, Pickup, and Delivery in the Trafficking of Cholesterol Among Cell Membranes*” is gratefully acknowledged. The computational resources and technical help at the Center for Computational Innovations (CCI) at the Rensselaer Polytechnic Institute (RPI), and the efficient and sustained access to the AiMOS supercomputer at CCI generously awarded through the COVID-19 High Performance Computing Consortium, are gratefully acknowledged. We are grateful for the computational resources under Project BIP109 at the Oak Ridge Leadership Computing Facility, which is a DOE Office of Science User Facility supported under Contract DE-AC05-00OR22725, and for the in-house computational resources of the David A. Cofrin Center for Biomedical Information in the Institute for Computational Biomedicine at Weill Cornell Medical College.

## Supplementary Information

### A. SUPPLEMENTARY FIGURES

**Supplementary Figure 1.**
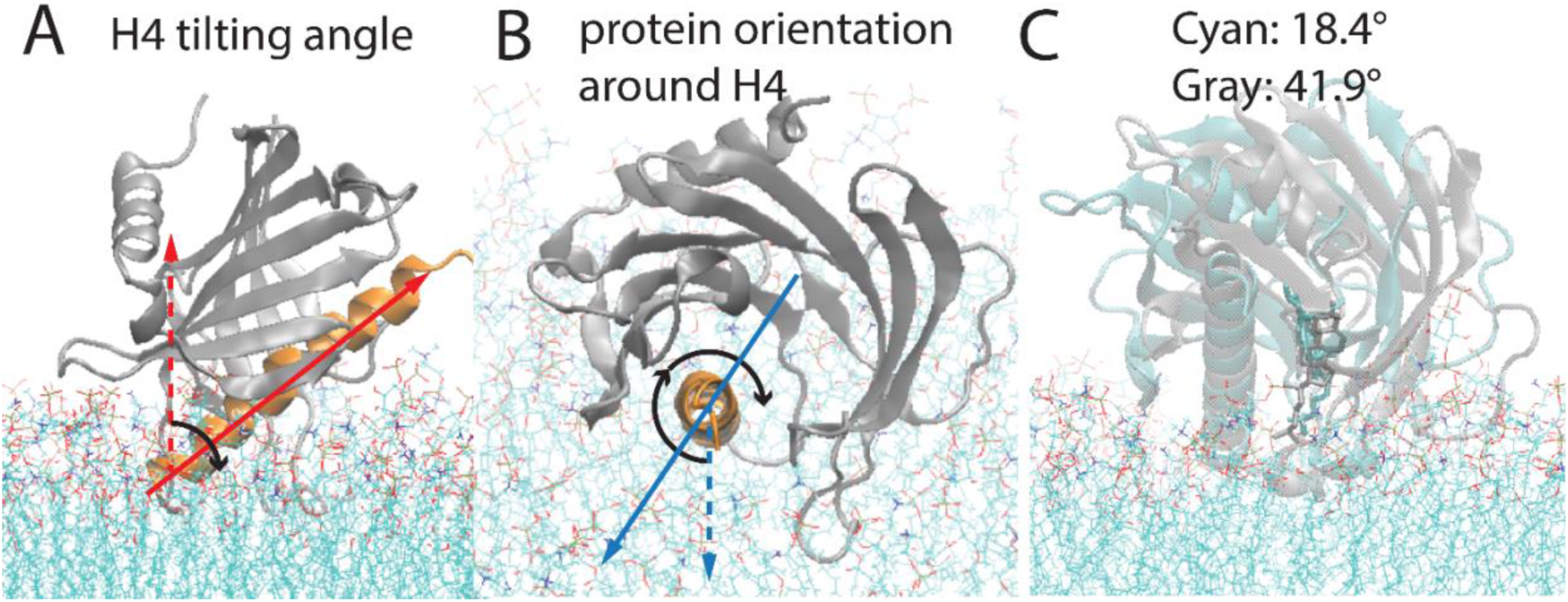
The orientation of StarD4 in relation to the membrane bilayers. **(A)** The definition of the H4 tilting angle: the angle between the “membrane norm” (red dashed line) and the “Helix4 axis” (red solid line, the first principal component of the Cα atoms of H4). **(B)** The definition of the protein orientation around H4: The angle is measured on the plane perpendicular to the Helix4 axis. The “membrane norm” (blue dashed line) and the “vector from the center of mass of StarD4 to the center of mass of H4” are the projected onto the plane perpendicular to the Helix4 axis, and the CV is defined by the angle between the two projections. **(C)** The representative conformations of the cholesterol-StarD4 complex at the protein orientation around H4 equals 18.4° (cyan) and equals 41.9° (gray).

**Supplementary Figure 2.**
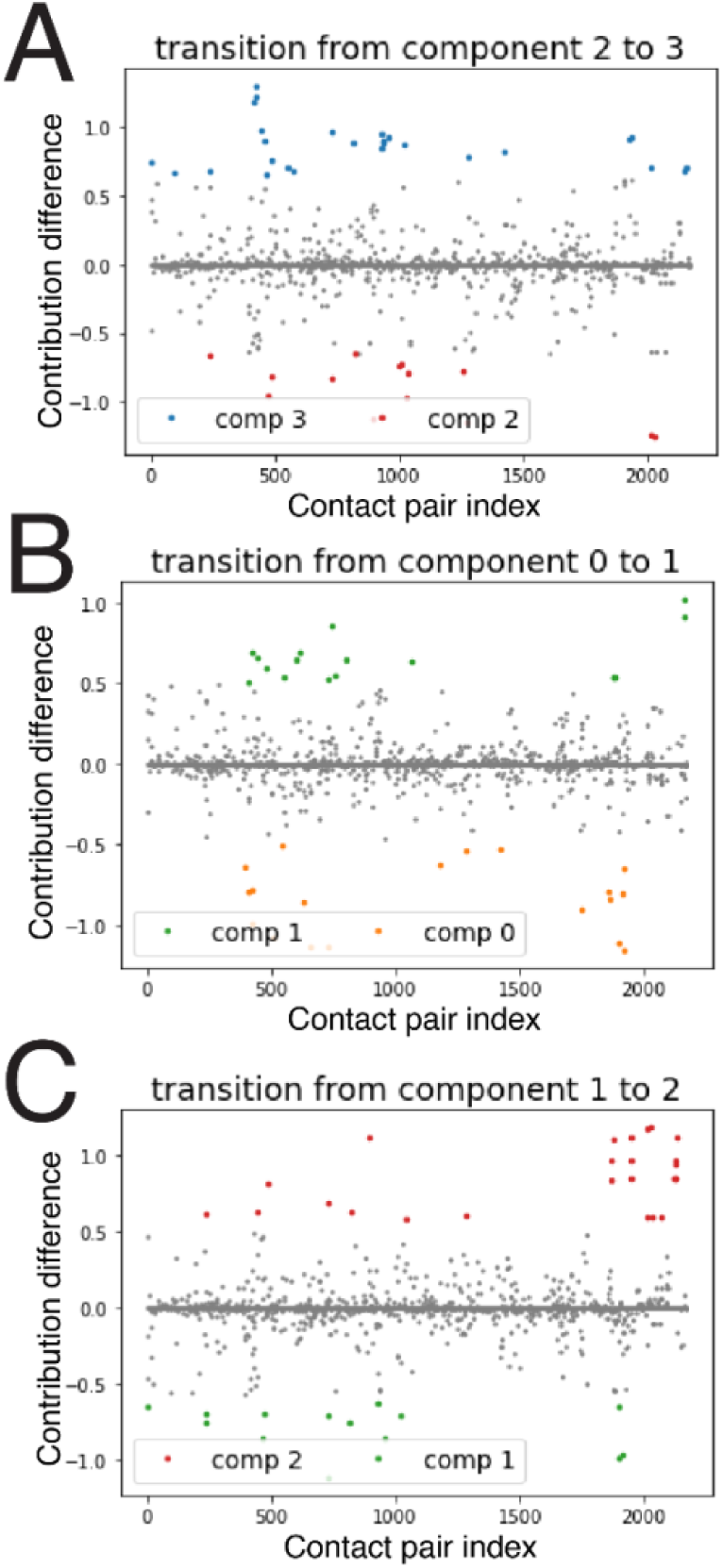
The structure-differentiating contact pairs (SDCPs) are revealed through the comparison of the normalized spatial components. **(A) SDCPs** for the transitioning of component 2→3; **(B)** for the transitioning of component 0→1; **(C)** for the transitioning of component 1→2. On the X axis are the 2171 data points representing the residue pairs. The Y axis shows the difference in the contribution of a residue pair to the conformational state in the transition between the two components. A positive contribution difference reflects the creation of a contact in this pair during the transition. A negative contribution difference reflects the breaking of the pair contact during the transition.

**Supplementary Figure 3.**
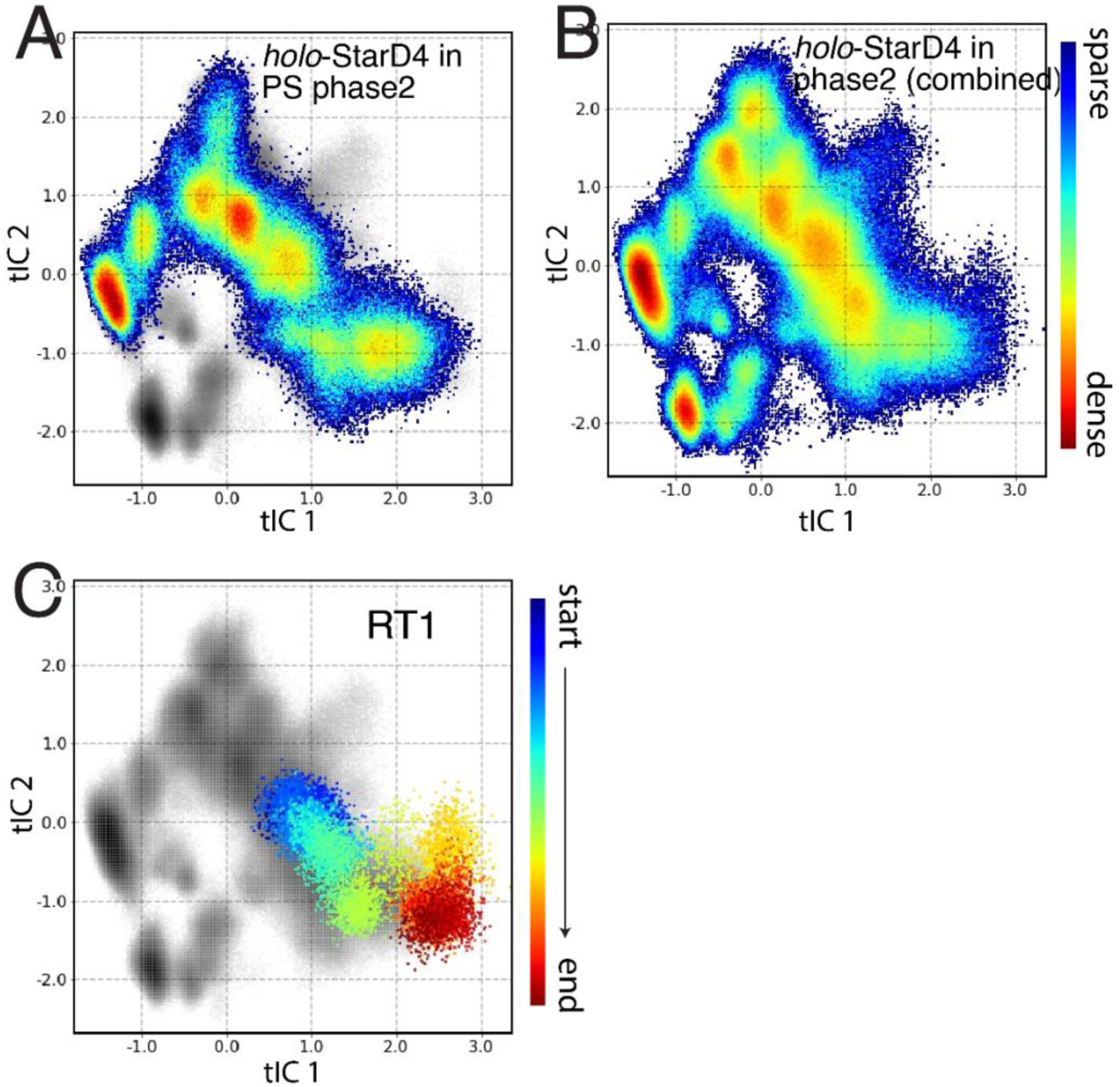
Population density maps of the conformational space of *holo*-StarD4. **(A) *holo*-StarD4** embedded on PS-containing membranes; **(B)** The combined population density maps for ***holo*-StarD4** on all three types of membranes. The color scale for the representation of population densities is shown on the right; **(C)** The time-evolution of the cholesterol releasing trajectory RT1 projected onto the combined conformational space. The trajectory evolution is color-coded from the starting point in blue, to the end point in red. The gray scale in the background represents the combined population density as in **(B)**.

**Supplementary Figure 4.**
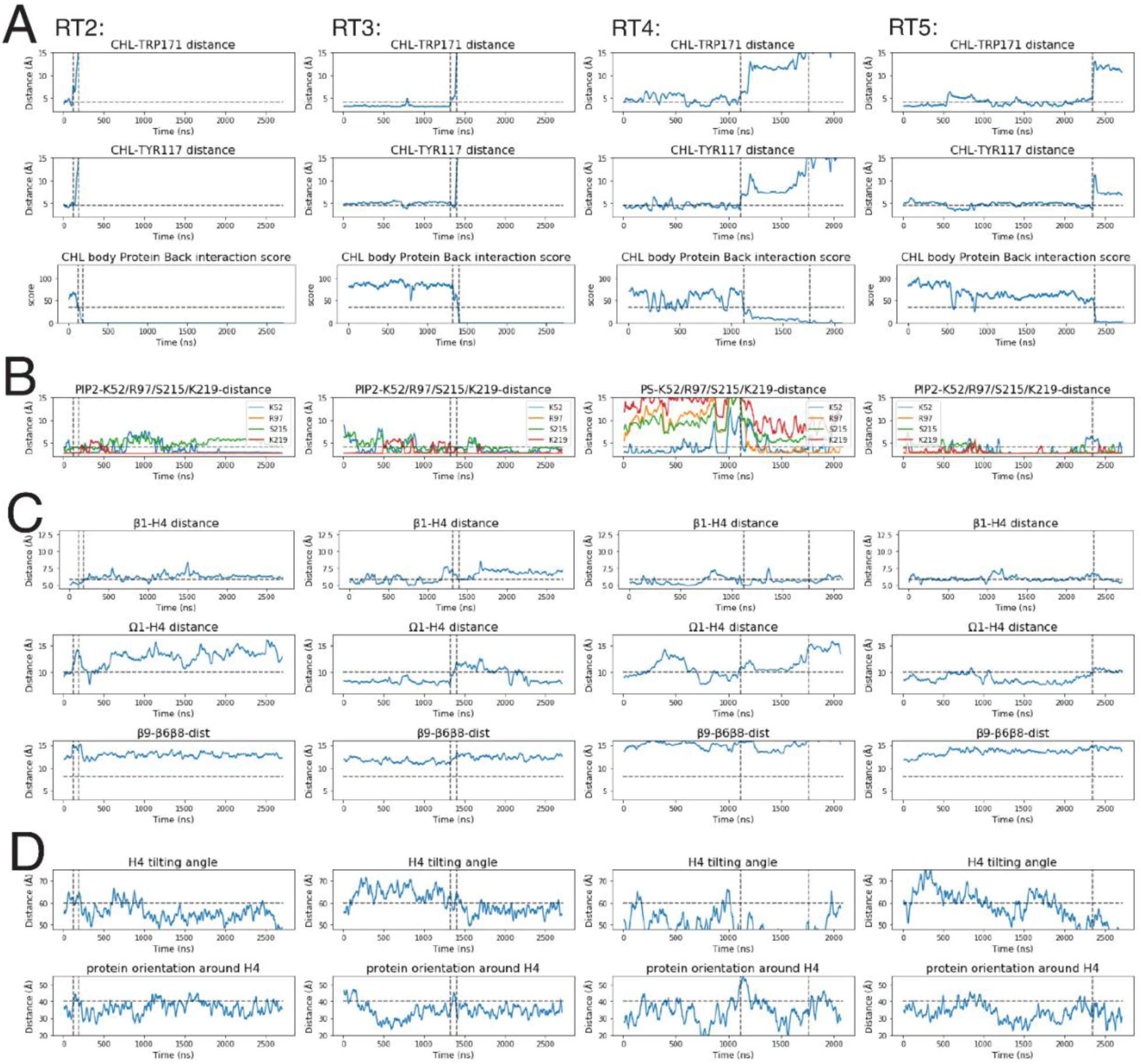
The conformational changes related to the pre-release state in the Release Trajectories 2-5. Data for RT2 to RT5 are shown in the columns from left to right. The rows depict: **(A)** The time-evolution of the CHL position relative to the binding sites in the pocket, and of the interaction between the CHL hydrocarbon ring and residues N166, C169, I189, T191 on β8 and β9 at the back of StarD4. **(B)** The time-evolution of the binding of anionic lipid in the cross-H4-binding mode. **(C)** The time-evolution of the conformational changes of StarD4, including the repositioning of H4 away from Ω1 towards β1, and the opening of Ω4 from β8&6. **(D)** The time-evolution of the orientation of StarD4 in relation to the membrane. The vertical dashed lines indicate timing of the CHL release process as shown in **(A)**. The horizontal dashed lines are auxiliary lines that indicate the features of the pre-release state.

**Supplementary Figure 5.**
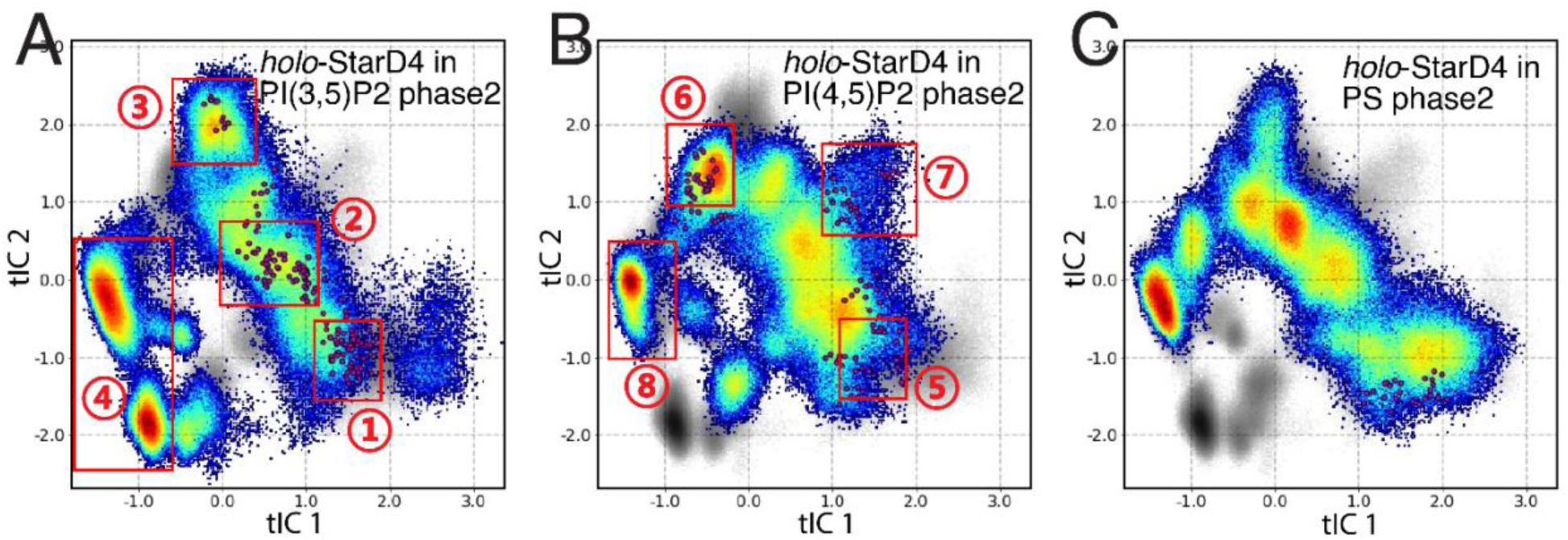
The initial seeds of the adaptive sampling in Phase3. The initial seeds of Phase3 simulations are indicated by red spots located at different conformational states on the tICA space of *holo*-StarD4 on the PI(3,5)P_2_-containing membrane **(in A)**, on the PI(4,5)P_2_-containing membrane **(in B)**, or on the PS-containing membrane **(in C)**.

**Supplementary Figure 6.**
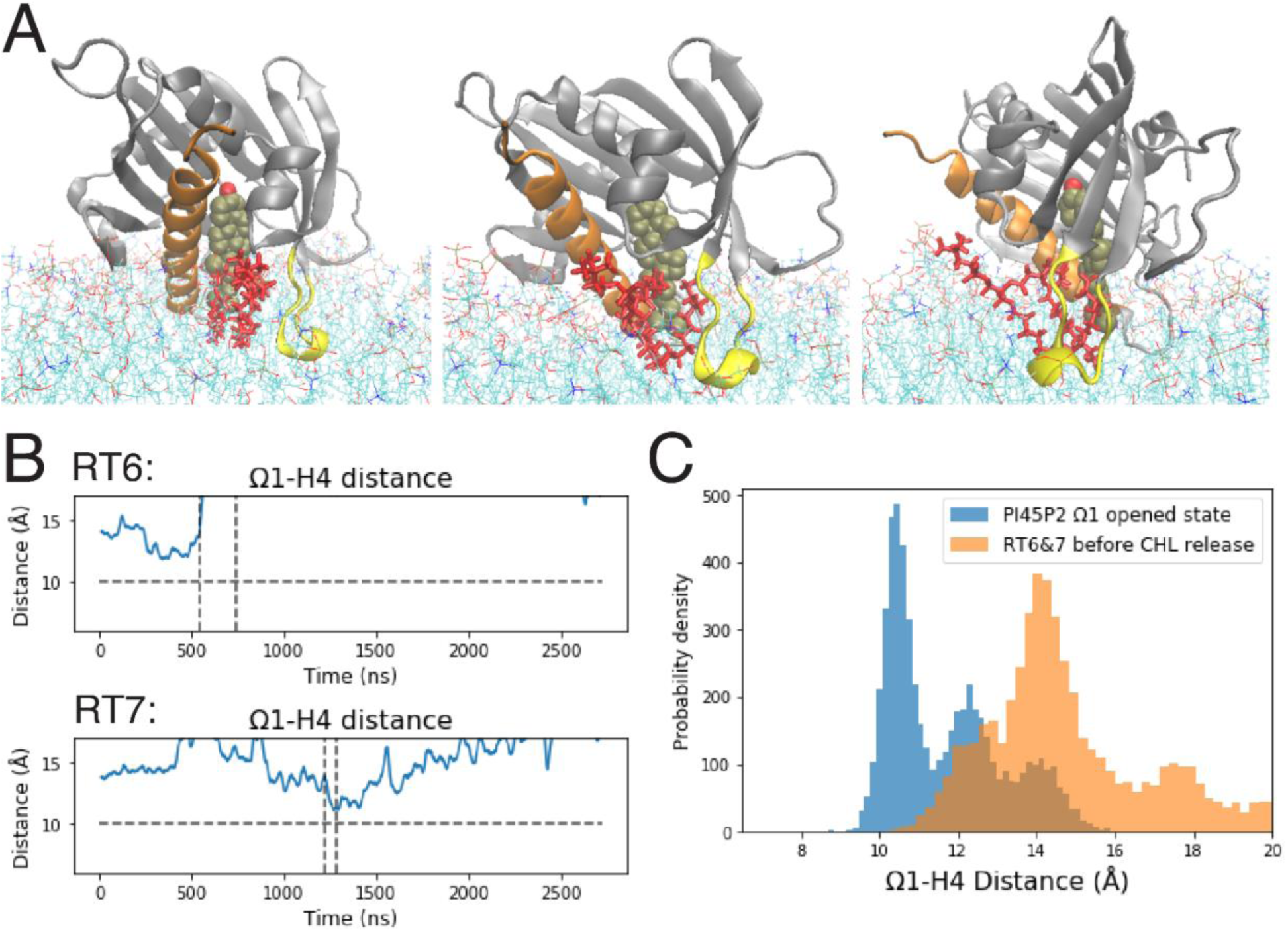
The initial structures of the (**RT6**, **RT7**) set feature an Ω1-open configuration on PI(4,5)P_2_-containing membranes (Fig. 7B state 7), characterized by the wide opening of the Ω1-H4 gate (Fig. 7D, 4^th^ column), and they shared with the pre-release state a similar pattern of a widely opened β8-Ω4 corridor (Fig. 7D, 3^rd^ column), but without the characteristic β1-H4 interaction (Fig. 7D, 6^th^ column). These features are likely related to the membrane lipid tail that had inserted between the Ω1 loop and the CHL in the initial frames (**Suppl.** Fig. 6A), which resulted in a persistent and wide opening of the Ω1-H4 gate (centered at CV “Ω1-H4-dist”=14Å). Notably, this feature is not observed in the pre-release state shared among RT1-RT5 (Fig. 7D, 4^th^ column, **Suppl. Fig. 6B,C**) **The CHL release path initiated from the high energy state with a widely opened Ω1-H4 gate due to an inserted lipid tail. (A)** The membrane lipid inside the Ω1-H4 gate of *holo*-StarD4 depicted from 3 different viewing angles 45° apart from the front to the right. The StarD4 is rendered in gray cartoon with H4 in orange and Ω1 in yellow. The cholesterol in shown in VDW, and the gate-penetrating lipid is shown in red licorice. **(B)** The time-evolution of the widely-opened the Ω1-H4 gate. The horizontal dashed line is an auxiliary line that indicates the threshold of the opening of Ω1-H4 gate. The vertical dashed lines indicate the CHL release process. **(C)** Comparison of the probability density histograms of the Ω1-H4 distance in the Ω1-H4 opened state (blue) and the release trajectories 6 and 7 (orange).

**Supplementary Figure 7.**
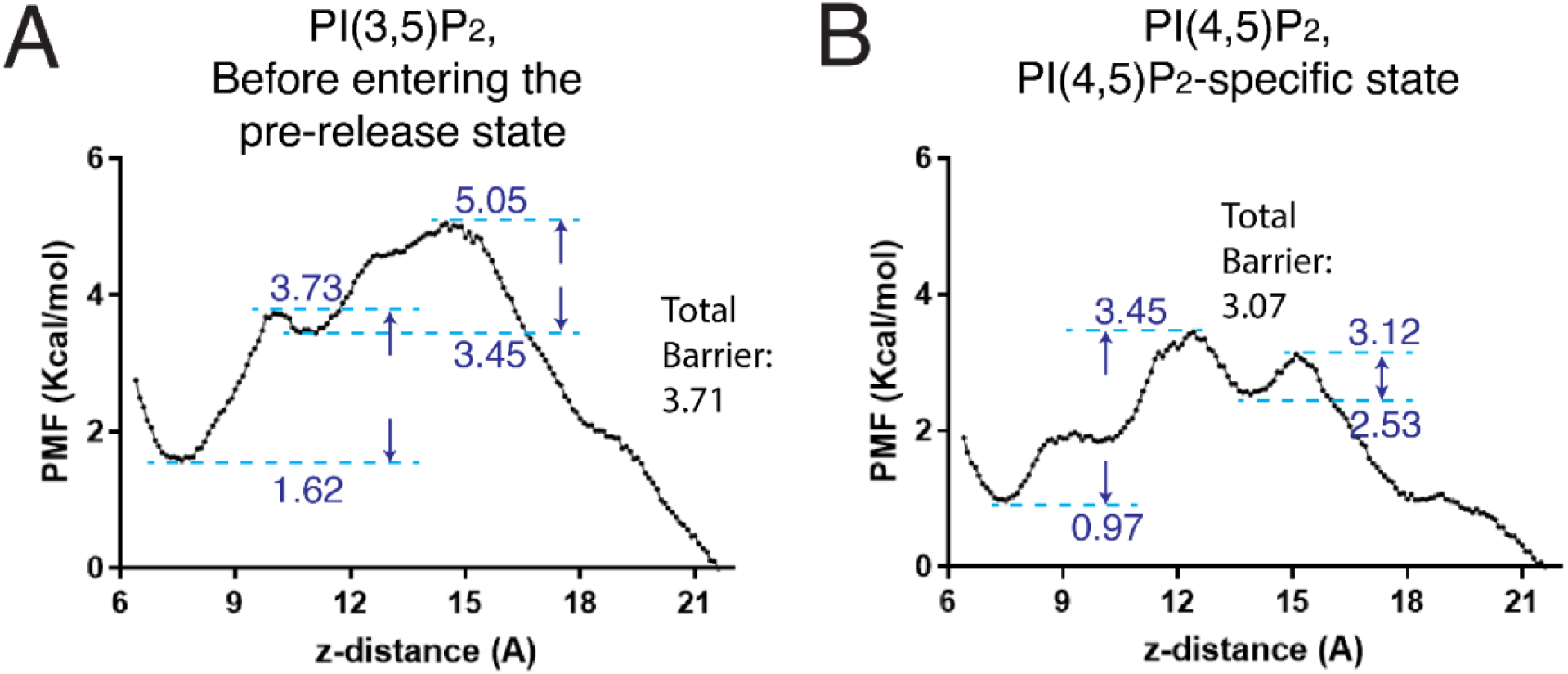
**The energy barrier along the reaction coordinate of CHL release from StarD4**, sampled with steered MD simulation and evaluated with umbrella sampling. The barrier is represented in terms of the potential of mean force. The reaction coordinate is the z-distance between the CHL and StarD4. The initial point is the common state on the PI(3,5)P_2_-containing membrane **(A**, state 2 in Fig. 6), and the PI(4,5)P_2_-specific state **(B**, state 6 in Fig. 6). Local maxima and minima are labeled along the reaction path, and the total barrier is shown in the sidenote.

**Supplementary Figure 8.**
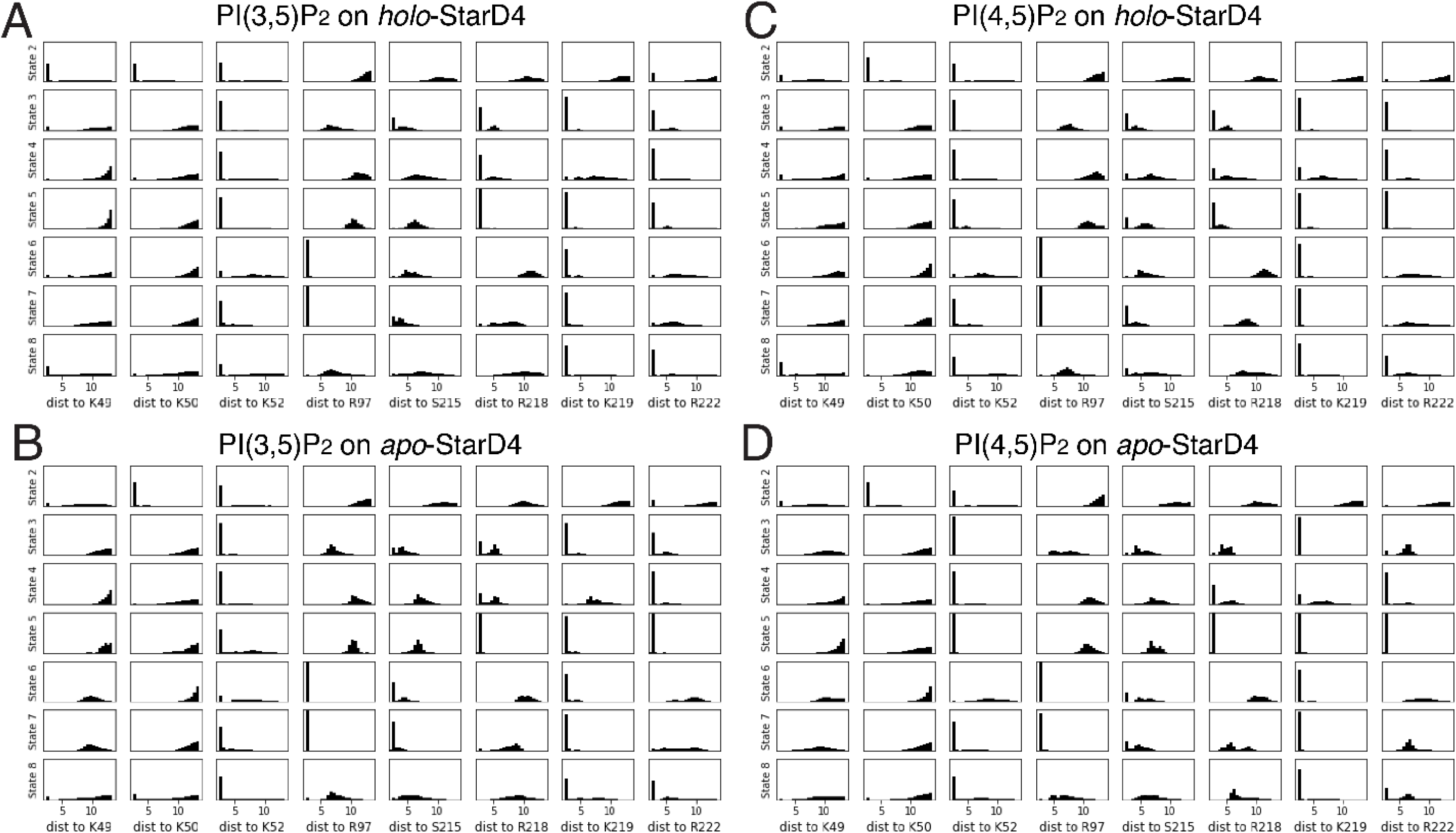
Characteristics of the PIP_2_ binding modes (modes 2 to 8 from top to bottom). The probability density histograms of the characteristic CV values show (from left to right) the interactions of PIP_2_ with the key residues K49, K50, K52, R97, S215, R218, K219, R222. **(A-C)** Show the characteristics of PIP_2_ binding modes in 4 scenarios: **(A)** PI(3,5)P_2_ binding on *holo*-StarD4, **(B)** PI(3,5)P_2_ binding on *apo*-StarD4, **(C)** PI(4,5)P_2_ binding on *holo*-StarD4, and **(D)** PI(4,5)P_2_ binding on *apo*-StarD4.

**Supplementary Figure 9.**
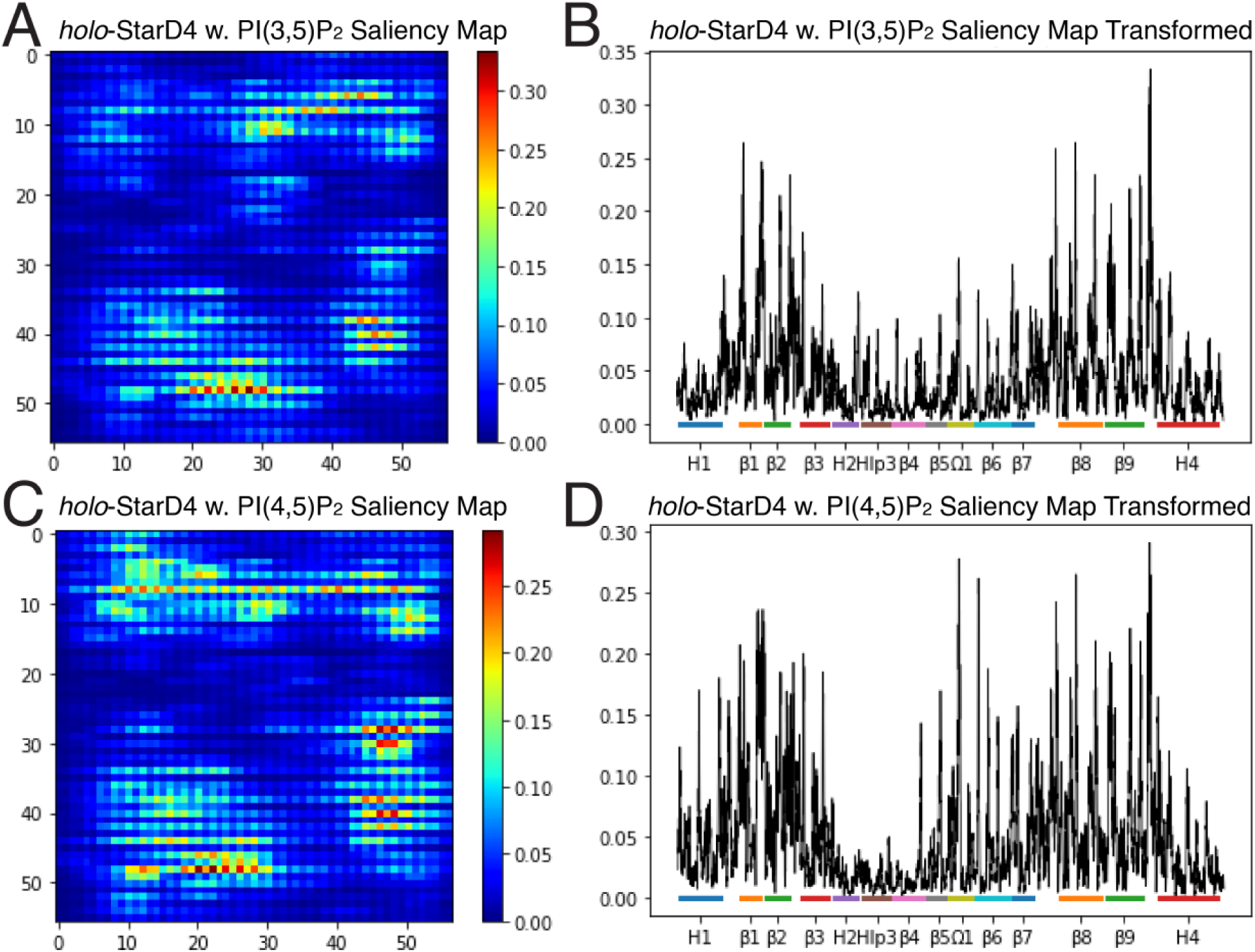
Identification of the molecular determinants of the decision of DNN. The attention maps of the atoms contributing to the decision of DNN in **(A)** the identification of StarD4 with PI(3,5)P_2_-binding, and **(C)** the identification of StarD4 with PI(4,5)P_2_-binding. Each pixel in the picture represents an atom, and the color code indicates its contribution to the decision, according to the color scale at the right. In **(B,D)** the attention maps shown in (A,C) are reshaped to list all the atoms on the x-axis, while the y-axis indicates their respective contribution to the decision. The color bar and tick marks on the x-axis identify the motifs to which the atom belongs.

**Supplementary Figure 10.**
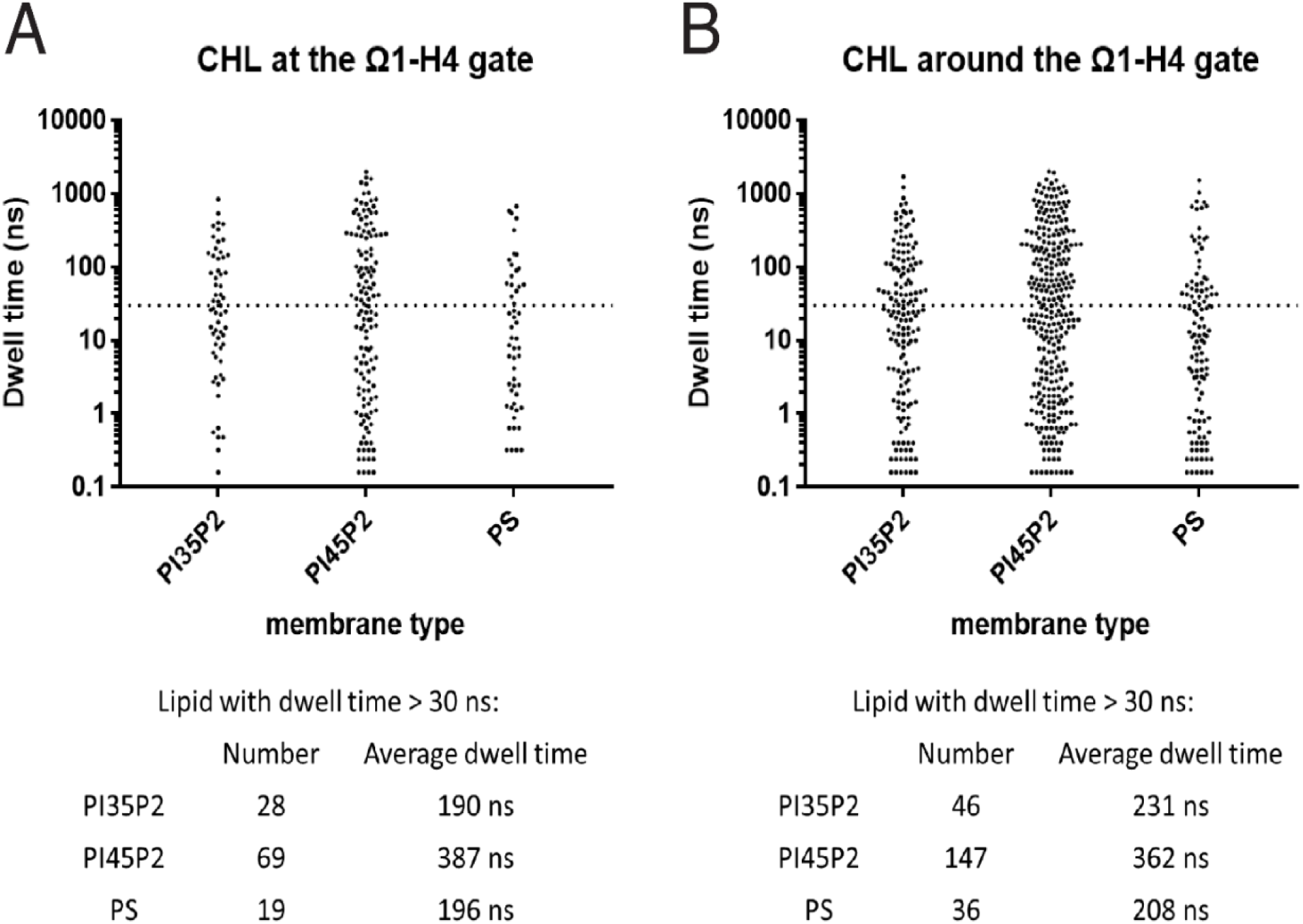
Scatter plots of membrane CHL dwell time adjacent to StarD4, compared between the StarD4 embedded on PI(3,5)P_2_-, PI(4,5)P_2_- and PS-containing membrane. Each dot indicates a CHL molecule, and the y-axis shows its dwell time at the Ω1-H4 gate **(A)** or adjacent to the Ω1-H4 gate **(B)**. The definition of the criteria is shown in the Supplementary Materials. The dashed line marks the 30 ns dwell time threshold. The average dwell time of CHLs with dwell time >30 ns is listed below the figure.

**Supplementary Figure 11.**
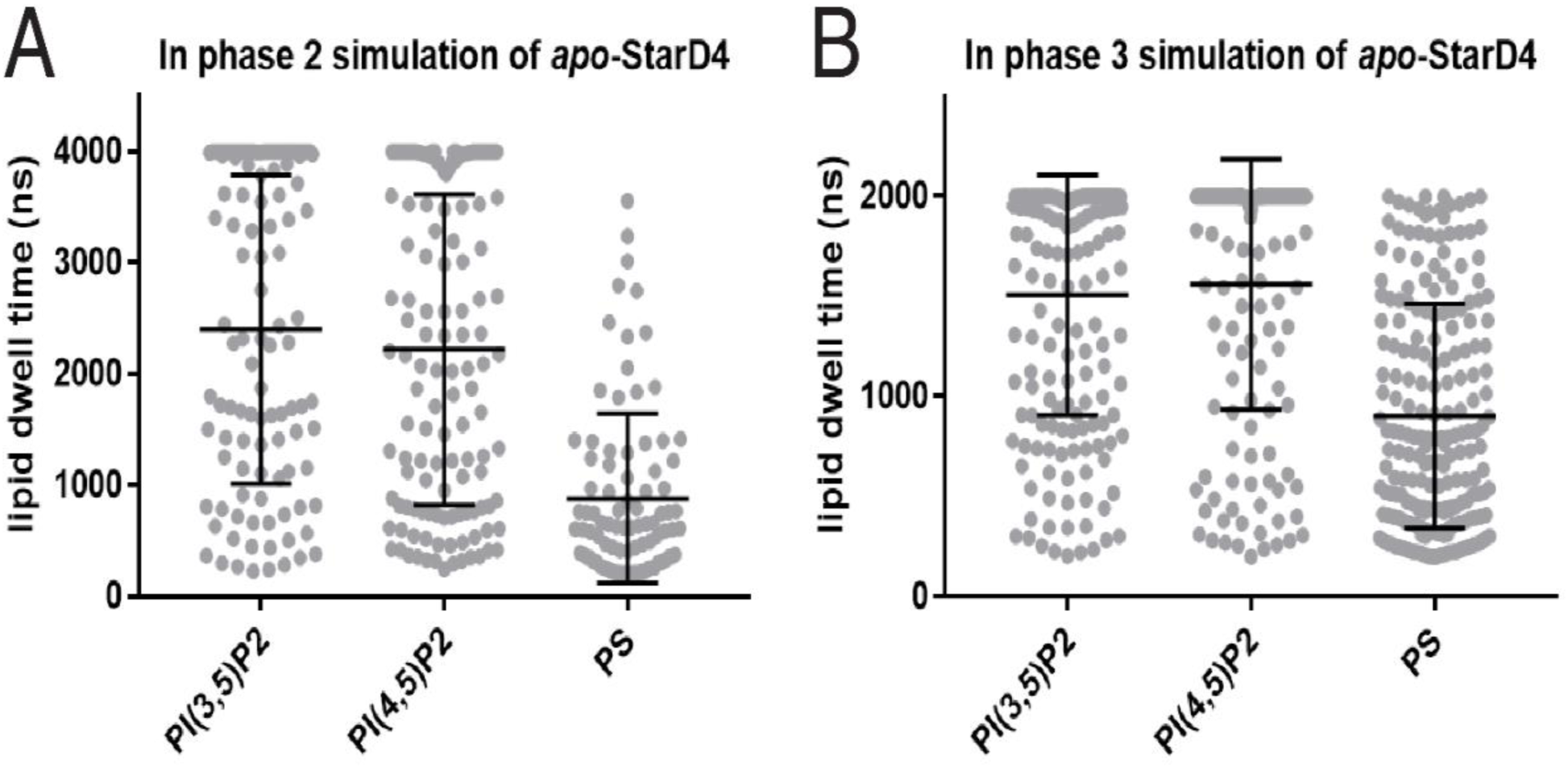
Scatter plots of the dwell time of anionic lipids interacting with StarD4 in membranes containing PI(3,5)P_2_, PI(4,5)P_2_ or PS. Each dot indicates an anionic lipid. An anionic lipid is considered to be interacting with StarD4 when any of its atom is with 4Å of the protein. Lipids with dwell time below 200 ns are excluded from the analysis. **(A)** Data obtained Phase 2 simulations of 4μs per replica; **(B)** Data from Phase 3 simulation of 2μs per replica. Statistical analyses show that the dwell time of PS is significantly shorter than the dwell time of either PI(3,5)P_2_ or PI(4,5)P_2_ with p<0.0001 by T test.

**Supplementary Figure 12.**
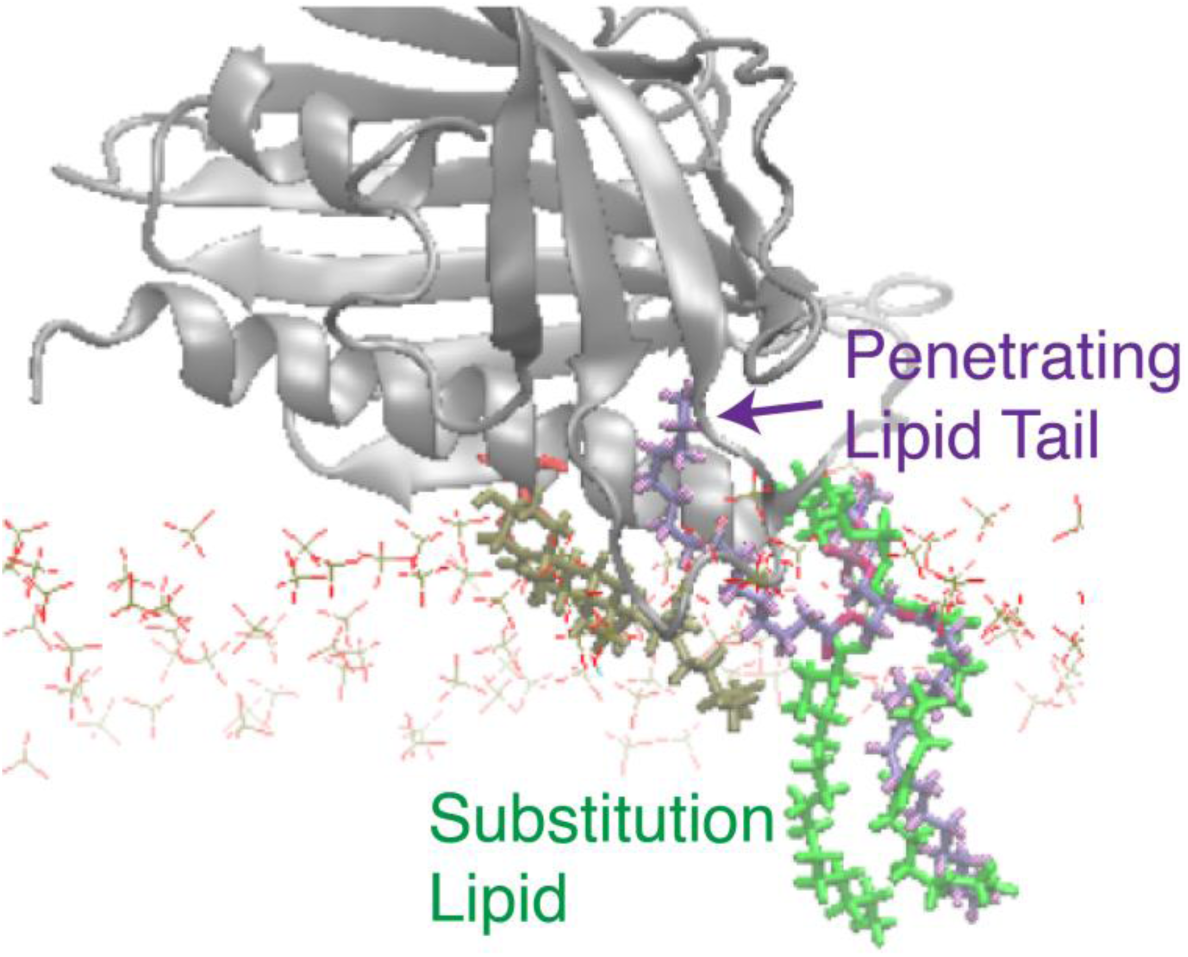
Seed construction by manual substitution of the gate-penetrating membrane lipid tail. To conduct the adaptive sampling of the extraction of membrane CHL by apo-StarD4, the initial seeds are chosen from the previous simulation with the membrane CHL positioned at the highest binding site, and an open H4-Ω1 gate. In the *apo*-StarD4, the opening of the H4-Ω1 gate is likely related to an inserted membrane lipid tail. In that case, we replaced the already inserted lipid with its own conformation it had before its insertion in the simulation in order to maintain the same interaction between the lipid and the protein in the expected insertion. The initial seeds bearing this manual modification are then equilibrated with the protein backbone and the target cholesterol constrained using the preparation protocol of StarD4-membrane complex as described.

**Supplementary Figure 13.**
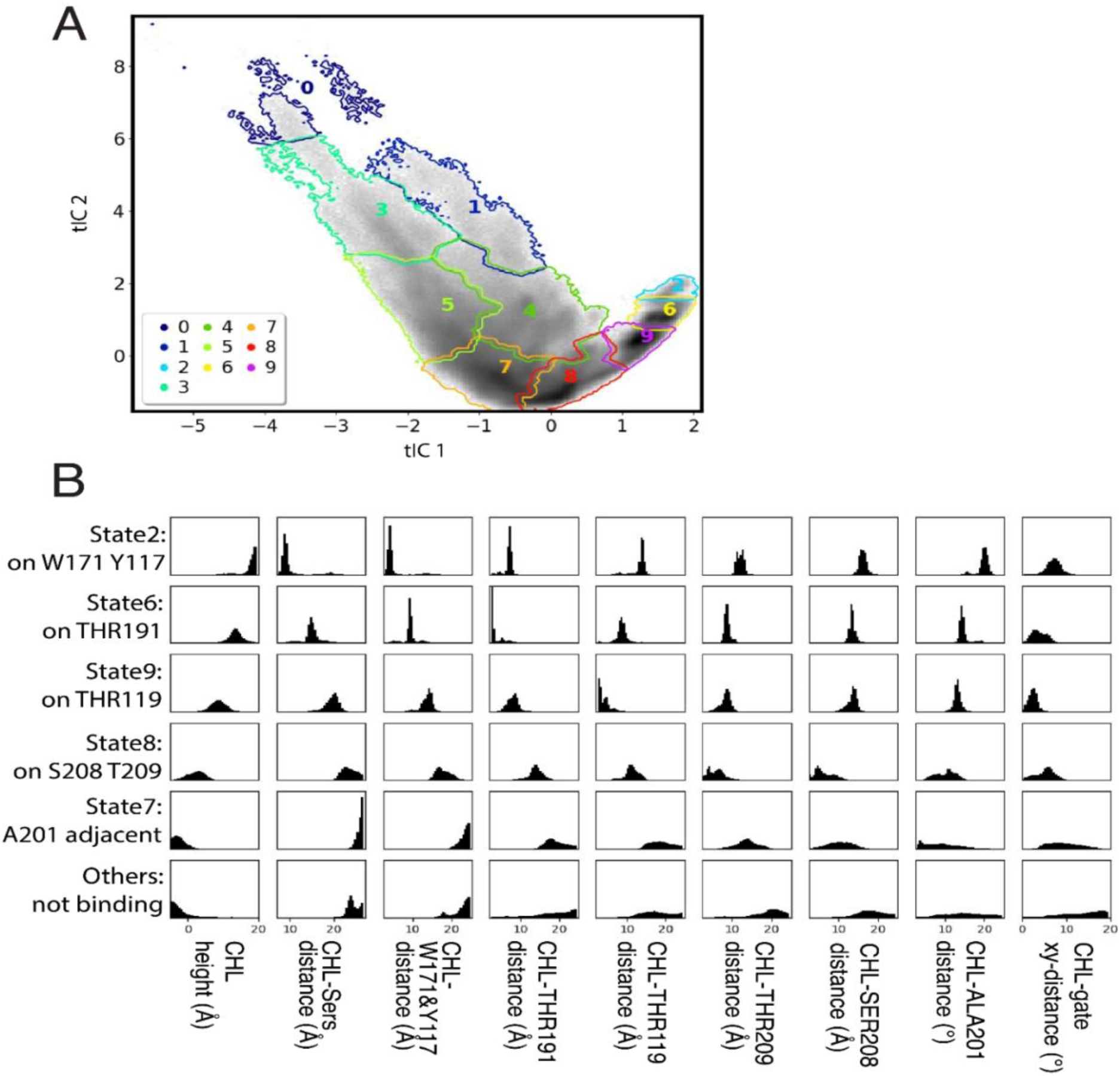
The characteristics of metastable states along the cholesterol uptake pathway. **(A)** 10 macrostates are outlined on the 2D tICA map defined in Fig. 18. **(B)** Structural characteristics of the macrostates along the cholesterol uptake pathway. The structural characteristics are represented by probability density histograms of the characteristic CV values that define the tICA space as in Fig. 18A. The states described (from top to bottom) are: macrostate 2 representing W171/Y117 binding, macrostate 6 representing T191 binding, macrostate 8 representing T119 binding, macrostate 7 representing S208/T209 binding. The other states are shown together.

**Supplementary Figure 14.**
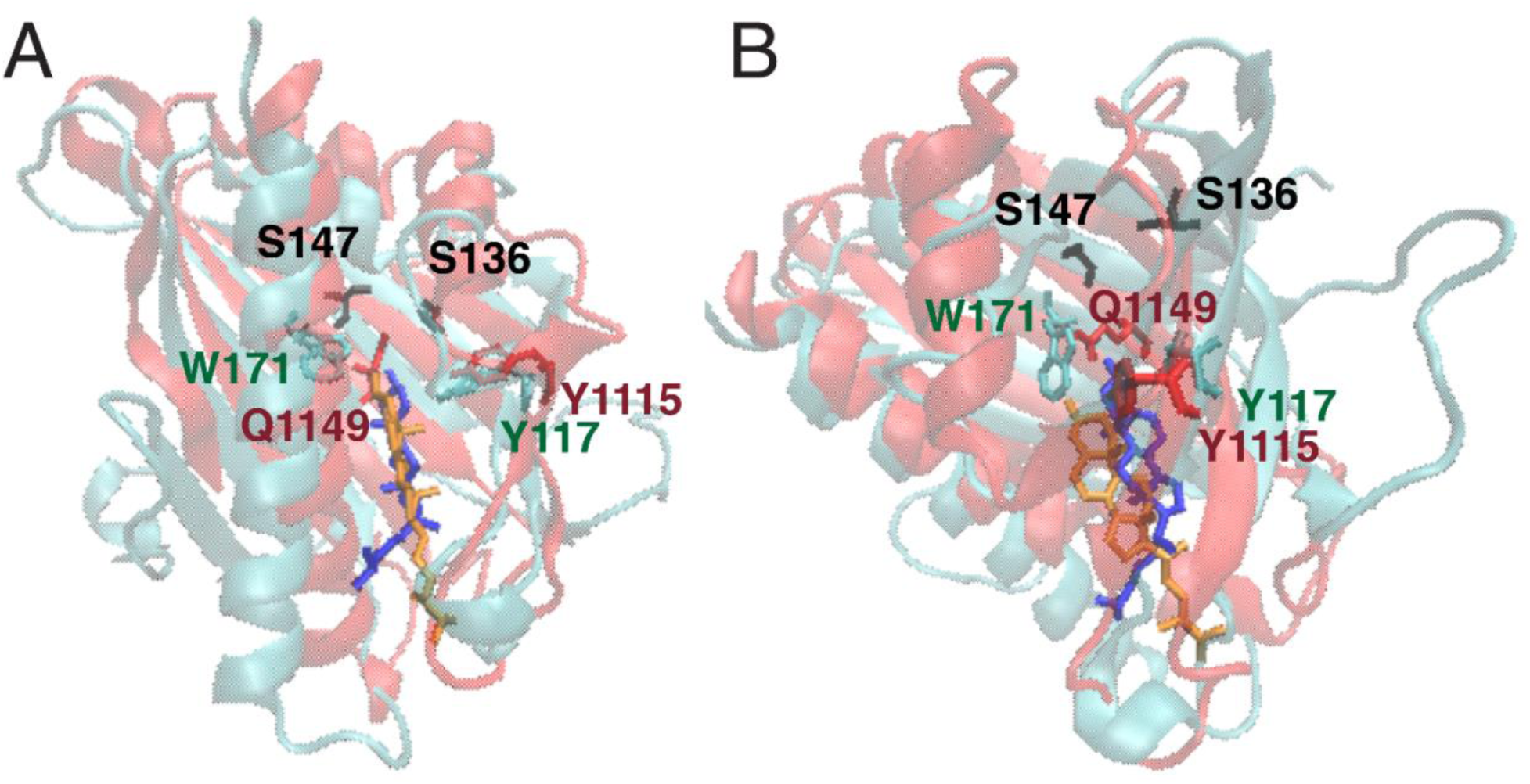
Comparison of sterol-binding conformations inthe crystal structure of LAM2 and the *holo*-StarD4 complex in simulation. **In both A and B** the representative conformation of the Trp-binding-CHL-StarD4 complex is rendered in cyan, with the W171 Y117 binding sites shown in cyan licorice, the S136 S147 in black licorice, and the cholesterol in blue licorice. The structure of LAM2 (PDB ID 5YS0) is rendered in red cartoon, with the sterol binding residues Q1149 Y1115 in red licorice, and its ligand sterol in orange licorice. Panel A: the front view; Panel B: the lateral view 90° from (A).

### SUPPLEMENTARY TABLES

**Supplementary Table 1:**
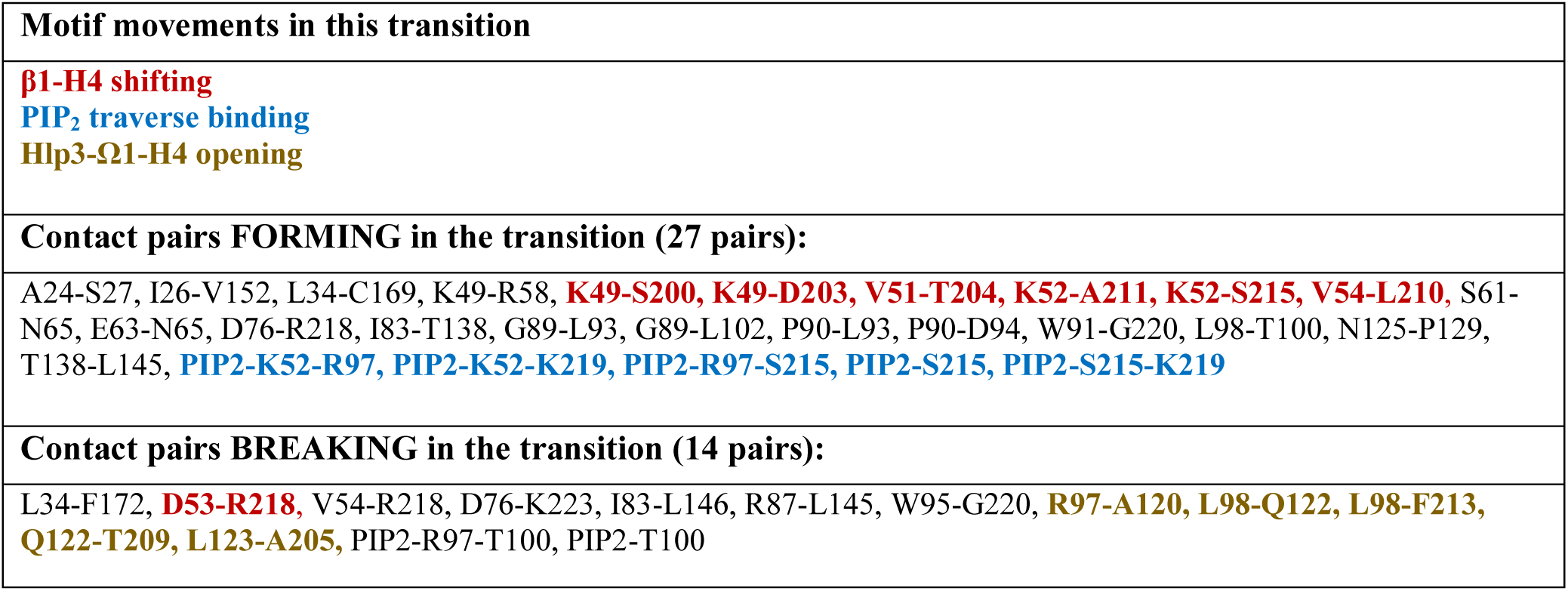
Structure-differentiating contact pairs in the transition event 2→3.

**Supplementary Table 2:**
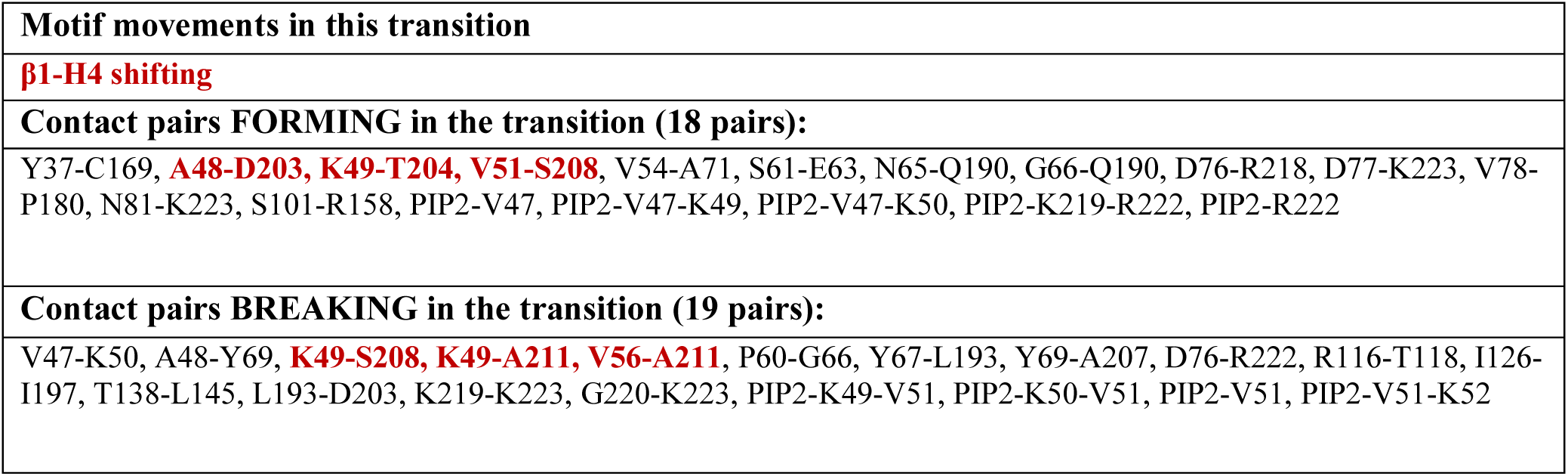
Structure-differentiating contact pairs in the transition event 0→1.

**Supplementary Table 3:**
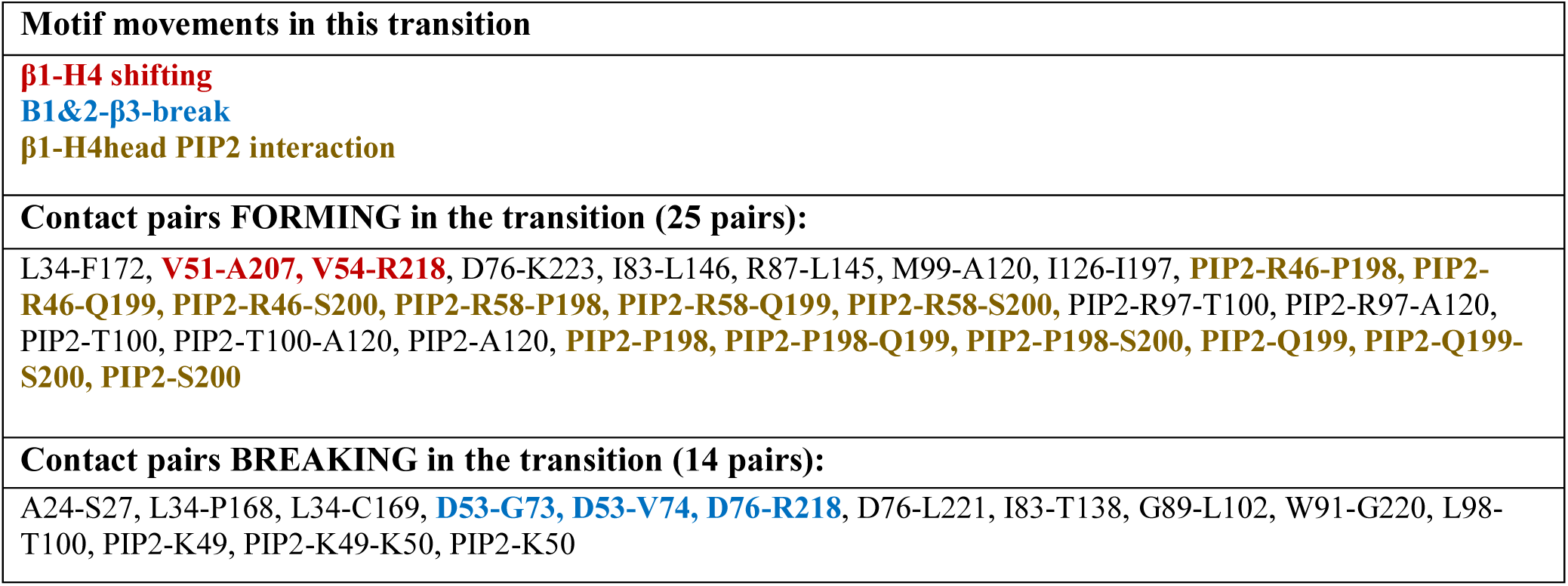
Structure-differentiating contact pairs in the transition event 1→2.

**Supplementary Table 4.**
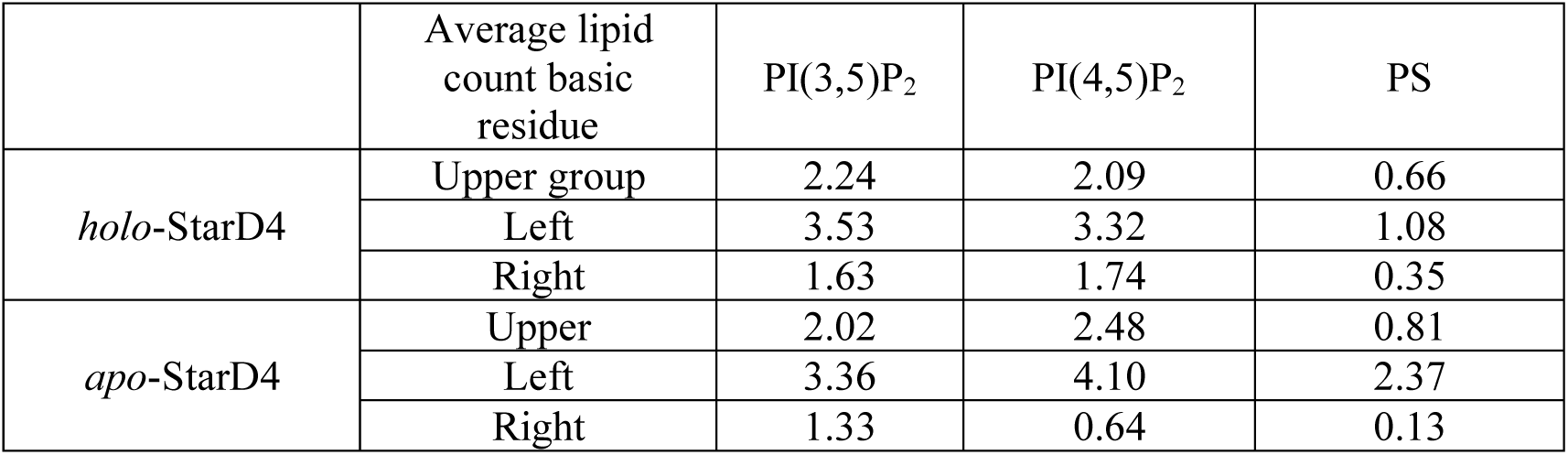
The average number of PIP2s binding to the basic residue groups. Upper group: K52, R97, S215, R218, K219, R222 Left group: R46, K49, K50, K52, R58 Right group: R116, R130, R158, R163, R194.

**Supplementary Table 5.**
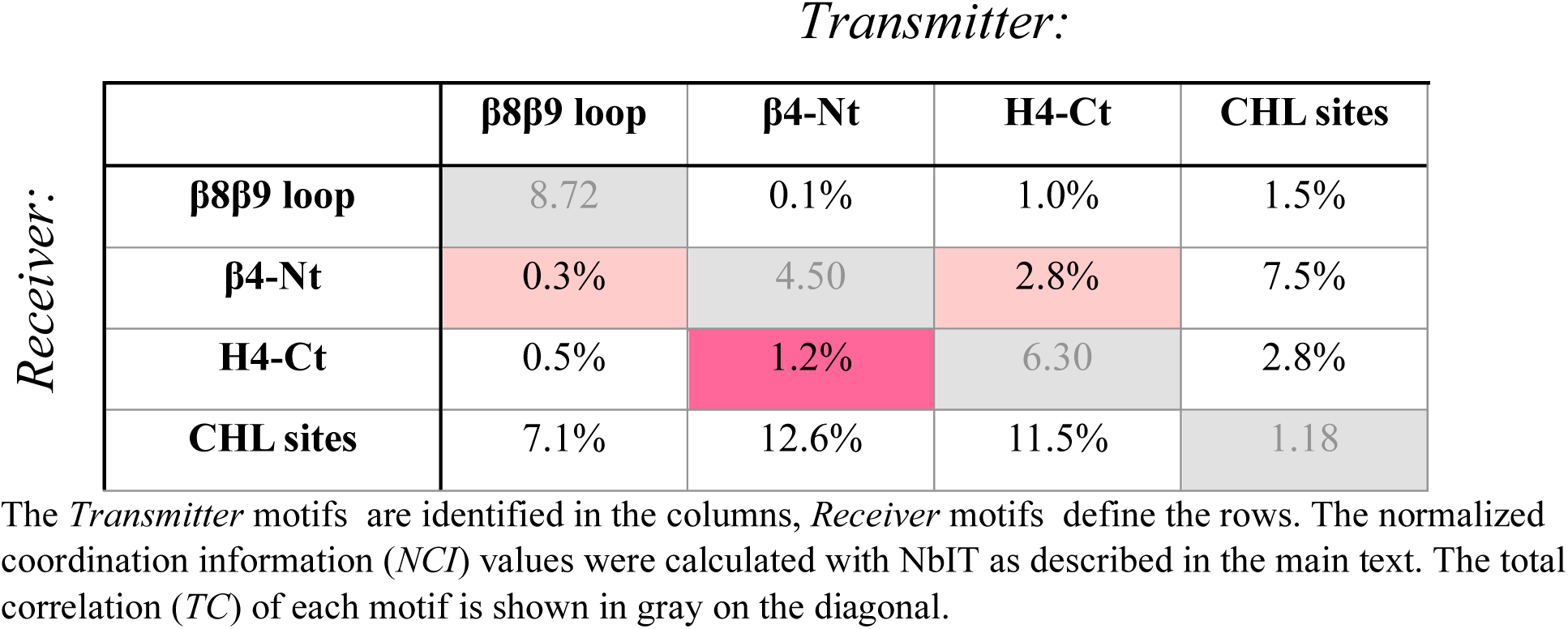
Normalized coordination information (*NCI*) between non-correlated sites that serve as negative controls of allostery, and indicate the background level of correlation noise resulting from the complex dynamics of the system.

| *Transmitter:* |  |  |  |  |  |
| --- | --- | --- | --- | --- | --- |
| *Receiver:* | | $\beta 8\beta 9$ loop | $\beta 4$-Nt | H4-Ct | CHL sites |
| | $\beta 8\beta 9$ loop | 8.72 | 0.1% | 1.0% | 1.5% |
| | $\beta 4$-Nt | 0.3% | 4.50 | 2.8% | 7.5% |
|  | H4-Ct | 0.5% | 1.2% | 6.30 | 2.8% |
|  | CHL sites | 7.1% | 12.6% | 11.5% | 1.18 |

The *Transmitter* motifs are identified in the columns, *Receiver* motifs define the rows. The normalized coordination information (*NCI*) values were calculated with NbIT as described in the main text. The total correlation (*TC*) of each motif is shown in gray on the diagonal.

**Supplementary Table 6.**
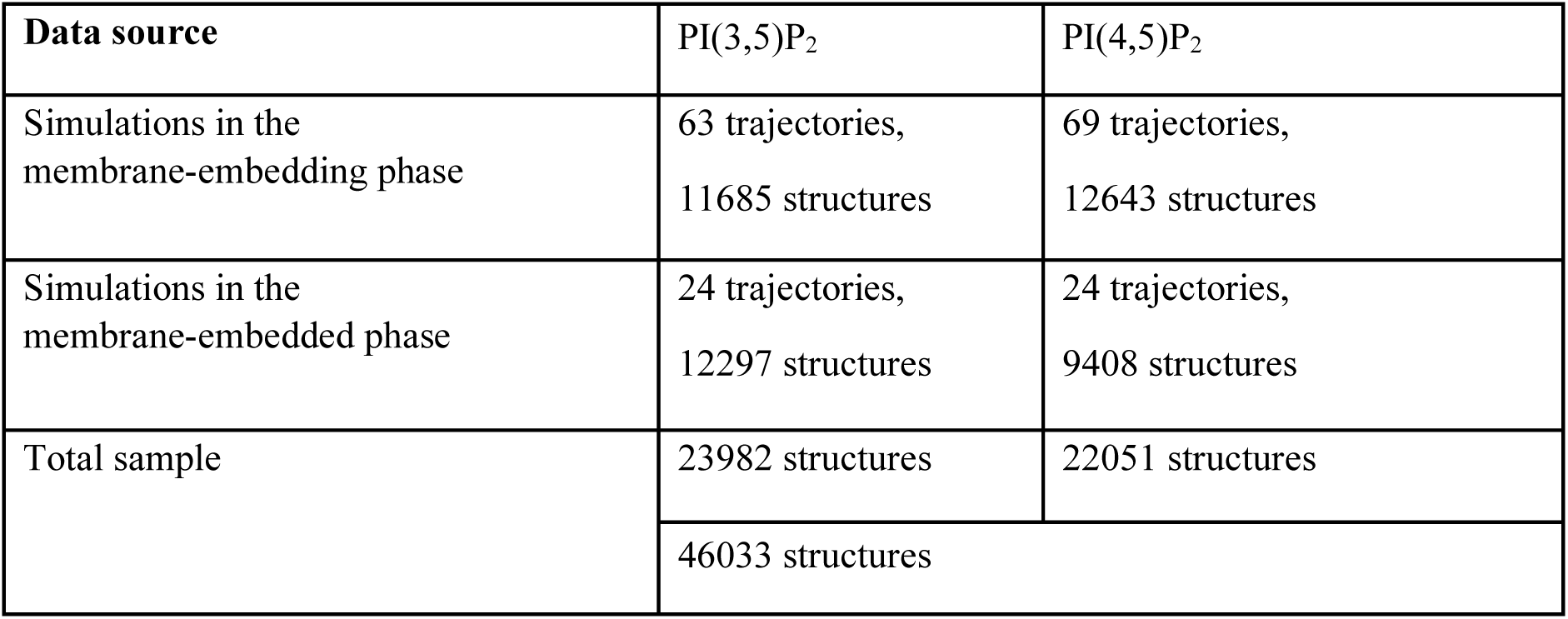
Origin of and sample numbers of simulation data used in the construction of DNN.

## Supplementary Methods

**Section 1.** Definitions of the collective variables capturing the dynamics of the cholesterol-StarD4-membrane complex and the events that lead to the pre-release state:

**H4 tilting angle (°):** The angle between the H4 norm vector and the membrane norm vector. See **Sup.** Fig. 1**. Protein orientation around H4 (°):** An angle defined in 3D space. See definition in **Sup.** Fig. 1.

**Water count within 4Å of CHL tail:** The number of waters within 4Å surrounding the eight-carbon branched aliphatic tail of the ligand cholesterol

**CHLtail membrane distance (Å):** The height of the center of mass (CoM) of atoms C22 C23 C24 C25 C26 C27 in CHL above the membrane surface defined by the average height of P atoms

**CHL-TYR117 distance (Å):** The distance from the CHL oxygen to the sidechain oxygen of Tyr117

**CHL-TRP171 distance (Å):** The distance from the CHL oxygen to the sidechain nitrogen of Trp171

**CHL-Sers distance (Å):** The distance from the CHL oxygen to the closest sidechain oxygen of Ser136 or Ser147

**CHL-W171&Y117 distance (Å):** The distance from the CHL oxygen to the CoM of Trp171 and Tyr117

**β1-H4 distance (Å):** The mean distance between the CoM of the sidechains of residue pairs: K49-S200, K49-D203, V51-T204, V51-A207, V51-S208, K52-A211

**PIP_2_#227-K52/R97/S215/K219 distance (Å):** Between a single PIP_2_ molecule (which is PIP_2_#227 in RT1) and a set of basic residues, measure the distances between the closest pairs from N or O atom in the basic residues to their closest O atoms in the PIP_2_

**Ω1-H4 distance (Å):** The mean distance between the CoM of the residue pairs: Q122-A205, Q122-T209, L123-I197, L123-A201, L123-A205, I127-I197, I127-A201, I127-A205

**Ω4-β6β8 distance (Å):** The mean distance between the backbone of the residue pairs D192-Y165, L193-G164, G195-V162, and between the backbone of R194 to the sidechain of R130

**β1-β2β3 distance (Å):** The mean distance between the backbone of the residue pairs K49-V56, K50-T55, V51-V54, K52-V74, D53-G73

**H4head-unfold (Å):** The mean distance between the CoM of the residue pairs Q199-D203, S200-T204, A201-A205

### Definition of interaction score between CHL body and Protein back

The interaction score between two atoms is defined by to their distance in the following way: The score is (6 minus the distance) when the distance is smaller than 6, and 0 when the distance is larger than 6. The interaction score between two motifs is the sum of interaction scores between all pairs of atoms of the two motifs.

The atoms considered are the carbons of the CHL (C1 to C19), and the heavy atoms the residues N166 C169 I189 Q190 T191 of the protein

**Section 2.** Definition of the collective variables capturing the dynamics of PIP2 binding on the upper basic residue patch:

**Distance to K52 (Å):** the distance between the N atom in LYS52 and the closest O atom in the lipid head of PIP2

**Distance to R97 (Å):** the distance between the closest pair of N atom in ARG97 and O atom in the lipid head of PIP2

**Distance to S215 (Å):** the distance between the O atom in S215 and the closest O atom in the lipid head of PIP2

**Distance to R218 (Å):** the distance between the closest pair of N atom in ARG218 and O atom in the lipid head of PIP2

**Distance to K219 (Å):** the distance between the N atom in LYS219 and the closest O atom in the lipid head of PIP2

**Section 3.** Definition of the collective variables capturing the dynamics of CHL along the CHL uptake pathway on apo-StarD4:

**Criteria of “CHL at the gate”:** The width of the Ω1-H4 gate is defined as the CoM distance between residues L123 I127 on Ω1 and residues A201 V202 A205 S208 on H4. An atom is considered located in between the Ω1-H4 gate, when both its distance to L123 I127 and its distance to A201 V202 A205 S208 are smaller than width of the Ω1-H4 gate divided by square root of 2. A membrane cholesterol is considered “at the gate” when any atom of the cholesterol is located in between the Ω1-H4 gate.

**Criteria of “CHL around the gate”:** A membrane cholesterol is considered “around the gate” when any atom of the cholesterol is located within 5Å of any atom in residues L123 I127 A201 V202 A205 S208.

**Random in 15∼25Å:** A membrane cholesterol is considered “Random in 15∼25Å” when any atom of the cholesterol is located within 25Å but not within 15Å of any atom in residues L123 I127 A201 V202 A205 S208.

**Membrane CHL tilting angle (°):** The angle between the membrane norm vector and the cholesterol axial vector (from the CoM of atoms C13 C14 C15 C16 C17 to the atom O3)

**CHL-height (Å):** The height of atom O3 of the cholesterol above the membrane surface defined by the average height of P atoms

**CHL-THR209 distance (Å):** The distance between the atom O3 of cholesterol to the sidechain oxygen in THR209

**CHL-SER208 distance (Å):** The distance between the atom O3 of cholesterol to the sidechain oxygen in SER208

**CHL-ALA201 distance (Å):** The distance between the atom O3 of cholesterol to the CoM of sidechain atoms in ALA201

**CHL-THR119 distance (Å):** The distance between the atom O3 of cholesterol to the sidechain oxygen in THR119

**CHL-THR191 distance (Å):** The distance between the atom O3 of cholesterol to the sidechain oxygen in THR191

**CHL-gate xy-distance (Å):** The distance between the atom O3 of cholesterol to the CoM of residues Q122 L123 I127 V202 A205 T209

**Ω1-Ω4 distance (Å):** The mean distance between the CoM of the residues R194 G195 M196 I197 on the Ω4-loop and the residues I126 I127 S128 on the Ω1-loop

**Section 4.** equilibration of the membrane-StarD4 complex at a distance using Mean-field model

In the MFM approach, StarD4 is represented at the detailed 3D atomistic level, while the membrane is simplified as a two-dimensional smooth charged surface that represents the behavior of lipid polar headgroups. In this hybrid model, StarD4 is placed above the membrane surface with the closest atom located 2Å away from the membrane surface. As described in ref (54,56), self-consistent minimization is employed to sample the rearrangement of membrane lipid position corresponding to the StarD4s by optimizing the governing mean-field-based free energy function (*F*), which comprises of the *electrostatic component* (*F*_*el*_), the *translational entropy* of mobile ions (*F*_*IM*_) that depends on the local lipid component densities φ_(*x*,*y*)_, and the *lipid mixing entropy* (*F*_*lip*_) that depends on the mobile ion concentrations *c*^−^ and *c*^+^ according to:

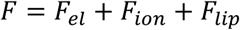

The *electrostatic component* is determined by:

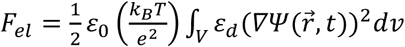

where 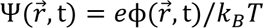 is the dimensionless electrostatic potential, with ϕ representing the electrostatic potential, and *e* being the elementary charge, ε_0_ being the vacuum permittivity, ε_*d*_ being the dielectric constant (ε_*d*_ = 80 in aqueous solution and 2 within protein and membrane).

The *translational entropy* of mobile ions is determined by:

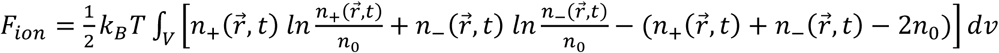

where 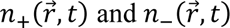 denotes the local concentrations of the positive and negative ions in the solution, and *n*_0_ is the bulk concentration of the ions.

The *lipid mixing entropy* is determined by:

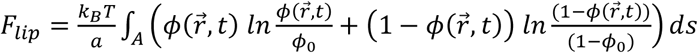

where 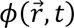 is the local mole fractions of the charged lipids in the membrane, and ϕ_0_ is the average composition of the charged lipid.

The minimization of this free energy function is carried out under the influence of the electrostatic field generated by the StarD4 protein, which yields a correspondingly optimized lipid rearrangement that is quantified from the local lipid density. The absorption energy of StarD4 into the membrane is determined by comparing the free energy of the StarD4-complex to the sum of the free energies of StarD4 and of the membrane considered separately. Here, the calculations were carried out in a (256 Å)^3^ cubic space containing a 0.15M ionic solution of monovalent counterions, corresponding to λ=8.09Å Debye length, under the temperature of 310K.

The positioning of StarD4 is systematically explored in various orientations while ensuring that all atoms are at least 2Å away from the membrane surface. The orientation of StarD4A is defined with a spherical coordinate, with its origin located at the center of mass of StarD4, its z-axis parallel to the norm vector of H4, and its x-axis perpendicular and intersect with the norm vector of H4. The absorption energy of StarD4 is first examined with StarD4 facing the membrane from six orientations:

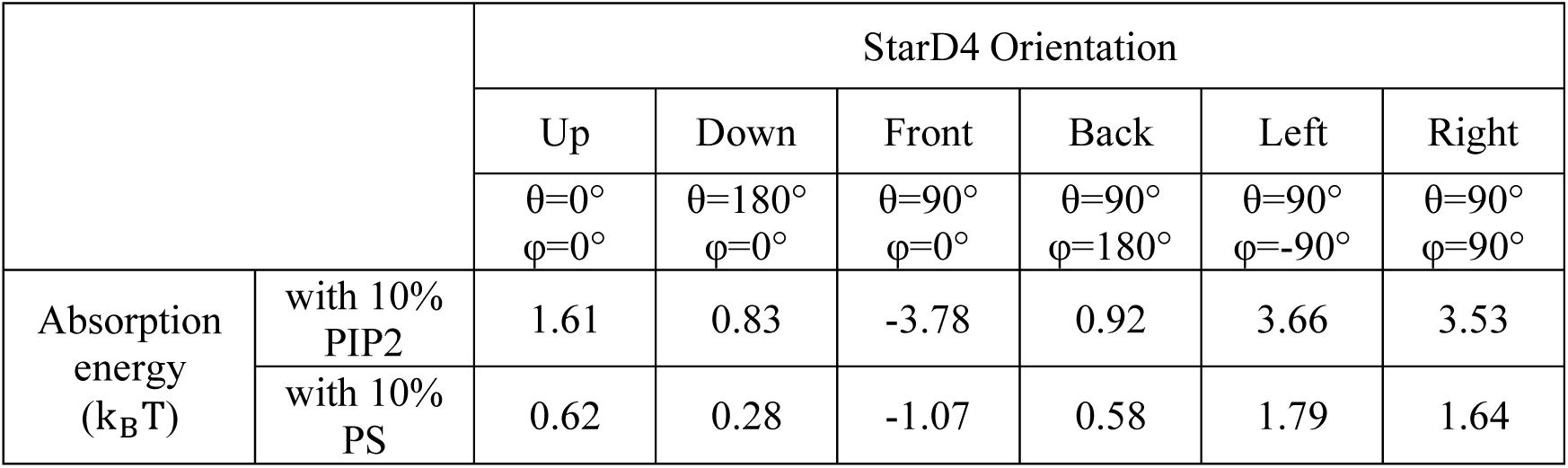

The best orientation of the membrane approaching StarD4 is determined as the position that yields the strongest absorption energy among the geometries sampled. Among the six orientations, the “Front” appeared as the most preferred orientation of StarD4, where its H4 is parallel to the membrane surface. To verify this result, eight more orientations are sampled with StarD4 rotated 45° from the “Front” direction, and the result confirmed that the “Front” direction yields the strongest absorption energy at this stage.

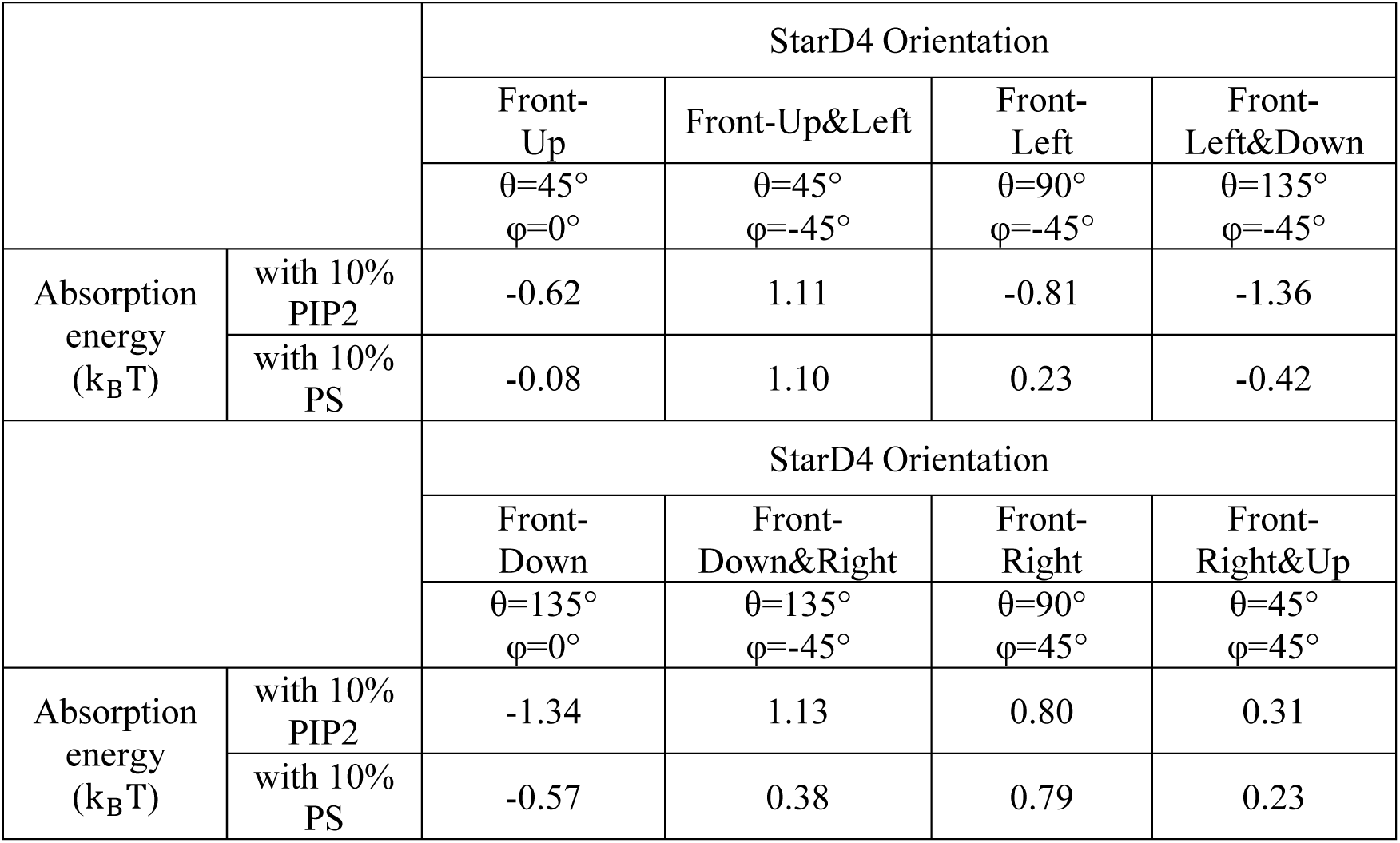

## Notes

### Competing Interest Statement

The authors have declared no competing interest.

### Summary of Updates

Streamline the presentation, add details for reproducibility, Figure 7 revised, Title changed, Supplementary files updated.

