## Supplementary Information- Complete for "Recognition of specific PIP2-subtype composition triggers the allosteric control mechanism for selective membrane targeting of cargo loading and release functions of the intracellular sterol transporter StarD4"

**Contains:**

A. **14 Supplementary Figures**
B. **5 Supplementary Tables**
C. **3 Sections of Supplementary Methods (definitions)**

#### A. SUPPLEMENTARY FIGURES

##### Supplementary Figure 1.

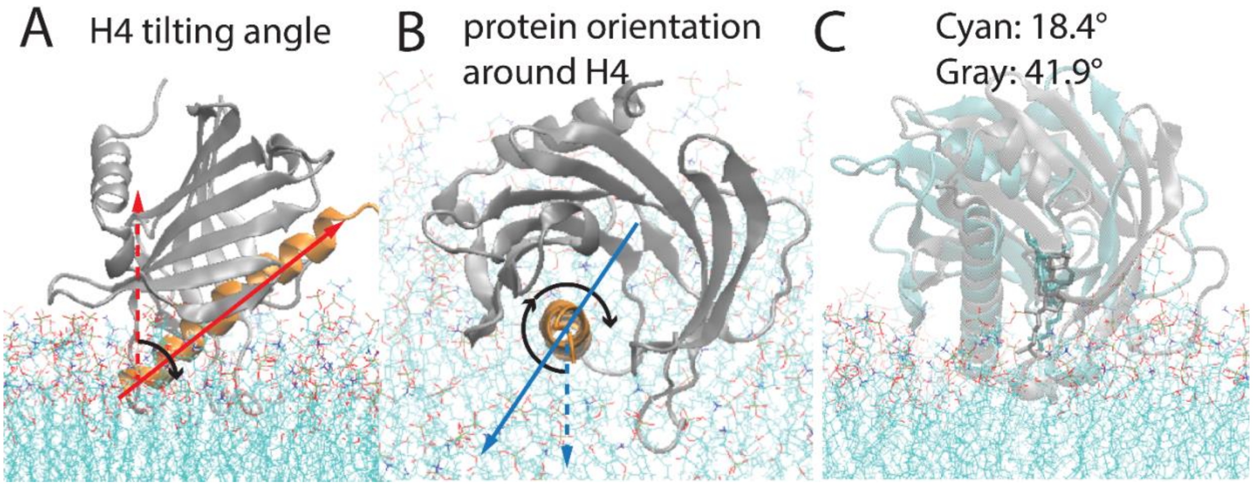


**The orientation of StarD4 in relation to the membrane bilayers.**

**(A)** The definition of the H4 tilting angle: the angle between the "membrane norm" (red dashed line) and the "Helix4 axis" (red solid line, the first principal component of the Cα atoms of H4).

**(B)** The definition of the protein orientation around H4: The angle is measured on the plane perpendicular to the Helix4 axis. The "membrane norm" (blue dashed line) and the "vector from the center of mass of StarD4 to the center of mass of H4" are the projected onto the plane perpendicular to the Helix4 axis, and the CV is defined by the angle between the two projections.

**(C)** The representative conformations of the cholesterol-StarD4 complex at the protein orientation around H4 equals 18.4° (cyan) and equals 41.9° (gray).

#### Supplementary Figure 2.

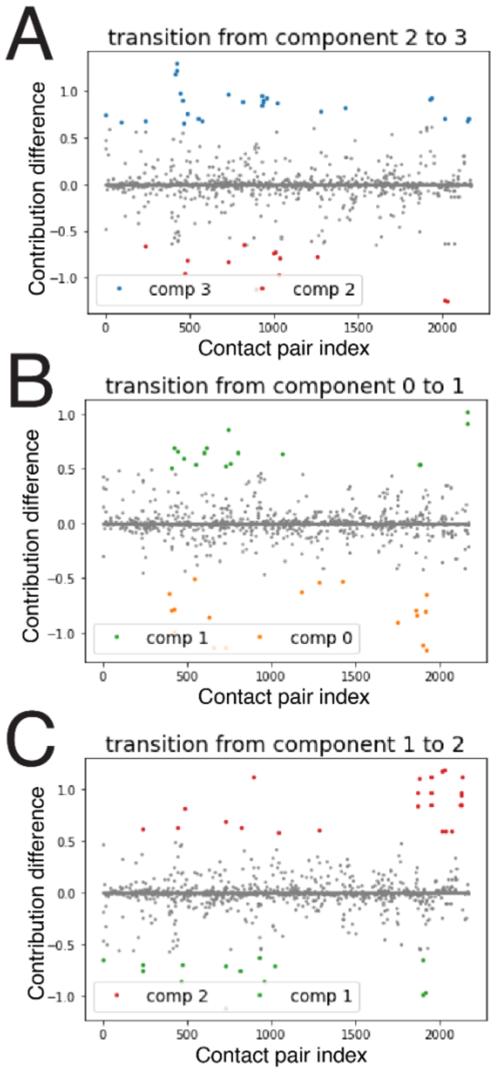


**The structure-differentiating contact pairs (SDCPs) are revealed through the comparison of the normalized spatial components. (A) SDCPs** for the transitioning of component 2→3; **(B)** for the transitioning of component 0→1; **(C)** for the transitioning of component 1→2. On the X axis are the 2171 data points representing the residue pairs. The Y axis shows the difference in the contribution of a residue pair to the conformational state in the transition between the two components. A positive contribution difference reflects the creation of a contact in this pair during the transition. A negative contribution difference reflects the breaking of the pair contact during the transition.

#### Supplementary Figure 3.

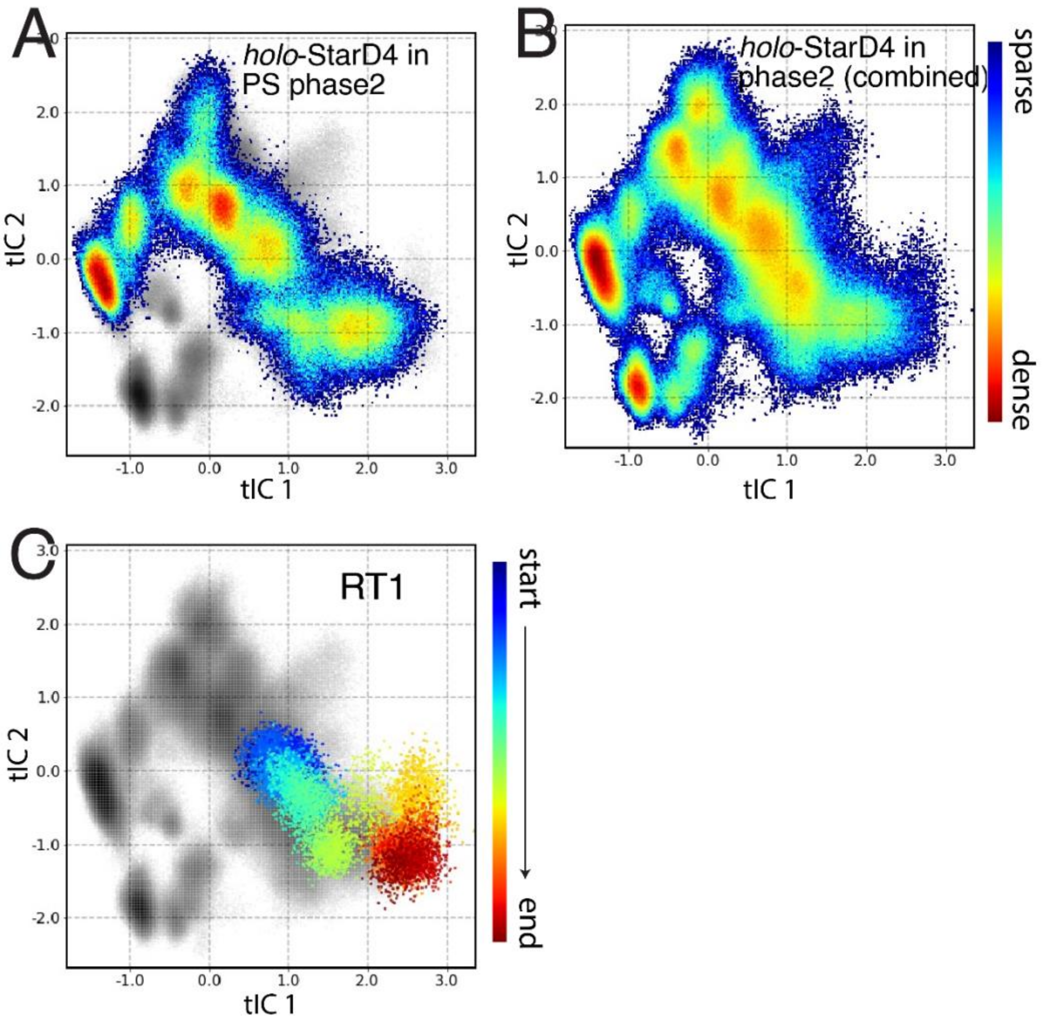


**Population density maps of the conformational space of *holo*-StarD4. (A) *holo*-StarD4** embedded on PS-containing membranes; **(B)** The combined population density maps for ***holo*-StarD4** on all three types of membranes. The color scale for the representation of population densities is shown on the right; **(C)** The time-evolution of the cholesterol releasing trajectory RT1 projected onto the combined conformational space. The trajectory evolution is color-coded from the starting point in blue, to the end point in red. The gray scale in the background represents the combined population density as in **(B)**.

**Supplementary Figure 4.**

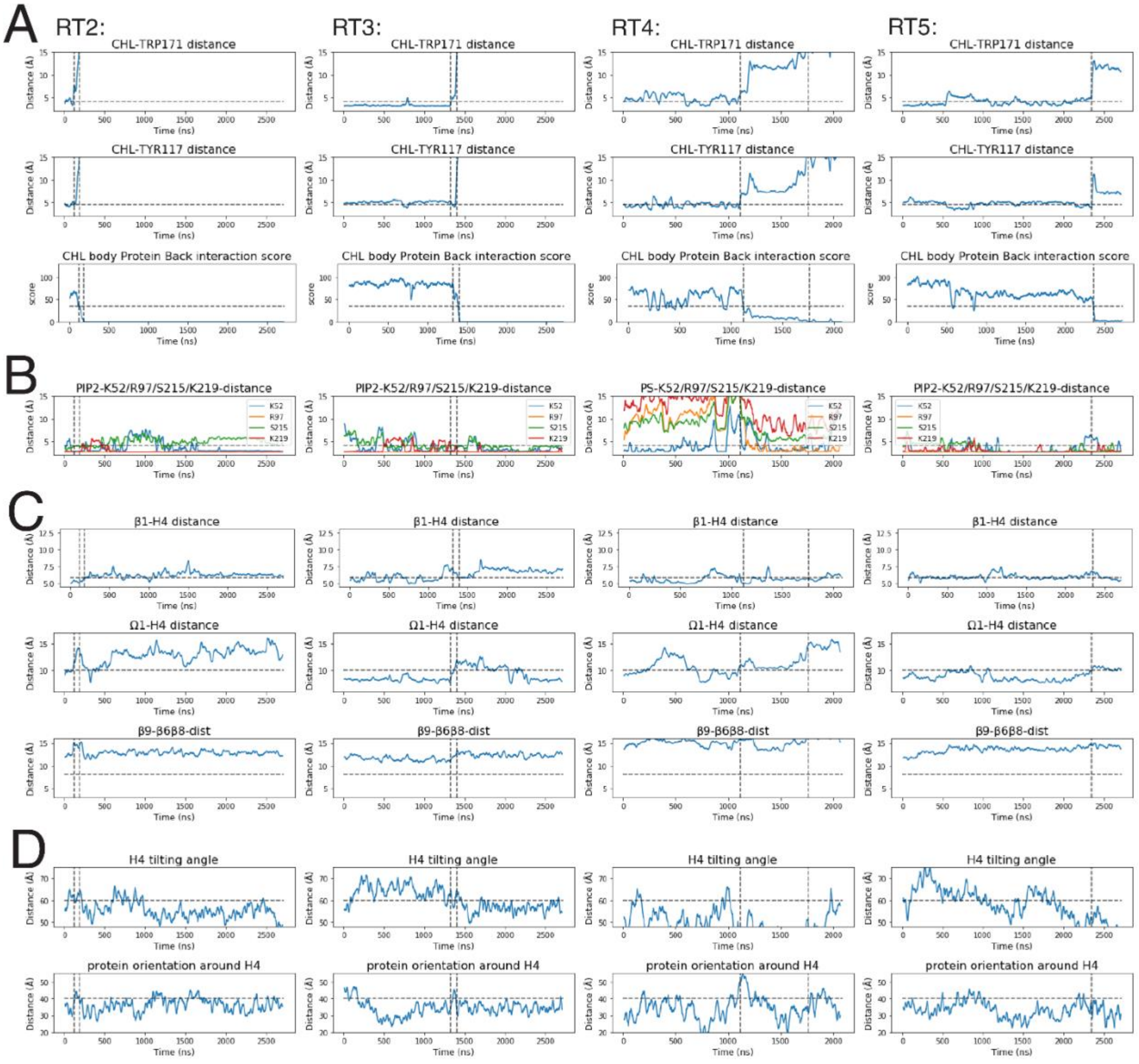


**The conformational changes related to the pre-release state in the Release Trajectories 2-5.** Data for RT2 to RT5 are shown in the columns from left to right. The rows depict: **(A)** The time-evolution of the CHL position relative to the binding sites in the pocket, and of the interaction between the CHL hydrocarbon ring and residues N166, C169, I189, T191 on β8 and β9 at the back of StarD4. **(B)** The time-evolution of the binding of anionic lipid in the cross-H4-binding mode. **(C)** The time-evolution of the conformational changes of StarD4, including the repositioning of H4 away from Ω1 towards β1, and the opening of Ω4 from β8&6. **(D)** The time-evolution of the orientation of StarD4 in relation to the membrane. The vertical dashed lines indicate timing of the CHL release process as shown in **(A)**. The horizontal dashed lines are auxiliary lines that indicate the features of the pre-release state.

#### Supplementary Figure 5.

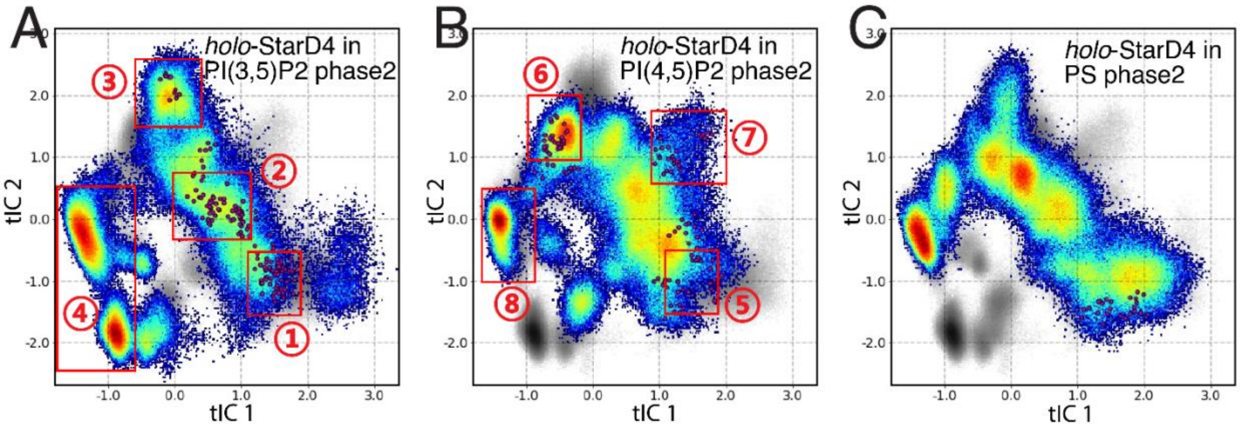


**The initial seeds of the adaptive sampling in Phase3**. The initial seeds of Phase3 simulations are indicated by red spots located at different conformational states on the tICA space of *holo*-StarD4 on the PI(3,5)$P_2$-containing membrane **(in A)**, on the PI(4,5)$P_2$-containing membrane **(in B)**, or on the PS-containing membrane **(in C)**.

### <u>Supplementary Figure 6.</u>

The initial structures of the (**RT6**, **RT7**) set feature an Ω1-open configuration on PI(4,5)$P_2$-containing membranes (**Fig. 7B** state 7), characterized by the wide opening of the Ω1-H4 gate (**Fig. 7D**, 4$^{th}$ column), and they shared with the pre-release state a similar pattern of a widely opened β8-Ω4 corridor (**Fig. 7D**, 3$^{rd}$ column), but without the characteristic β1-H4 interaction (**Fig. 7D**, 6$^{th}$ column). These features are likely related to the membrane lipid tail that had inserted between the Ω1 loop and the CHL in the initial frames (**Suppl. Fig. 6A**), which resulted in a persistent and wide opening of the Ω1-H4 gate (centered at CV "Ω1-H4-dist"=14Å). Notably, this feature is not observed in the pre-release state shared among RT1-RT5 (**Fig. 7D**, 4$^{th}$ column, **Suppl. Fig. 6B,C**)

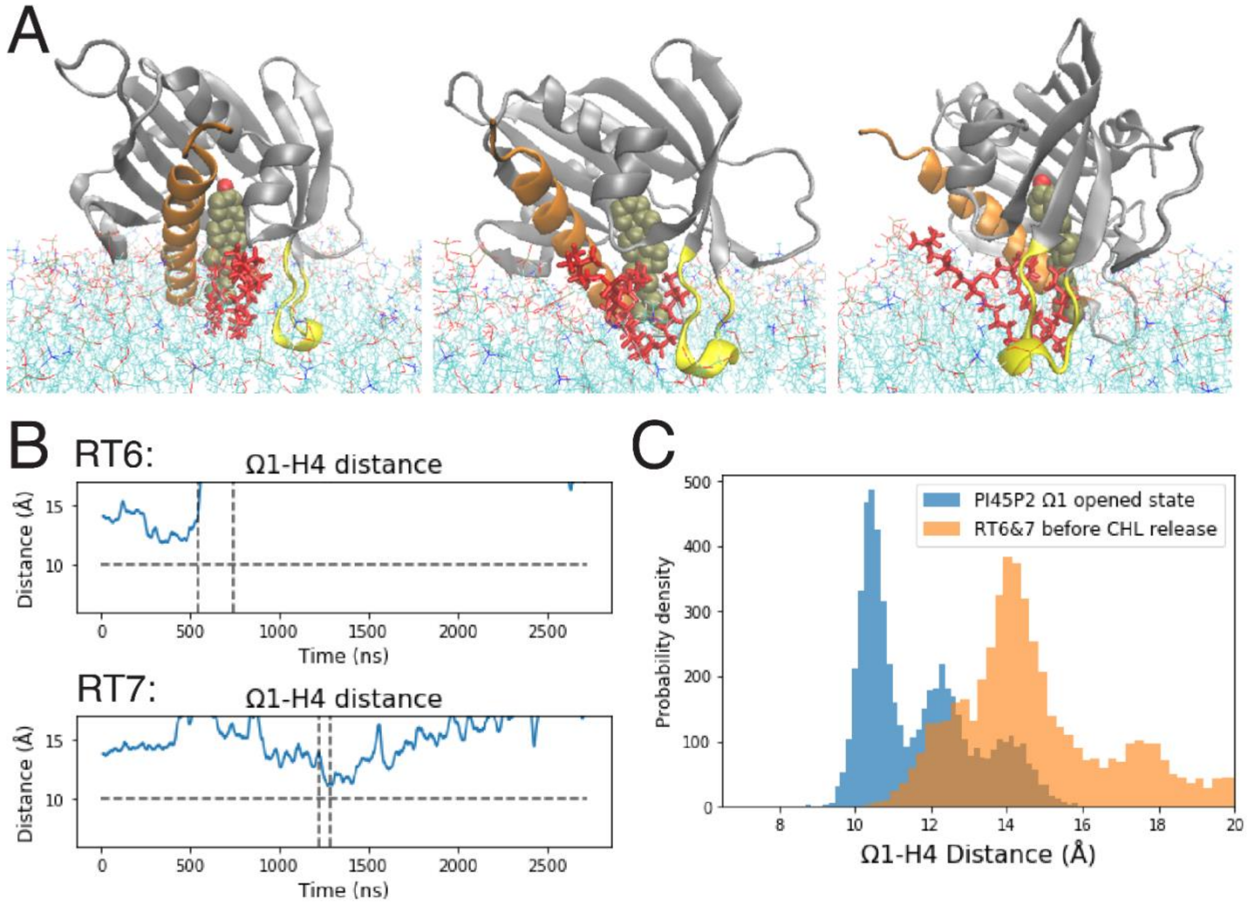


**The CHL release path initiated from the high energy state with a widely opened Ω1-H4 gate due to an inserted lipid tail. (A)** The membrane lipid inside the Ω1-H4 gate of *holo*-StarD4 depicted from 3 different viewing angles 45° apart from the front to the right. The StarD4 is rendered in gray cartoon with H4 in orange and Ω1 in yellow. The cholesterol in shown in VDW, and the gate-penetrating lipid is shown in red licorice. **(B)** The time-evolution of the widely-opened the Ω1-H4 gate. The horizontal dashed line is an auxiliary line that indicates the threshold of the opening of Ω1-H4 gate. The vertical dashed lines indicate the CHL release process. **(C)** Comparison of the probability density histograms of the Ω1-H4 distance in the Ω1-H4 opened state (blue) and the release trajectories 6 and 7 (orange).

The best orientation of the membrane approaching StarD4 is determined as the position that yields the strongest absorption energy among the geometries sampled. Among the six orientations, the "Front" appeared as the most preferred orientation of StarD4, where its H4 is parallel to the membrane surface. To verify this result, eight more orientations are sampled with StarD4 rotated 45° from the "Front" direction, and the result confirmed that the "Front" direction yields the strongest absorption energy at this stage.

| | | StarD4 Orientation | | | |
|---|---|---|---|---|---|
| | | Front-Up | Front-Up&Left | Front-Left | Front-Left&Down |
| | | θ=45° φ=0° | θ=45° φ=-45° | θ=90° φ=-45° | θ=135° φ=-45° |
| Absorption energy ($k_BT$) | with 10% PIP2 | -0.62 | 1.11 | -0.81 | -1.36 |
| | with 10% PS | -0.08 | 1.10 | 0.23 | -0.42 |
| | | StarD4 Orientation | | | |
| | | Front-Down | Front-Down&Right | Front-Right | Front-Right&Up |
| | | θ=135° φ=0° | θ=135° φ=-45° | θ=90° φ=45° | θ=45° φ=45° |
| Absorption energy ($k_BT$) | with 10% PIP2 | -1.34 | 1.13 | 0.80 | 0.31 |
| | with 10% PS | -0.57 | 0.38 | 0.79 | 0.23 |

**<u>Supplementary Figure 7</u>.**

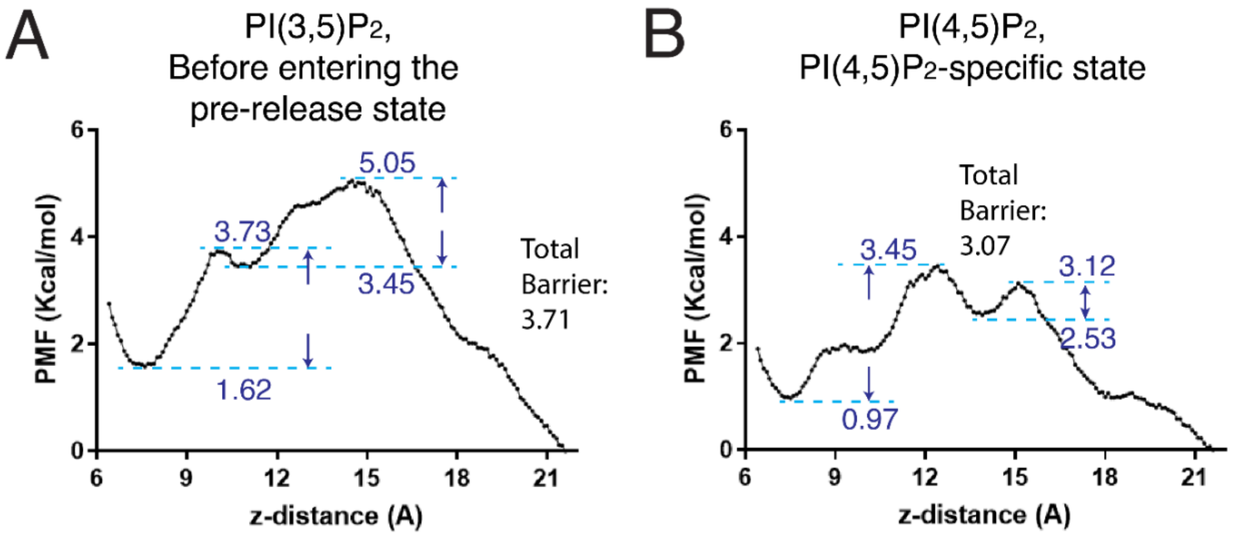


**The energy barrier along the reaction coordinate of CHL release from StarD4**, sampled with steered MD simulation and evaluated with umbrella sampling. The barrier is represented in terms of the potential of mean force. The reaction coordinate is the z-distance between the CHL and StarD4. The initial point is the common state on the PI(3,5)$P_2$-containing membrane **(A**, state 2 in Fig. 6**),** and the PI(4,5)$P_2$-specific state **(B**, state 6 in Fig. 6**)**. Local maxima and minima are labeled along the reaction path, and the total barrier is shown in the sidenote.

#### Supplementary Figure 8.

**Characteristics of the $PIP_2$ binding modes (modes 2 to 8 from top to bottom).** The probability density histograms of the characteristic CV values show (from left to right) the interactions of $PIP_2$ with the key residues K49, K50, K52, R97, S215, R218, K219, R222. **(A-C)** Show the characteristics of $PIP_2$ binding modes in 4 scenarios: **(A)** PI(3,5)$P_2$ binding on *holo*-StarD4, **(B)** PI(3,5)$P_2$ binding on *apo*-StarD4, **(C)** PI(4,5)$P_2$ binding on *holo*-StarD4, and **(D)** PI(4,5)$P_2$ binding on *apo*-StarD4.

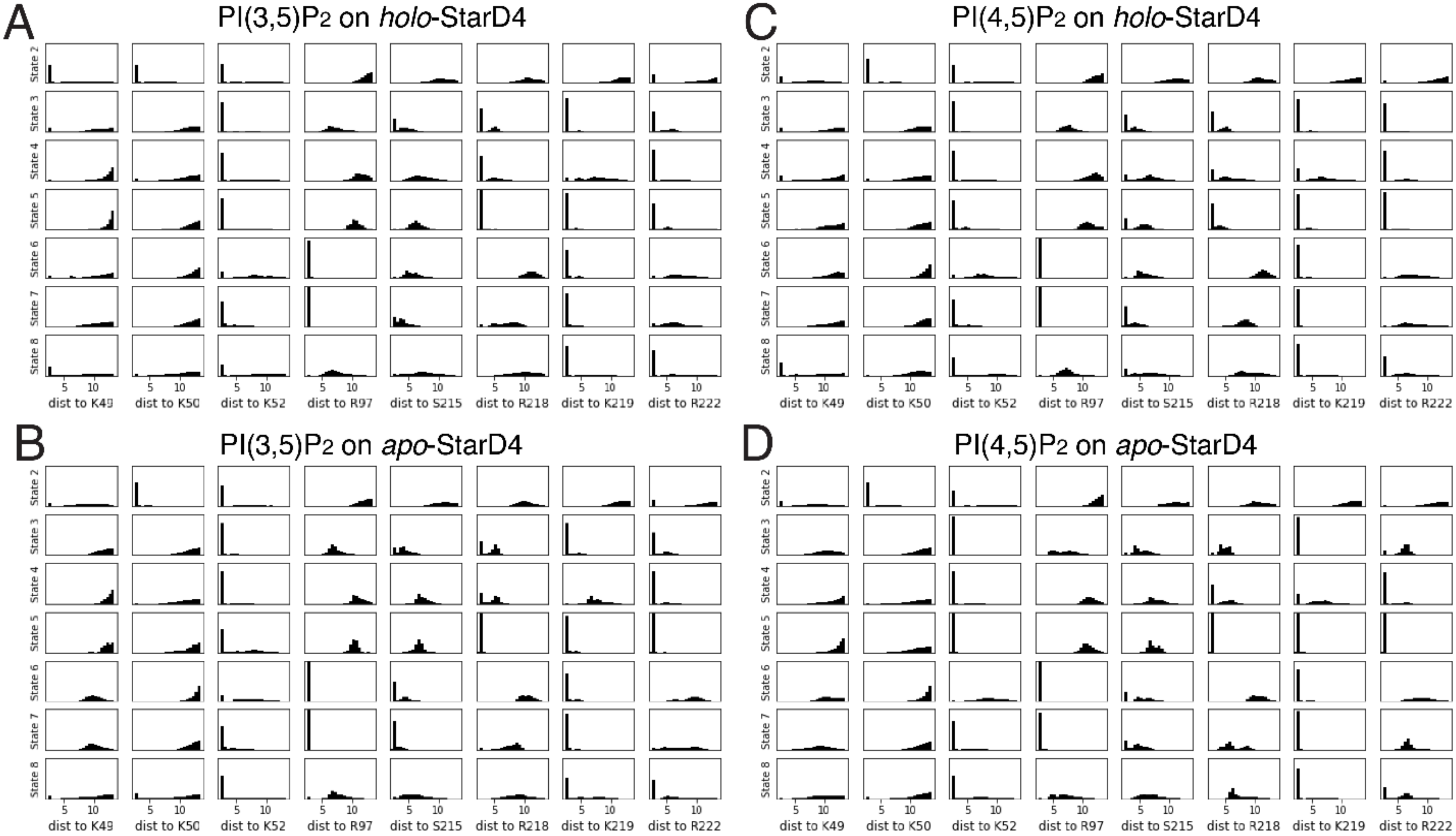

#### Supplementary Figure 9.

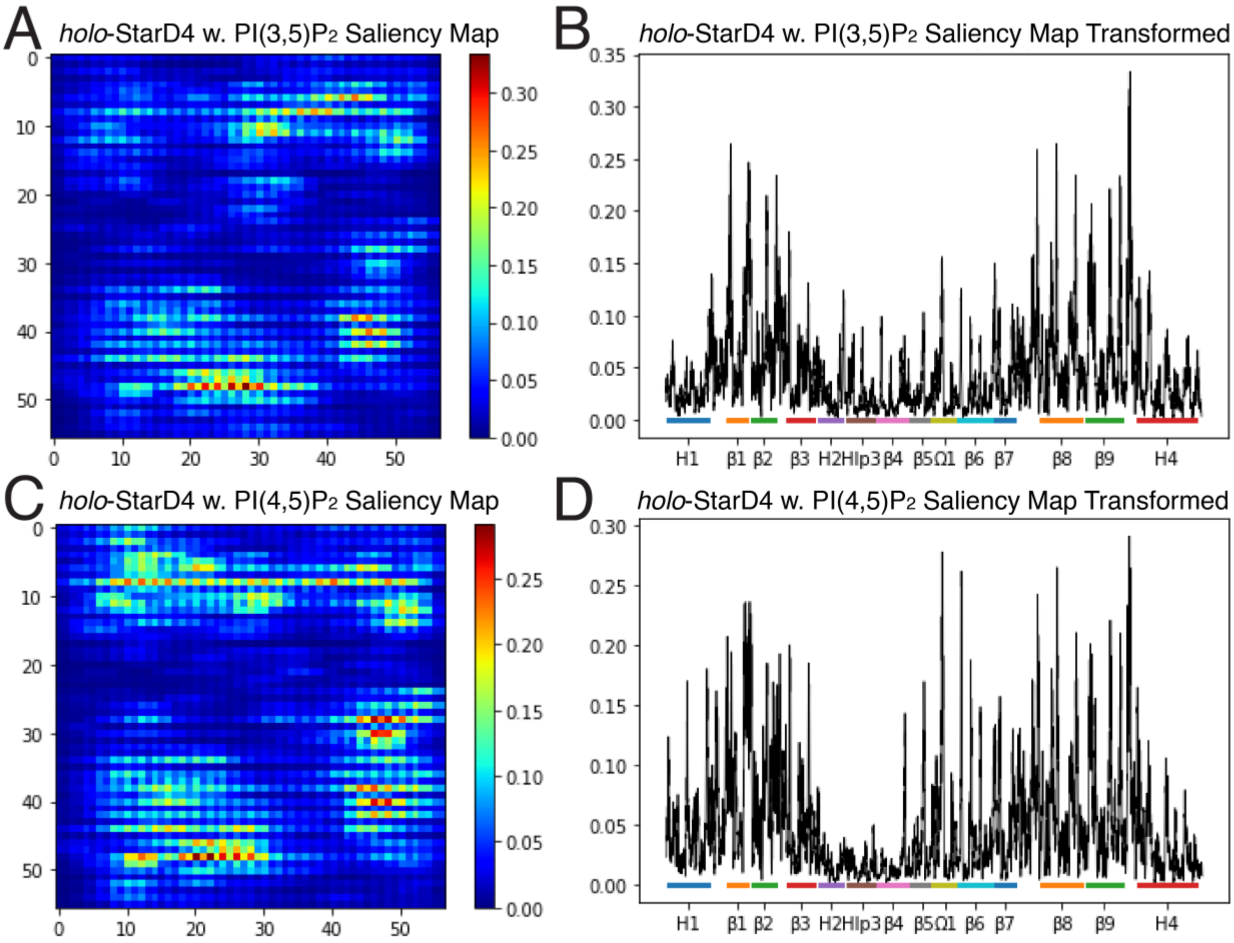


**Identification of the molecular determinants of the decision of DNN.** The attention maps of the atoms contributing to the decision of DNN in **(A)** the identification of StarD4 with PI(3,5)$P_2$-binding, and **(C)** the identification of StarD4 with PI(4,5)$P_2$-binding. Each pixel in the picture represents an atom, and the color code indicates its contribution to the decision, according to the color scale at the right. In **(B,D)** the attention maps shown in (A,C) are reshaped to list all the atoms on the x-axis, while the y-axis indicates their respective contribution to the decision. The color bar and tick marks on the x-axis identify the motifs to which the atom belongs.

#### Supplementary Figure 10.

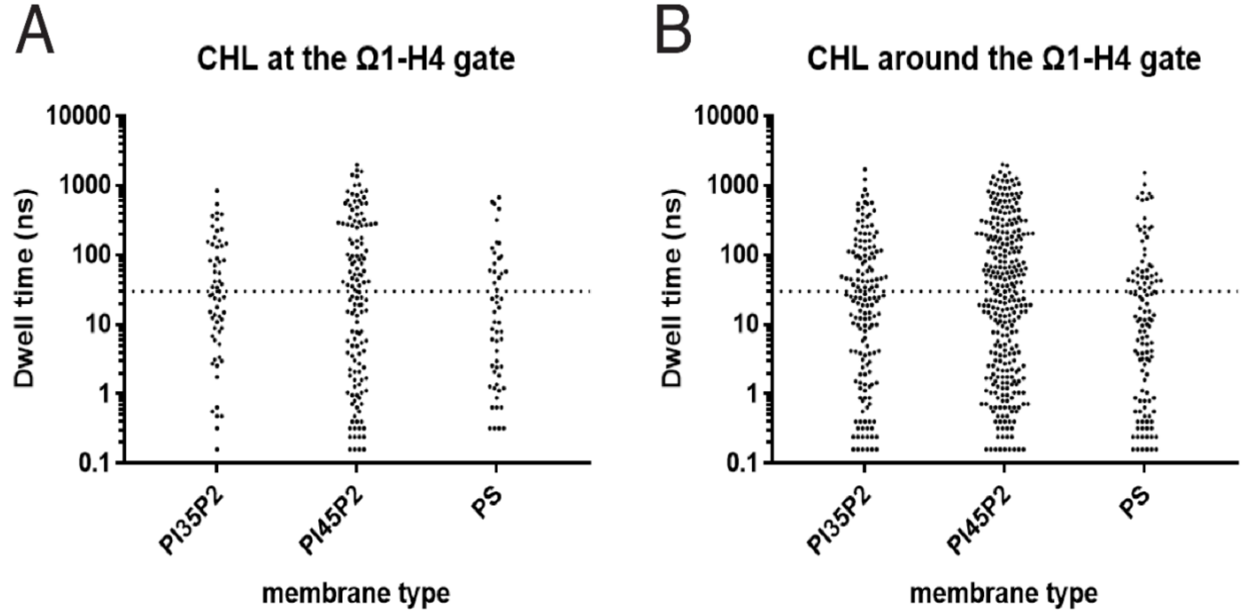


Lipid with dwell time > 30 ns:

| | Number | Average dwell time |
|---|---|---|
| PI35P2 | 28 | 190 ns |
| PI45P2 | 69 | 387 ns |
| PS | 19 | 196 ns |

Lipid with dwell time > 30 ns:

| | Number | Average dwell time |
|---|---|---|
| PI35P2 | 46 | 231 ns |
| PI45P2 | 147 | 362 ns |
| PS | 36 | 208 ns |

**Scatter plots of membrane CHL dwell time adjacent to StarD4, compared between the StarD4 embedded on PI(3,5)$P_2$-, PI(4,5)$P_2$- and PS-containing membrane.** Each dot indicates a CHL molecule, and the y-axis shows its dwell time at the Ω1-H4 gate **(A)** or adjacent to the Ω1-H4 gate **(B)**. The definition of the criteria is shown in the Supplementary Materials. The dashed line marks the 30 ns dwell time threshold. The average dwell time of CHLs with dwell time >30 ns is listed below the figure.

**Supplementary Figure 11.**

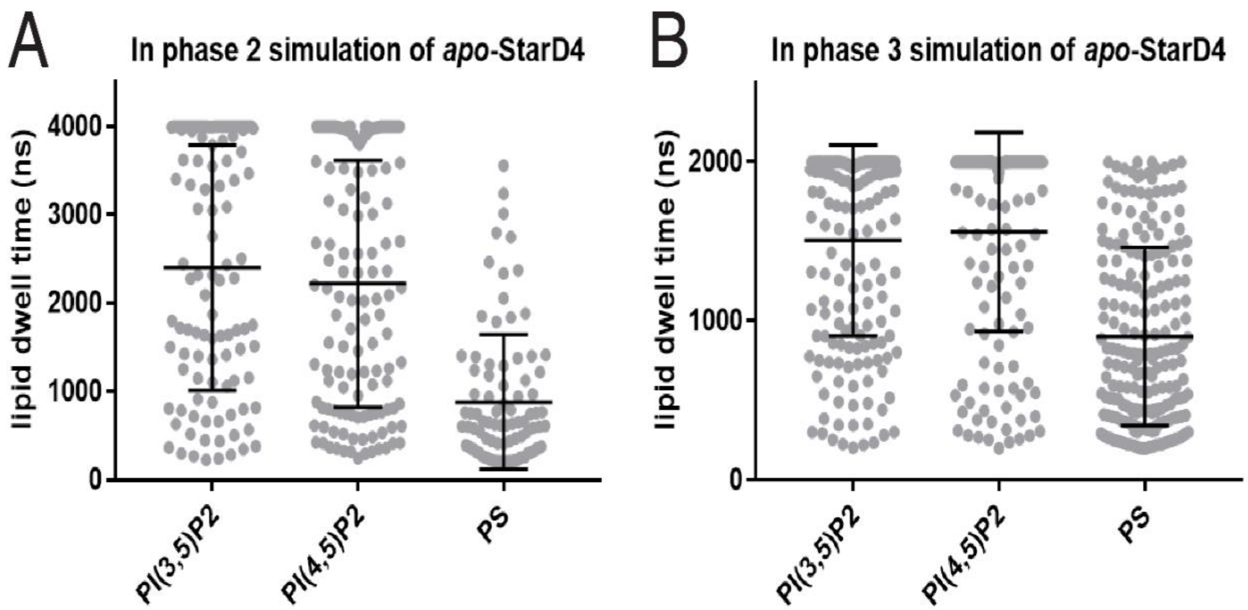


**Scatter plots of the dwell time of anionic lipids interacting with StarD4 in membranes containing PI(3,5)$P_2$, PI(4,5)$P_2$ or PS.** Each dot indicates an anionic lipid. An anionic lipid is considered to be interacting with StarD4 when any of its atom is with 4Å of the protein. Lipids with dwell time below 200 ns are excluded from the analysis. **(A)** Data obtained Phase 2 simulations of 4µs per replica; **(B)** Data from Phase 3 simulation of 2µs per replica. Statistical analyses show that the dwell time of PS is significantly shorter than the dwell time of either PI(3,5)$P_2$ or PI(4,5)$P_2$ with $p<0.0001$ by T test.

#### Supplementary Figure 12.

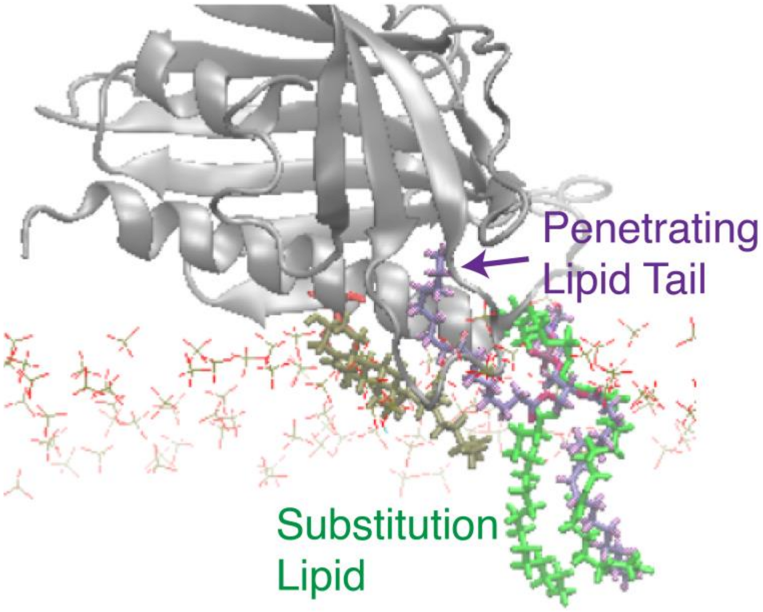


**Seed construction by manual substitution of the gate-penetrating membrane lipid tail.** To conduct the adaptive sampling of the extraction of membrane CHL by apo-StarD4, the initial seeds are chosen from the previous simulation with the membrane CHL positioned at the highest binding site, and an open H4-Ω1 gate. In the *apo*-StarD4, the opening of the H4-Ω1 gate is likely related to an inserted membrane lipid tail. In that case, we replaced the already inserted lipid with its own conformation it had before its insertion in the simulation in order to maintain the same interaction between the lipid and the protein in the expected insertion. The initial seeds bearing this manual modification are then equilibrated with the protein backbone and the target cholesterol constrained using the preparation protocol of StarD4-membrane complex as described.

**Supplementary Figure 13.**

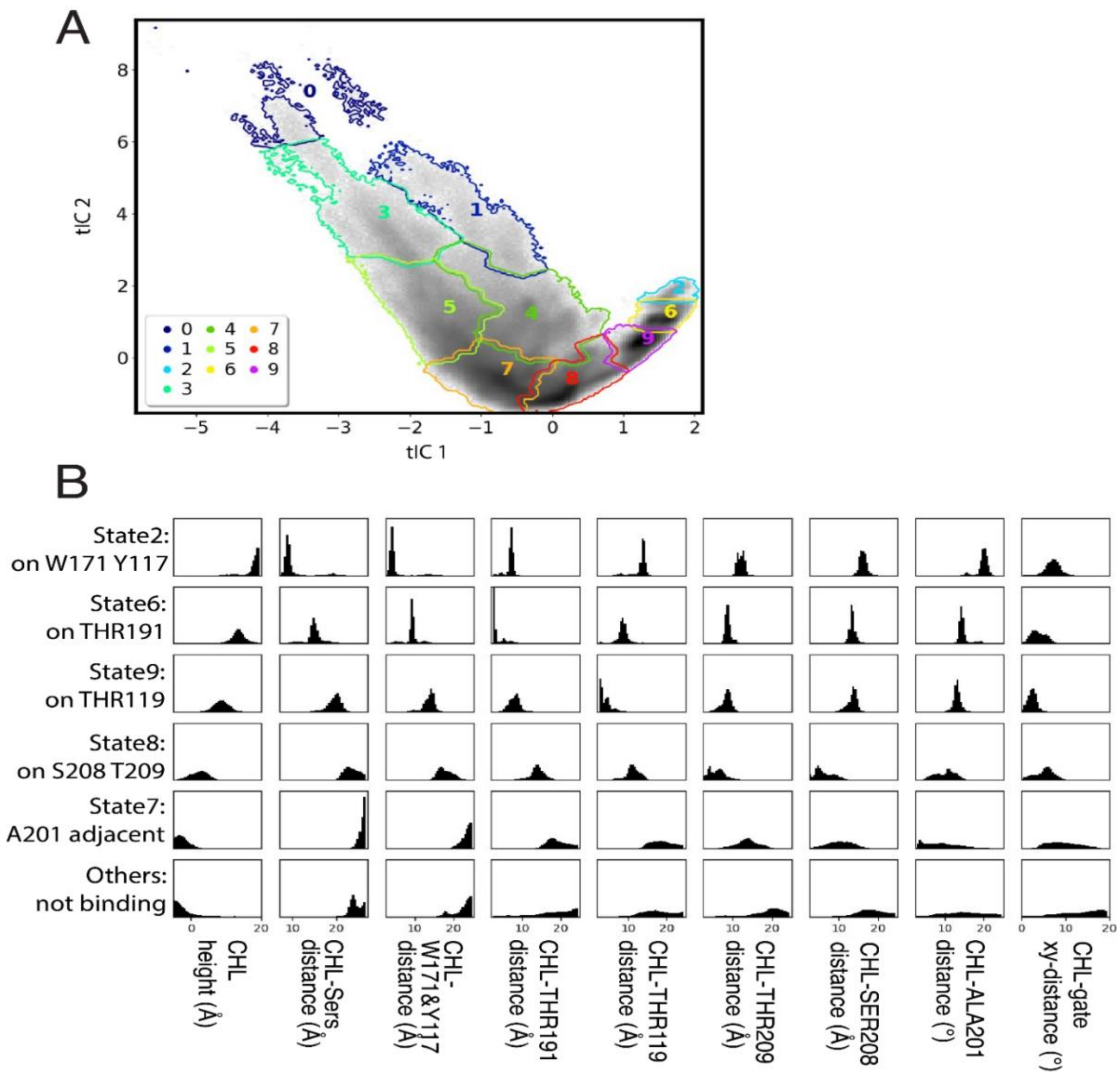


**The characteristics of metastable states along the cholesterol uptake pathway. (A)** 10 macrostates are outlined on the 2D tICA map defined in Fig. 18. **(B)** Structural characteristics of the macrostates along the cholesterol uptake pathway. The structural characteristics are represented by probability density histograms of the characteristic CV values that define the tICA space as in Fig. 18A. The states described (from top to bottom) are: macrostate 2 representing W171/Y117 binding, macrostate 6 representing T191 binding, macrostate 8 representing T119 binding, macrostate 7 representing S208/T209 binding. The other states are shown together.

#### Supplementary Figure 14.

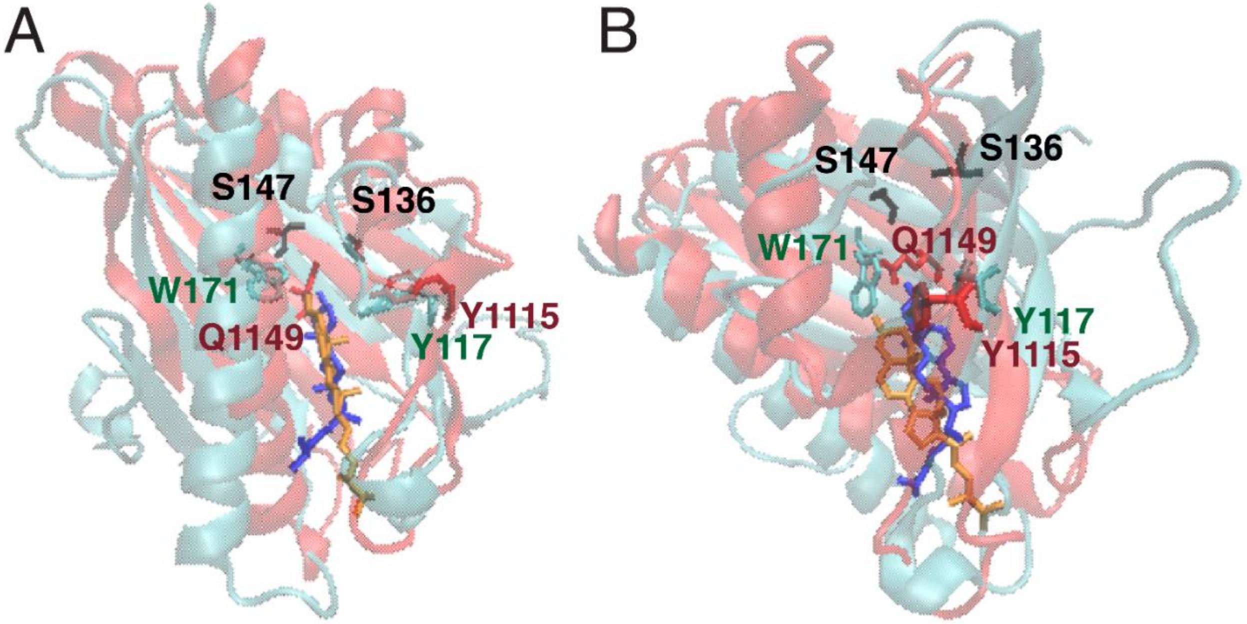


**Comparison of sterol-binding conformations inthe crystal structure of LAM2 and the *holo*-StarD4 complex in simulation. In both A and B** the representative conformation of the Trp-binding-CHL-StarD4 complex is rendered in cyan, with the W171 Y117 binding sites shown in cyan licorice, the S136 S147 in black licorice, and the cholesterol in blue licorice. The structure of LAM2 (PDB ID 5YS0) is rendered in red cartoon, with the sterol binding residues Q1149 Y1115 in red licorice, and its ligand sterol in orange licorice. Panel A: the front view; Panel B: the lateral view 90° from (A).

### SUPPLEMENTARY TABLES

**Supplementary Table 1: Structure-differentiating contact pairs in the transition event 2→3**

| Motif movements in this transition |
|---|
| **β1-H4 shifting**<br>**$PIP_2$ traverse binding**<br>**Hlp3-Ω1-H4 opening** |
| **Contact pairs FORMING in the transition (27 pairs):** |
| A24-S27, I26-V152, L34-C169, K49-R58, **K49-S200, K49-D203, V51-T204, K52-A211, K52-S215, V54-L210**, S61-N65, E63-N65, D76-R218, I83-T138, G89-L93, G89-L102, P90-L93, P90-D94, W91-G220, L98-T100, N125-P129, T138-L145, **PIP2-K52-R97, PIP2-K52-K219, PIP2-R97-S215, PIP2-S215, PIP2-S215-K219** |
| **Contact pairs BREAKING in the transition (14 pairs):** |
| L34-F172, **D53-R218**, V54-R218, D76-K223, I83-L146, R87-L145, W95-G220, **R97-A120, L98-Q122, L98-F213, Q122-T209, L123-A205,** PIP2-R97-T100, PIP2-T100 |

**Supplementary Table 2: Structure-differentiating contact pairs in the transition event 0→1**

| Motif movements in this transition |
|---|
| **β1-H4 shifting** |
| **Contact pairs FORMING in the transition (18 pairs):** |
| Y37-C169, **A48-D203, K49-T204, V51-S208**, V54-A71, S61-E63, N65-Q190, G66-Q190, D76-R218, D77-K223, V78-P180, N81-K223, S101-R158, PIP2-V47, PIP2-V47-K49, PIP2-V47-K50, PIP2-K219-R222, PIP2-R222 |
| **Contact pairs BREAKING in the transition (19 pairs):** |
| V47-K50, A48-Y69, **K49-S208, K49-A211, V56-A211**, P60-G66, Y67-L193, Y69-A207, D76-R222, R116-T118, I126-I197, T138-L145, L193-D203, K219-K223, G220-K223, PIP2-K49-V51, PIP2-K50-V51, PIP2-V51, PIP2-V51-K52 |

**Supplementary Table 3:** **Structure-differentiating contact pairs in the transition event 1→2**

| **Motif movements in this transition** |
|---|
| **β1-H4 shifting**<br>**B1&2-β3-break**<br>**β1-H4head PIP2 interaction** |
| **Contact pairs FORMING in the transition (25 pairs):** |
| L34-F172, **V51-A207, V54-R218**, D76-K223, I83-L146, R87-L145, M99-A120, I126-I197, **PIP2-R46-P198, PIP2-R46-Q199, PIP2-R46-S200, PIP2-R58-P198, PIP2-R58-Q199, PIP2-R58-S200,** PIP2-R97-T100, PIP2-R97-A120, PIP2-T100, PIP2-T100-A120, PIP2-A120, **PIP2-P198, PIP2-P198-Q199, PIP2-P198-S200, PIP2-Q199, PIP2-Q199-S200, PIP2-S200** |
| **Contact pairs BREAKING in the transition (14 pairs):** |
| A24-S27, L34-P168, L34-C169, **D53-G73, D53-V74, D76-R218**, D76-L221, I83-T138, G89-L102, W91-G220, L98-T100, PIP2-K49, PIP2-K49-K50, PIP2-K50 |

**<u>Supplementary Table 4.</u> The average number of PIP2s binding to the basic residue groups:**

Upper group: K52, R97, S215, R218, K219, R222

Left group: R46, K49, K50, K52, R58

Right group: R116, R130, R158, R163, R194

| | Average lipid count basic residue | $PI(3,5)P_2$ | $PI(4,5)P_2$ | PS |
|---|---|---|---|---|
| *holo*-StarD4 | Upper group | 2.24 | 2.09 | 0.66 |
| | Left | 3.53 | 3.32 | 1.08 |
| | Right | 1.63 | 1.74 | 0.35 |
| *apo*-StarD4 | Upper | 2.02 | 2.48 | 0.81 |
| | Left | 3.36 | 4.10 | 2.37 |
| | Right | 1.33 | 0.64 | 0.13 |

**<u>Supplementary Table 5.</u> Normalized coordination information (*NCI*) between non-correlated sites that serve as negative controls of allostery, and indicate the background level of correlation noise resulting from the complex dynamics of the system**

*Transmitter:*

*Receiver:*

| | β8β9 loop | β4-Nt | H4-Ct | CHL sites |
|---|---|---|---|---|
| **β8β9 loop** | 8.72 | 0.1% | 1.0% | 1.5% |
| **β4-Nt** | 0.3% | 4.50 | 2.8% | 7.5% |
| **H4-Ct** | 0.5% | 1.2% | 6.30 | 2.8% |
| **CHL sites** | 7.1% | 12.6% | 11.5% | 1.18 |

The *Transmitter* motifs are identified in the columns, *Receiver* motifs define the rows. The normalized coordination information (*NCI*) values were calculated with NbIT as described in the main text. The total correlation (*TC*) of each motif is shown in gray on the diagonal.

**<u>Supplementary Table 6.</u> Origin of and sample numbers of simulation data used in the construction of DNN**

<table>
<tr><th>Data source</th><th>PI(3,5)$P_2$</th><th>PI(4,5)$P_2$</th></tr>
<tr><td>Simulations in the membrane-embedding phase</td><td>63 trajectories,<br>11685 structures</td><td>69 trajectories,<br>12643 structures</td></tr>
<tr><td>Simulations in the membrane-embedded phase</td><td>24 trajectories,<br>12297 structures</td><td>24 trajectories,<br>9408 structures</td></tr>
<tr><td rowspan="2">Total sample</td><td>23982 structures</td><td>22051 structures</td></tr>
<tr><td colspan="2">46033 structures</td></tr>
</table>

$$F = F_{el} + F_{ion} + F_{lip}$$

The *electrostatic component* is determined by:

$$F_{el} = \frac{1}{2}\varepsilon_0\left(\frac{k_BT}{e^2}\right)\int_V \varepsilon_d(\nabla\Psi(\vec{r},t))^2 dv$$

where $\Psi(\vec{r},\mathrm{t}) = e\phi(\vec{r},\mathrm{t})/k_BT$ is the dimensionless electrostatic potential, with $\phi$ representing the electrostatic potential, and $e$ being the elementary charge, $\varepsilon_0$ being the vacuum permittivity, $\varepsilon_d$ being the dielectric constant ($\varepsilon_d = 80$ in aqueous solution and 2 within protein and membrane).

The *translational entropy* of mobile ions is determined by:

$$F_{ion} = \frac{1}{2}k_BT\int_V\left[n_+(\vec{r},t)\,ln\frac{n_+(\vec{r},t)}{n_0} + n_-(\vec{r},t)\,ln\frac{n_-(\vec{r},t)}{n_0} - (n_+(\vec{r},t) + n_-(\vec{r},t) - 2n_0)\right]dv$$

where $n_+(\vec{r},t)$ and $n_-(\vec{r},t)$ denotes the local concentrations of the positive and negative ions in the solution, and $n_0$ is the bulk concentration of the ions.

The *lipid mixing entropy* is determined by:

$$F_{lip} = \frac{k_BT}{a}\int_A\left(\phi(\vec{r},t)\,ln\frac{\phi(\vec{r},t)}{\phi_0} + \left(1-\phi(\vec{r},t)\right)ln\frac{(1-\phi(\vec{r},t))}{(1-\phi_0)}\right)ds$$

where $\phi(\vec{r},t)$ is the local mole fractions of the charged lipids in the membrane, and $\phi_0$ is the average composition of the charged lipid.

The minimization of this free energy function is carried out under the influence of the electrostatic field generated by the StarD4 protein, which yields a correspondingly optimized lipid rearrangement that is quantified from the local lipid density. The absorption energy of StarD4 into the membrane is determined by comparing the free energy of the StarD4-complex to the sum of the free energies of StarD4 and of the membrane considered separately. Here, the calculations were carried out in a $(256\ \text{Å})^3$ cubic space containing a 0.15M ionic solution of monovalent counterions, corresponding to λ=8.09Å Debye length, under the temperature of 310K.

The positioning of StarD4 is systematically explored in various orientations while ensuring that all atoms are at least 2Å away from the membrane surface. The orientation of StarD4A is defined with a spherical coordinate, with its origin located at the center of mass of StarD4, its z-axis parallel to the norm vector of H4, and its x-axis perpendicular and intersect with the norm vector of H4. The absorption energy of StarD4 is first examined with StarD4 facing the membrane from six orientations:

| | | StarD4 Orientation | | | | | |
|---|---|---|---|---|---|---|---|
| | | Up | Down | Front | Back | Left | Right |
| | | θ=0° φ=0° | θ=180° φ=0° | θ=90° φ=0° | θ=90° φ=180° | θ=90° φ=-90° | θ=90° φ=90° |
| Absorption energy ($k_BT$) | with 10% PIP2 | 1.61 | 0.83 | -3.78 | 0.92 | 3.66 | 3.53 |
| | with 10% PS | 0.62 | 0.28 | -1.07 | 0.58 | 1.79 | 1.64 |
